# Leaf spectroscopy reveals drought response variation in *Fagus sylvatica* saplings from across the species’ range

**DOI:** 10.1101/2024.07.23.604726

**Authors:** Dave Kurath, Sofia J. van Moorsel, Jolanda Klaver, Tis Voortman, Barbara Siegfried, Yves-Alain Brügger, Aboubakr Moradi, Ewa A. Czyż, Marylaure de La Harpe, Guido L. Wiesenberg, Michael E. Schaepman, Meredith C. Schuman

**Author notes:** Corresponding author: Sofia J. van Moorsel.

## Abstract

The common European beech (*F. sylvatica*), sensitive to prolonged drought, is expected to shift its distribution with climate change. To persist in novel environments, young trees rely on the capacity to express diverse response phenotypes. Several methods exist to study drought effects on trees and their diverse adaptive mechanisms, but these are usually destructive and challenging for the large sample numbers needed to investigate biological variation.

We conducted a common garden experiment outdoors, but under controlled watering conditions, with 180 potted two-year-old saplings from 16 beech provenances across the species’ range, representing three distinct genetic clusters. Drought stress was simulated by interrupting irrigation and stomatal conductance and soil moisture were used to assess drought severity. We measured leaf reflectance of visible to short-wave infrared electromagnetic radiation to determine drought-induced changes in biochemical and structural traits derived from spectral indices and a model of leaf optical properties.

We quantified changes in pigmentation, water balance, nitrogen, lignin, epicuticular wax, and leaf mass per area in drought-treated saplings, revealing differences in likely adaptive responses to drought. *Fagus sylvatica* saplings from the Iberian Peninsula showed signatures of greater drought resistance, i.e., the least drought-induced change in spectrally derived traits related to leaf pigments and leaf water content. We demonstrate that high-resolution leaf spectroscopy is an effective and non-destructive tool to assess individual drought responses that can characterize functional intraspecific variation among young beech trees. Next, this approach should be scaled up to canopy-level or airborne spectroscopy to support drought response assessments of forests.

**Plain language summary:** The common European beech tree, which is sensitive to prolonged droughts, is expected to experience local population declines due to climate change. To survive in a drier and warmer climate, young beech trees must show a variety of adaptive responses. Assessing this variation within the species is challenging, and traditional methods often harm the trees, limiting large-scale studies of their variability. We conducted an outdoor experiment with 180 potted young beech saplings from various European regions, simulating a severe drought by halting irrigation. Using advanced leaf reflectance measurements, we tracked biochemical and structural changes in the leaves, such as pigmentation, water content, and other traits. Our results highlight that beech saplings from the Iberian Peninsula demonstrated greater drought resistance, showing fewer changes compared to saplings from other regions. This study underscores the effectiveness of non-destructive, high-resolution leaf spectroscopy in assessing individual drought responses, revealing important insights into the adaptive capacity of beech trees under changing climatic conditions.

**Key Points:**

- Leaf reflectance measurements effectively track drought-induced trait changes in beech saplings in a non-destructive way.
- Beech saplings from the Iberian Peninsula show greater drought resistance with fewer biochemical and structural changes.
- High-resolution leaf spectroscopy reveals adaptive capacity differences within European beech populations under simulated drought stress.

## 1 Introduction

As a result of climate change, Europe faces prolonged drought periods that will threaten forest ecosystems (Cook et al., 2016; IPCC, 2023). The extended duration of local precipitation deficits will increase the risk of water stress and mortality for trees (C. D. Allen et al., 2010), and the series of recent record-breaking summers in northern Europe in 2003, 2018, and 2022 indicate what Europe’s forest ecosystems may experience in the near future (Milĺan, 2014; Hermann et al., 2023; Buras et al., 2020; Sturm et al., 2022). The common European beech (*Fagus sylvatica*) is a dominant and widespread tree species in Europe (Durrant et al., 2016), but its dominance comes at the cost of considerable drought sensitivity compared to co-occurring broadleaf tree species (Packham et al., 2012; Leuschner, 2020). Consequently, beech faces a disproportionate risk of habitat loss and growth decline, and may ultimately change its natural distribution due to reduced competitiveness (Martinez del Castillo et al., 2022; Leuschner, 2020; Rigling et al., 2019; Dittmar et al., 2003; Schmied et al., 2023; Rose et al., 2009; Kreyling et al., 2014; Aranda et al., 2015; Geßler et al., 2006).

To establish and persist, young trees rely on phenotypic variation, i.e., the potential to express diverse phenotypes given their genetic background and environmental conditions (Arnold et al., 2019; Benito Garźon et al., 2019; Whitman & Ananthakrishnan, 2009). Intra- and interspecific diversity are expected to increase the probability of survival for young trees in communities experiencing novel environmental conditions, such as more intense or prolonged drought (Blondeel et al., 2024; Gazol et al., 2023). Intraspecific variation encompasses a range of physiological and morphological traits providing options for individuals to acclimate, and on which selection can operate, so species can adapt. Different individuals within a species may vary in their drought response, making some more likely to survive (Nguyen et al., 2017). Because drought responses are complex, involving orchestrated changes in physiology, differences among trees in individual traits may not predict differences in drought responses, and multiple strategies are possible (Martínez-Vilalta et al., 2023). Thus, greater within-species variation can serve as a buffer against environmental stress, ensuring that at least some members of a species or population can withstand prolonged periods of drought. The spatial distribution of this variation within versus between populations may determine which, and how many populations can acclimate and adapt: those differences that are inherited across generations provide material for adaptation.

Acclimation of trees to drought is apparent through several physiological changes in leaf traits (Pflug et al., 2018; Leuzinger et al., 2005; Tognetti et al., 1995; Rukh et al., 2023). As an immediate response, for example, plants can close their stomata to prevent hydraulic failure (McDowell et al., 2008). An anisohydric species (Leuschner, 2020), beech shows high stomatal sensitivity to water deficit and adapts stomatal conductance to optimize photosynthesis in water-scarce situations (Peuke et al., 2002). Photosynthetic pigments such as chlorophyll *a* and *b* (*C_ab_*) are essential for oxygenic photosynthesis in plants, and water deficit and accompanying oxidative stress can reduce chlorophyll content, corresponding to decreased photosynthetic capacity (Wu et al., 2016; F. Hu et al., 2023; Mafakheri et al., 2010; Gai et al., 2023). Carotenoids (*C_car_*) – xanthophyll cycle pigments and carotenes – serve as important photoprotective pigments by absorbing excess solar energy in the blue to green wavelengths, and protect cells from oxidative stress by reacting with singlet oxygen and free radicals, although regeneration e.g. via ascorbic acid is necessary to prevent damage due to oxidized carotenoid products (T. Sun et al., 2022; Havaux, 2014; Ben Abdallah et al., 2017; Baccari et al., 2020; Bacelar et al., 2007; Edge & Truscott, 2018). Thus, variation in the chlorophyll to carotenoid ratio *C_ab_/C_car_* may be a good indicator for drought stress in plants (Green & Durnford, 1996). Anthocyanins (*C_ant_*), which are generally responsible for the red colouration of senescent leaves, also act as antioxidants and absorb light in the UV, blue and green ranges (Gould, 2004; Saha et al., 2020). Drought stress increases oxidative stress, which can lead to anthocyanin accumulation (Z. Li & Ahammed, 2023). Long-term responses include photoprotective mechanisms, such as an increased ratio of xanthophyll cycle pigments (VAZ) to chlorophylls. Drought-stressed plants also show reversible downregulation of photosystem II (PSII), which results in a decrease of quantum yield and photosynthetic efficiency, but also reduces the generation of free radicals from photosynthesis (Galĺe & Feller, 2007; Garćıa-Plazaola & Becerril, 2000). Equivalent water thickness (EWT), which is a direct physical representation of water within the leaves, is associated with leaf-level drought tolerance (Féret et al., 2019; Junttila et al., 2022). Leaf water potential (Ψ) includes all energy gradients that move water such as turgor, osmotic and matrix potential and indicates increased stress levels and downregulation of physiological processes with progressing drought (Walthert et al., 2021). The biosynthesis of lignin, which is found in the cell walls of plants and is responsible for the overall stability of plant cells, is up-regulated during water deficit (Y. Hu et al., 2009; Liu et al., 2018; Brinkmann et al., 2003). Leaf epicuticular waxes restrict diffusion across the cuticle and reduce oxidative stress by absorbing UV light, and their composition may change in response to environmental factors (Speckert et al., 2023; Jenks & Ashworth, 1998). Constitutive differences in leaf traits can also be related to drought responses. Leaf mass per area (LMA) is usually calculated with leaf dry mass and is determined by many bulk constituents of leaves including starch, sugars, and proteins. LMA is often related to abiotic stress responses such as drought (Wright et al., 2004; Wellstein et al., 2017; Quero et al., 2006; De La Riva et al., 2016), but a sudden withholding of water does not cause LMA variation (Poorter et al., 2009).

Structural and biochemical profiling commonly require destructive sampling and laboratory analysis (Pflug et al., 2018; Jacquemoud et al., 1996). These are usually precise, but difficult to scale due to time and costs, and the requirement to work with harvested leaf tissue can introduce artefacts (Burnett, Serbin, et al., 2021).

High-resolution leaf spectroscopy, a contact measurement closely related to field and imaging spectroscopy, offers a relatively cost-effective, non-destructive, and scalable solution for retrieving leaf traits directly from living leaves on plants (Jacquemoud & Baret, 1990; Féret et al., 2019). Spectroscopy of individual leaves is commonly conducted using a contact probe with a standardized light source and backgrounds.

It measures aspects of leaf optical properties (LOP), which describe how leaves physically interact with electromagnetic radiation. Since the biochemical and structural composition of leaves determine how they reflect, transmit, absorb, or fluoresce light, variation in LOP thus carries information about leaf traits (Jacquemoud & Ustin, 2019). Recent studies highlighted that spectral indices may not entirely reflect leaf structural and biochemical properties (Petibon et al., 2023), however, a growing body of literature links spectra to plant trait variation (e.g. (D’Odorico et al., 2023; Meireles et al., 2020; Asner & Martin, 2011; Z. Wang et al., 2022)).

Leaf traits are estimated from spectra based on empirical or physical approaches (Féret et al., 2017). Traits are derived empirically using machine learning methods such as partial least squares regression (PLSR), which require training data and yield somewhat, although not exclusively, dataset-specific models (Burnett, Anderson, et al., 2021; Féret et al., 2019; Ji et al., 2024). A simple empirical-based method, spectral indices are validated using laboratory measurements but do not require training data; these comprise reflectance values at different bands and are developed from two or more wavelengths in which the target of interest causes a relative absorption difference (Jacquemoud & Ustin, 2019; Verrelst et al., 2015).

Indices have been found useful to estimate the relative values of traits such as photosynthetic activity (J. A. Gamon et al., 2016), and are computationally very efficient (Jacquemoud & Ustin, 2019). A physical approach, radiative transfer models (RTMs) such as the extensively used open-source model propríetés spectrales (PROSPECT) (Jacquemoud & Baret, 1990), rely on the physical properties of electromagnetic radiation and its interaction with matter such as absorption and scattering coefficients, and are thus theoretically robust (Feret et al., 2008). In PROSPECT, broadleaves are represented as a generalized plate model with uniform layers (specifically air – cell wall interfaces) quantified as a structure parameter *N*, characterizing the leaf mesophyll structure (W. A. Allen et al., 1969; Stokes, 1860; Spafford et al., 2021; Jacquemoud & Baret, 1990), with which directional hemispherical reflectance and transmittance between 400 nm and 2500 nm are simulated (Schaepman-Strub et al., 2006). One of the most recent PROSPECT developments is called PROSPECT-D(-ynamic). It allows *C_ab_*, *C_car_*, *C_ant_*, *N*, EWT, and LMA as leaf trait parameter inputs to simulate leaf spectral profiles, and outperforms previous versions (Féret et al., 2017; Feret et al., 2008). The inversion of PROSPECT-D enables the retrieval of leaf traits based on provided reflectance, or transmittance data. The inversion makes use of optimization algorithms with a corresponding merit function (Jacquemoud et al., 1996; Tarantola, 2005). PROSPECT RTMs have been successfully applied across remote sensing and plant ecology research (Serbin et al., 2014; Zhao et al., 2014; Asner et al., 2014; J. Sun et al., 2018; Kothari & Schweiger, 2022).

To study intraspecific variation in the drought responses of young *F. sylvatica* trees, we conducted a common garden experiment on 180 two-year-old saplings originating from 16 different provenances across the species’ range. We determined their population genetic structure, identifying three main genetic clusters. We simulated a drought scenario by restricting water for 14 days in early summer. We measured leaf reflectance with a field spectroradiometer at the end of the drought period to quantify spectral variation between the control (continued watering) and treatment groups, and to derive leaf traits using both spectral indices and an RTM approach (PROSPECT-D inversion). We also measured soil moisture, leaf transpiration, and leaf water content as indicators of drought effects, as well as leaf mass per area and pigmentation, for comparison to spectroscopy measurements. We hypothesized that (i) drought-treated saplings show different responses in spectral indices and simulated leaf traits compared to the control group that can be interpreted in physiological terms, and (ii) there is intraspecific variation in drought responses dependent on the sapling’s genetic background.

## 2 Materials and Methods

### 2.1 Seed collection, germination, and growth conditions

Seeds were collected in 2020 between August and October from well-protected sites across Europe, thus having a relatively high probability of harbouring populations representing natural migration more than introductions for forestry (see supplementary methods). European beech was the dominant tree species at all sites. The majority of seed collections were sampled directly from branches of trees located in an area of 10 x 200 meters. However, we also conducted bulk collections from the ground not more than 1 km from a core area of trees used for leaf spectroscopy measurements in Czyż et al. (2023). Seeds were placed in labelled containers per collection, and upon arrival, dried in mesh envelopes (1/collection) in closed plastic boxes under a gentle nitrogen flow. Once dried, the seeds were transferred to paper envelopes and preserved in sealed plastic bags at 5° C until February 2021 (for more details, see supplementary methods).

Seeds of 94 collections total from 24 sites with varying mean annual precipitation and temperature (see Table 1, Table S1). The sites spanned multiple major climate zones in Europe, from subtropical to temperate climates: temperate oceanic climate, hot-summer Mediterranean climate and warm-summer humid continental climate. The seeds were brought to a nursery on 25 February 2020 (47°35’48.9”N, 9°11’40.5”E, Kressibucher AG, Berg Thurgau, Switzerland), where they were imbibed in water for 24 hours, and then sown in 5 x 5 x 20 cm pots in flats (max 4 seeds/pot) with mixed humus and wood chips and no additional fertilizers. A wire grid was installed over the pots and the flats were put outside and exposed to the environment. Damage to the dicotyledons was mitigated by bringing the flats inside a closed shelter when temperatures fell below freezing. Seedlings germinated by July 2021, with seedlings emerging for 43 of the 94 collections, from 16 of the 24 sites, and an overall seedling emergence rate of 3.5% (7.9% for the 43 collections in which seedlings emerged). A summary of the resulting provenances is provided in Table 1.

**Table 1.** Summary of *F. sylvatica* provenances including sample sizes *n*, elevation, coordinates, date of collection, mean temperature (*T_mean_*) and mean annual precipitation (MAP). Both MAP and *T_mean_* were downloaded from WorldClim (Hijmans et al., 2005). The Koeppen-Geiger climate zone was taken from R package *kgc* (v1.0.0.2) (Bryant et al., 2017). *Cfb* temperate oceanic climate, *Csa* Hot-summer Mediterranean climate, *Dfb* Warm-summer humid continental climate. Seeds started to germinate in the last week of February 2021.

| Label | Country | Locality | n | Lat./Lon. | Elevation<br>(m.a.s.l.) | $T_{mean}$<br>(°C) | MAP<br>(mm) | KG<br>zone | Collection<br>date |
| --- | --- | --- | --- | --- | --- | --- | --- | --- | --- |
| BES | Belgium | Sonian Forest | 11 | 50.751/4.423 | 168 | 10.16 | 805 | Cfb | 02.09.2020 |
| CHL | Switzerland | Laegern | 8 | 47.479/8.364 | 733 | 9.21 | 1137 | Cfb | 07.08.2020 |
| ESM | Spain | Moncayo Natural Park | 11 | 41.791/-1.810 | 1479 | 9.88 | 556 | Cfb | 23.09.2020 |
| ESP | Spain | San Juan de la Pena | 31 | 42.508/-0.676 | 1200 | 10.88 | 720 | Cfb | 24.09.2020 |
| FRB | France | Sainte Baume | 24 | 43.328/5.769 | 950 | 13.56 | 715 | Csa | 19.09.2020 |
| FRM | France | Massane | 6 | 42.491/3.032 | 722 | 14.20 | 604 | Csa | 25.08.2020 |
| HBV | Bosnia | Vlasenica | 9 | 44.168/18.919 | 1050 | 10.60 | 1216 | Cfb | 2020 |
| HRP | Croatia | Paklenica National Park | 18 | 44.360/15.467 | 924 | 7.82 | 1267 | Cfb | 16.09.2020 |
| ITB | Italy | Bracciano Nature Reserve | 2 | 42.174/12.156 | 584 | 13.70 | 234 | Csa | 21.08.2020 |
| ITE | Italy | Etna | 2 | 37.709/15.045 | 1721 | 11.41 | 699 | Csa | 17.08.2020 |
| ITM | Italy | Madonie Natural Park | 2 | 37.858/14.042 | 1711 | 13.46 | 567 | Csa | 16.08.2020 |
| PLB | Poland | Bieszczadzki Natural Park | 2 | 49.232/22.472 | 893 | 6.01 | 933 | Dfb | 11.09.2020 |
| PLW | Poland | Bukowa Mountain | 19 | 49.669/18.860 | 662 | 7.19 | 925 | Dfb | 09.09.2020 |
| ROS | Romania | Șinca | 8 | 45.654/25.164 | 865 | 9.80 | 1182 | Dfb | 13.09.2020 |
| RSF | Serbia | Fruška Gora National Park | 9 | 45.138/19.636 | 474 | 11.09 | 645 | Cfb | 17.09.2020 |
| SLK | Slovenia | Krokar forest | 18 | 45.543/14.763 | 1178 | 7.71 | 1464 | Cfb | 17.09.2020 |
| Total |  |  | 180 |  |  |  |  |  |  |

In November 2021, saplings were transported to the Hauenstein garden center in Rafz, Zurich, Switzerland (47°36’36”N, 8°32’35”E). The plants were transplanted into tall 4 L pots filled with an equally proportioned mixture of peat-free potting soil and low-organic matter sandy soil. The potting soil was enriched with Osmocote Exact Standard NPK (Mg) 12-14M (ICL Group, Tel Aviv-Jaffa, Israel) with micro-nutrients as fertilizer, containing 15% total nitrogen (N), 8% phosphorus pentoxide (*P*_2_*O*_5_), 11% potassium oxide (*K*_2_*O*) and 2% total magnesium oxide (MgO). During the winter, the saplings were in a winter tunnel, where temperatures were maintained at circa 7° C. The saplings were watered by an overhead irrigation system as needed. Furthermore, Multikraft BB Start (Multikraft, Pichl/Wels, Austria) was used to treat the soil with ectomycorrhizal fungi. Once it was determined that the saplings were no longer at risk of frost damage, the pots were moved outside the winter tunnel (12 May 2022). We controlled aphid infestation by repeatedly spraying NeemAzal-T/S (3ml/L in water, Andermatt Biogarten, Grossdietwil, Switzerland) directly onto the leaves.

### 2.2 Common garden experiment

We conducted our experiment with 180 2-year-old saplings from 16 provenances across the natural distribution range in June 2023. The saplings were transported from the Hauenstein nursery to the experiment site at the University of Zurich, Zurich, Switzerland (47°23’44”N, 8°33’05”E; elevation 509 m.a.s.l.). We implemented a 2 x 5 randomized block arrangement, in which each block formed a 3 x 6 grid (Fig. S1). Each block contained 18 saplings, which were placed in the slots of a steel grid, which were spaced approximately 1 m apart to minimize interactions between saplings of different blocks. Pots were placed on black polypropylene fabric for weed control, and secured to the steel grid with hemp cord (Fig. 1b). We installed an all-in-one weather station (ATMOS 41, METER Group, Pullman, WA United States) to measure local weather conditions such as precipitation, humidity and air temperature.

**Figure 1.**
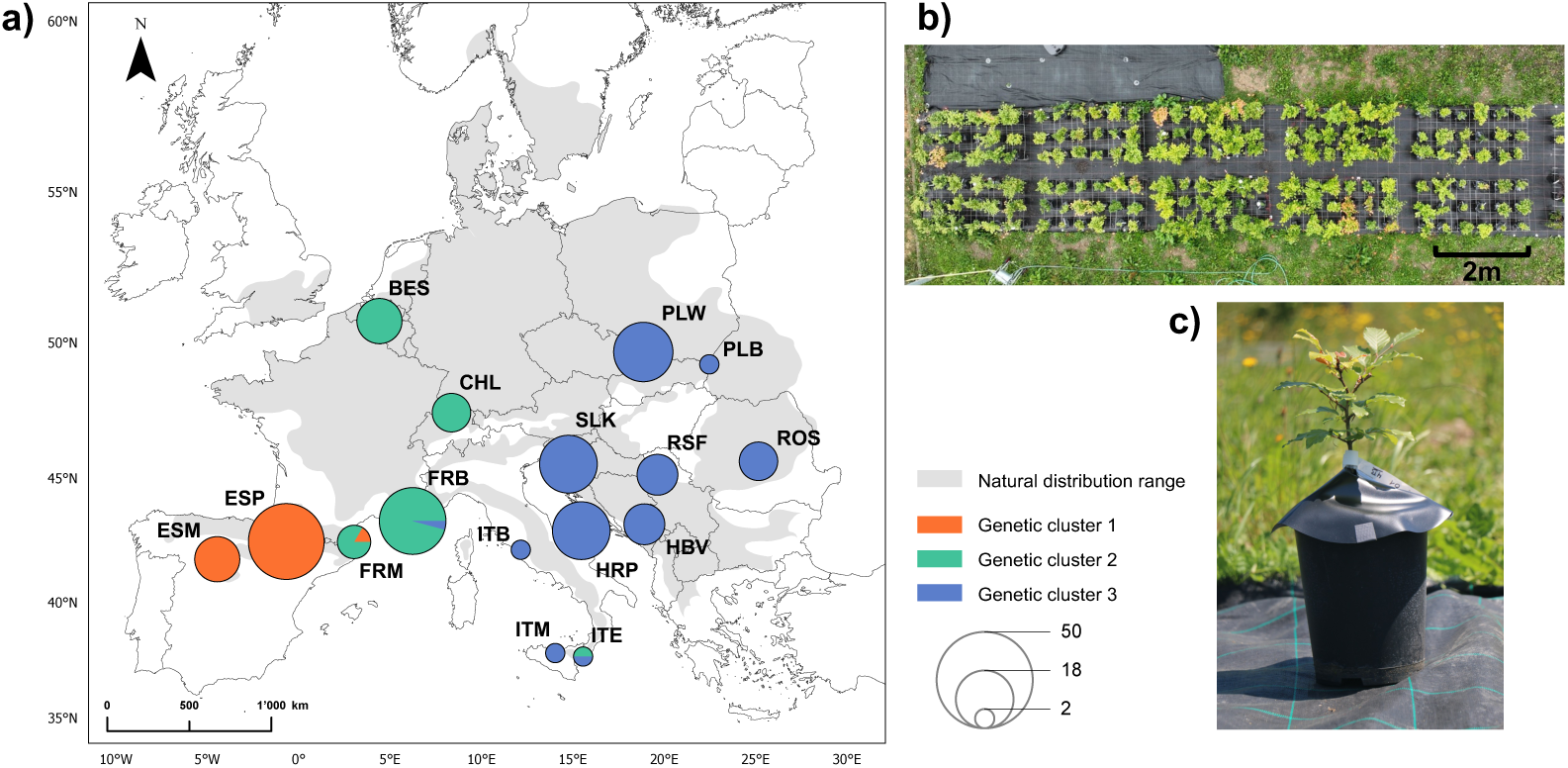
a) European map depicting sampling locations and natural distribution range of *F. sylvatica* (EUFORGEN) (Caudullo et al., 2017). The first two letters in the abbreviations are country identifiers, and the third letter is the site identifier. The provenances are shown as pie charts with the proportions of the samples belonging to the three genetic clusters. The map was created in ArcGIS Pro (v3.1) (ESRI Inc., 2023). b) UAV footage of the experiment site in June 2023. c) Rain cover over the pots used to simulate the drought scenario.

We maximized the diversity of provenances within the blocks, so each provenance was represented in every block whenever possible (Fig. S1a). For each provenance, half of the saplings were assigned to the drought treatment and half to the control (for details, see Fig. S1a). Seed collection was not considered, due to both the limited and unequal number of seed collections per provenance and the limited and unequal number of individuals per collection (see Table S2). The placement of the saplings within the blocks was randomized. In April 2023, we measured the length of the primary shoot using a flexible tape measure from the root collar along the shoot to the highest point in its natural position. We then used these plant height measurements to select individuals to have a similar distribution of heights among the blocks (Fig. S1b).

### 2.3 Drought treatment

We selected a drought duration that would avoid tree mortality in the experiment while inducing realistic drought stress. Currently, Switzerland encounters an average of 11 consecutive rain-free days each year, and it is anticipated that these dry periods may extend to as many as 19 days in the midterm future (NCCS, 2018). Therefore, the 14-day experimental drought is a scenario that beech saplings are projected to experience in Switzerland in the shorter-term future.

During the simulated drought period, both groups were subjected to a controlled exclusion from rain using cone-shaped rain covers which enclosed the entire pot (Fig. 1c), but the control group was watered regularly according to demand and at least every other day. Covers were made of waterproof and UV-resistant PVC black pond foil (Heissner GmbH, Lauterbach, Hessen, Germany) and fixed on the sapling stem with Parafilm (Pechiney Plastic Packaging Inc., Chicago, IL, United States) beneath the first stem node. Three thin wooden sticks were used to elevate the cover to leave a gap between the foil and the pot to enable air circulation. In addition, we inverted the pot trays in the treatment group to avoid the retention of water in the soil.

We prioritized sapling survival by *ad hoc* irrigation of the drought-treated saplings: when severe wilting was observed, we watered wilting saplings with 600 mL water (see Table S3 for irrigation timeline and details). Wilting was monitored daily and quantified by a wilting index. The wilting index, ranging from 0 to 4, was categorized into quartiles. A score of 0 denoted the absence of wilting, while scores 1-4 indicated wilting levels corresponding to ≤ 25%, *>* 25% to ≤ 50%, *>* 50% to ≤ 75%, and *>* 75% of leaves per saplings, respectively. Our aim was to avoid score 4, as such severe wilting would likely lead to mortality.

To monitor soil moisture and air temperature, we installed 20 TMS-4 probes (TOMST, Prag, Czech Republic) (Wild et al., 2019) in pots from block A to F. The probes covered as many provenances as possible while capturing the full range of plant heights within a single block. Soil moisture and soil temperature were sampled in an interval of 15 minutes during the entire experiment (Fig. S2).

### 2.4 Leaf spectroscopy

Leaf spectroscopy measurements were taken during the second half of the drought (days 9 to 14, corresponding to June 19^th^ to 24^th^, 2023). We constrained the measurement time to ±3 hours from solar noon. Therefore, within one day, we measured two blocks (one control, one treatment block) totalling 36 individuals and 72 leaves (2/individual), except for one day when 72 individuals corresponding to 144 leaves were measured. We used the ASD FieldSpec 4 Standard-Res spectrometer (ASD Inc., Boulder, USA), which measures electromagnetic radiation across a total of 2151 bands covering the spectral range from 350 nm to 2500 nm. We used a plant probe with a leaf clip (model A122317, serial N° 455, ASD Inc., Boulder, USA) with a calibrated low-intensity halogen light source. The leaf clip allows the non-destructive handling of the leaves and the isolation of external illumination. We used the spectral acquisition software RS^3^ (ASD Inc., Boulder, CO, USA) with respective instrument configurations (Table S4). Before taking the measurements, the spectrometer was warmed up for at least 30 minutes to decrease measurement errors due to variation in ambient and internal temperatures (Hueni & Bialek, 2017). For each sapling, we measured two random sun-exposed top-of-canopy leaves. We placed the leaf clip on the leaf adaxial surface, filling the measurement window of the plant probe and avoiding the midrib. Spectral measurements consisted of four successive measurements for each leaf: the white reference (*R_w_*), the white reference with target leaf (*T_w_*), the black reference (*R_b_*), and the black background reference with target leaf (*T_b_*). We gathered 5 readings per measurement, totalling 20 measurements per leaf. We recalibrated (optimized) the sensor gain with the RS^3^ software against the white reference background of the leaf clip after measuring 10 saplings to correct for differences in temperature sensitivity of the three detectors resulting in deviations of the white reference reflectance value from one (Hueni & Bialek, 2017). The leaf clip and plant probe were regularly checked for dirt or other contamination and cleaned, or white and black reference stickers were replaced as needed.

### 2.5 Stomatal conductance

During the drought, we measured stomatal conductance (*g_s_*) of a subset of the saplings (n=141) with an SC-1 leaf porometer (METER Group, Pullman, WA United States) according to the manufacturer’s protocol. Along with the spectral measurements, we constrained the measurement time to ±3 hours from solar noon due to high variation during the day (Leuschner, 2020). Given this constraint and the significant time required for each measurement, it was not feasible to measure all the beech saplings in a single day. Therefore, we measured two to three blocks each day. To standardize the strong and immediate effects of changing conditions on stomatal conductance during the 6-hour measurement window, we always measured a sapling from the control directly after measuring one from the treatment. With this ”paired” approach, the solar radiation and air temperature were on average similar for treatment and control, which allows comparison between them. We calibrated the device according to the manufacturer’s manual on each measurement day. When measuring, we took the values as soon as they stabilized. Per sapling, we measured two representative sun-exposed leaves on the adaxial side, where the stomata of beech are found (Peat & Fitter, 1994).

### 2.6 Validation of modelled pigments and structural traits

To establish a validation dataset for the modelled pigments and structural and biochemical traits (*C_ab_*, *C_car_*, EWT and LMA), we collected leaves from across the control and treatment group. For pigment extraction, immediately after their spectral measurement, the leaf samples (n=56) were stored in black plastic bags on dry ice to prevent degradation of the pigments and within an hour moved to – 80° C. To measure leaf area and fresh weight, we used a second leaf from the same sapling. To get the fresh weight, the leaf samples (n=56) were weighed immediately after their spectral measurement using a portable field scale. Leaf area was calculated using the image analysis program ImageJ (v1.38) (Schneider et al., 2012) based on a photo taken above parallel to the leaf with a size standard (see Fig. S3 for an example).

Dry weight was assessed after 72 h at 70° C using a semi-micro balance (MSU125P, Sartorius, Göttingen, Germany, 0.015 mg repeatability). We then calculated EWT and LMA based on the equations by Feret (2019, see also supplementary methods for more details (Féret et al., 2019)).

For a subset of the collected samples (n=20), the pigment extraction and subsequent HPLC analysis was done according to the protocol proposed by Petibon and Wiesenberg (2022) and conducted under subdued light to prevent pigment degradation. We modified the gradient program and the eluent composition of the HPLC method to optimize peak separation (see supplementary methods). We identified and quantified seven major pigments (chlorophyll a and b, *α*− and *β*-carotene, lutein, neoxanthin, and zeaxanthin) based on the measurement of standard solutions containing the respective analyte (see supplementary methods for the standards’ sources). From the results we calculated the parameters *C_ab_* (sum of chlorophyll a and b) and *C_car_* (sum of a/b-carotene, lutein, neoxanthin, zeaxanthin) in *mg.cm*^2^ as measure for the chlorophyll and carotenoid concentration. We observed co-elutionof *α*and *β*-carotene, for which we used the *α*-carotene standard calibration. Additionally, we assigned all remaining peaks to one of the following compound classes: pheophytin, chlorophyll-derivatives, carotene-derivatives or unknown compound, based on their absorption spectra and the decision tree proposed by Petibon and Wiesenberg (2022). For these peaks, no quantification was carried out due to the lack of standard material. Since the method was optimized to retrieve chlorophylls and carotenoids, we were unable to assess anthocyanin concentrations (Figure S4a, b); however, measurements included one purchased copper beech (a variety with purple leaves due to a mutation that inhibits anthocyanin breakdown (Packham et al., 2012)) as an extreme phenotype for comparison, and anthocyanins were detected in this sample (Figure S4c), but not in any other samples.

### 2.7 DNA extraction and whole-genome sequencing

We sampled leaf tissue (one to two mature leaves, i.e. roughly 0.1 g per plant) for genetic analysis by wrapping the leaves in aluminium foil and freezing them on dry ice in June 2022. The samples were stored at -80° C until further processing.

Prior to grinding, samples were flash-frozen in liquid nitrogen. Grinding was performed using 3 mm glass beads at 30 Hz for three min in a TissueLyser II (Qiagen). Plant DNA was then extracted using the Norgen Plant and Fungi DNA extraction kit (Norgen Biotek, Thorold, Ontario, Canada), and following the manufacturer’s protocol with some optimization steps added to minimize RNA contamination and increase lysis, yield, and purity (described in detail in Czyż et al. (2023)). The quality and quantity of the purified DNA was assessed with Qubit dsDNA HS Assay Kits (Thermo Fisher Scientific), a DS-11+ microvolume spectrophotometer (DeNovix), and agarose gel electrophoresis. PCR-based library preparation and Illumina paired-end whole-genome sequencing (n=180 samples) were conducted by Novogene (UK). We obtained 11 GB of raw data per sample, resulting in a mean 20x coverage of the 541 Mbp nuclear reference genome (Mishra et al., 2022).

### 2.8 Data analyses

#### 2.8.1 Spectral data

To process the spectral data, we used R (v4.3.1) (R Core Team, 2023) with the *spectrolab* package (v0.0.18) (Meireles & Schweiger, 2020). We spliced the bands at sensor transitions, and interpolated them with the *match sensor()* function. Due to the low signal:noise ratio and higher absolute uncertainty in the 350-400 nm region, we excluded that range (Petibon et al., 2021).

The measurement per sample consisted of five replicates each of a reference measurement with both black (*R_b_*) and white backgrounds (*R_w_*) and target leaf with both black (*T_b_*) and white backgrounds (*T_w_*) (see section 2.4). To calculate the total reflectance of each leaf, we took the mean of the five scans for each measurement. The mean reflectance values were then used to calculate the leaf sample reflectance *R_l_* based on the equation by Miller et al. (1987):

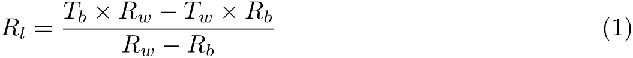

The value is the scaled percentage of reflected radiation, where 1 indicates 100% reflection and 0 means 0% reflection. We derived information about leaf traits by calculating indices with *R_l_*. Fig. 2a shows the mean spectral reflectance across control and treatment group. Fig. 1b-d show individual reflectances depending on their genetic clusters. We calculated previously published spectral indices (Table 2) that are indicative for leaf biochemical traits and structural traits.

**Figure 2.**
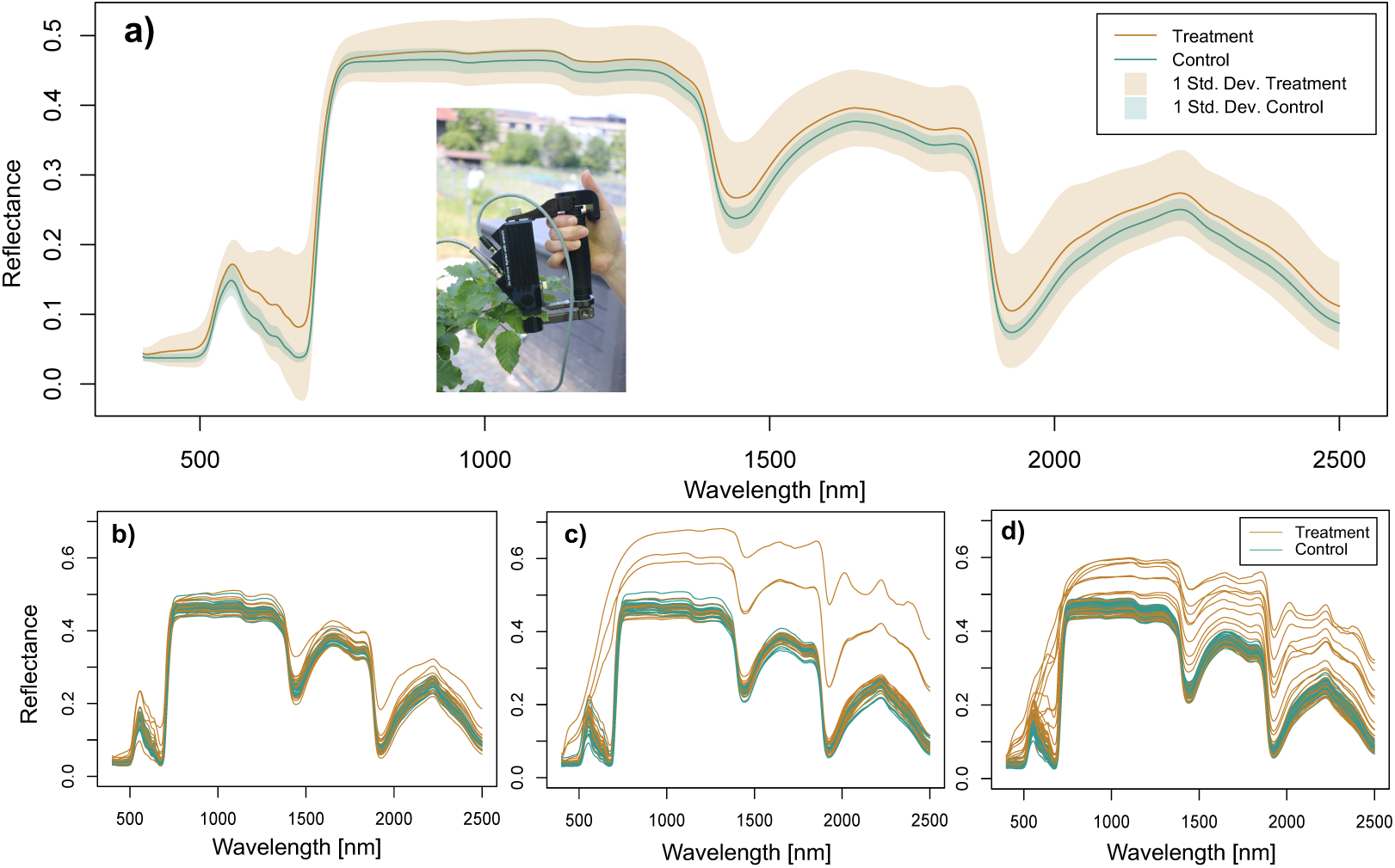
a) Mean reflectance in the spectral range from 400-2500 nm of the leaves from the treatment (orange lines, n=90) and control (green lines, n=90). Shown are calculated reflectance spectra following splice correction (see section 2.8.1). The inset shows the leaf clip used for the spectral measurements. b) Sample reflectances for saplings belonging to genetic cluster 1 (n=43). c) Sample reflectances for saplings from genetic cluster 2 (n=48). d) Sample reflectances for saplings belonging to genetic cluster 3 (n=89).

**Table 2.**
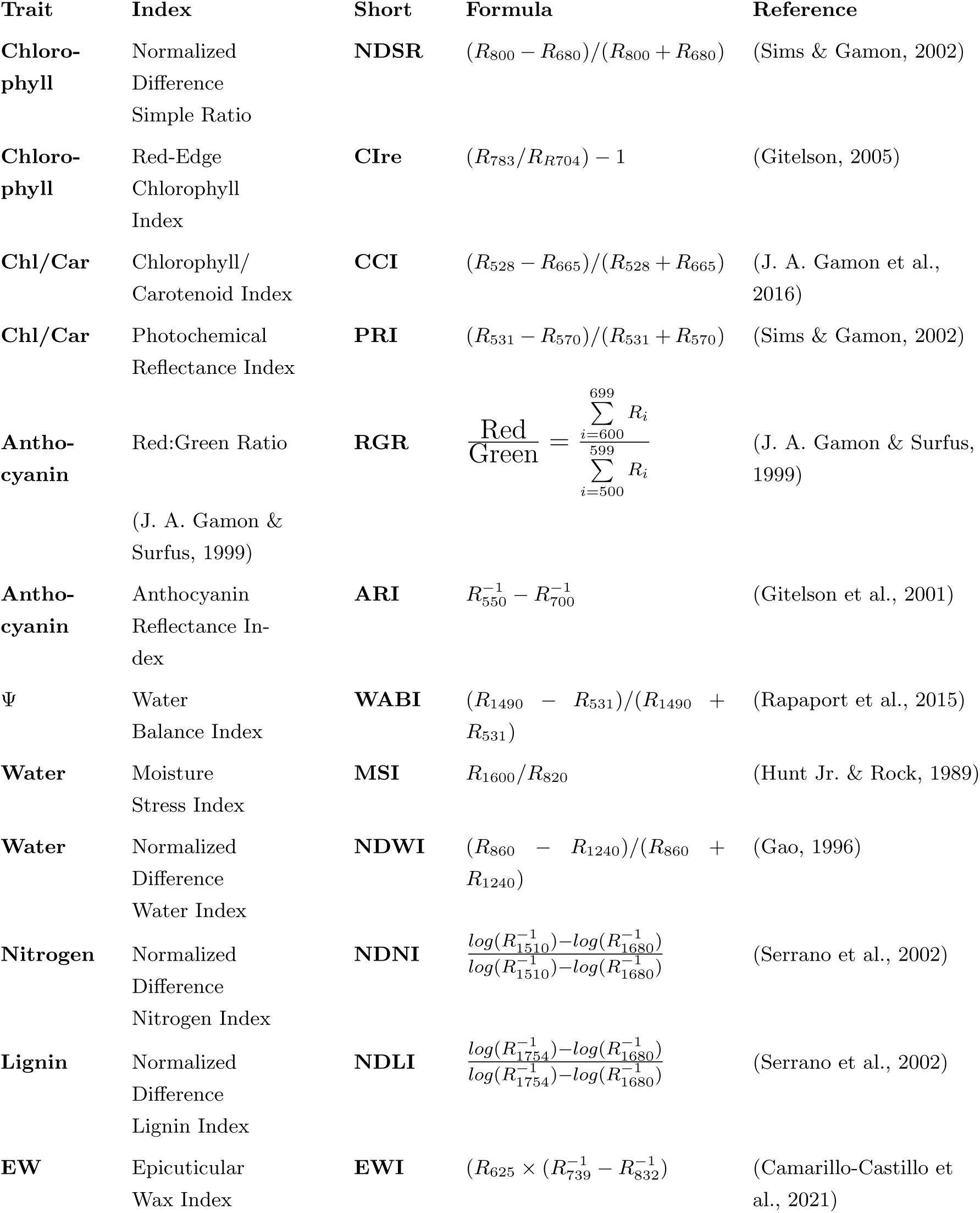
Summary of spectral indices including equation, author, and year of publication. *R_λ_* indicates reflectance at wavelength *λ*.

Chlorophylls absorb in blue and red regions capturing light for photosynthesis, with a maximal absorbance at 670 nm. Due to spectral interference in blue with carotenoids as well as anthocyanins in green, many empirical models use reflectances in the 700 nm regions (Lichtenthaler et al., 1996; Datt, 1999; Buschmann & Nagel, 1993). The NDSR Sims and Gamon (2002) considers *R* from the centre of the broad range bands of the often used normalized difference vegetation index (NDVI). CIre on the other hand is based on the transitional wavelength position between low reflectance in red and high reflectance in NIR.

It is challenging to estimate carotenoid content through indices due to the overlap between chlorophyll and carotenoid absorption peaks. It is easier to estimate the ratio of carotenoid to chlorophylls for relative carotenoid content. The CCI was developed to track seasonally dependent photosynthetic rates J. A. Gamon et al. (2016). Similarly, the PRI is a proxy for photosynthetic light use efficiency (Sims & Gamon, 2002). Both are carotenoid-sensitive indices and practical tools to monitor photosynthetic activity (Sasagawa et al., 2022).

As with carotenoids, the absorption of anthocyanins with a peak absorption at around 550 nm coincides with chlorophyll absorption, leading to comparable challenges in their estimation (Sims & Gamon, 2002). The RGR is based on the red:green ratio and can estimate the anthocyanin to chlorophyll ratio (J. A. Gamon & Surfus, 1999). The ARI (Gitelson et al., 2001) allows the accurate estimation of anthocyanin accumulation in stressed leaves.

Leaf water potential Ψ is an indicator of the whole plant water status and indicates how a plant may respond to water stress (Rodriguez-Dominguez et al., 2022). The WABI is a spectral index used for proximally tracking water status changes in grapevines (Rapaport et al., 2017). Similarly, the MSI correlates to relative water content to infer moisture stress (Hunt Jr. & Rock, 1989). Equivalent water thickness (EWT) is associated with leaf-level water status and thus sensitive to dryness stress and can be calculated either by directly measuring the variables or estimated with indices (Féret et al., 2019). Finally, we used the NDWI to estimate water content (Gao, 1996; Xu, 2006).

Foliar nitrogen and foliar lignin concentration are usually estimated when assessing ecosystem processes such as growth and decomposition. They have an absorption peak at 1510 nm and 1754 nm, respectively. NDNI and NDLI effectively indicated leaf nitrogen and lignin in shrub vegetation (Serrano et al., 2002). Epicuticular wax in plants may be considered as the first line of protection against biotic and abiotic stressors, with the EWI being the most accurate narrowband index to estimate waxes(Camarillo-Castillo et al., 2021).

To run PROSPECT-D, we employed the *prospect* package (v1.3.0) by Féret et al. (2017) in the R statistical software (v4.3.1) (R Core Team, 2023). To use the inversion and thus to estimate the output parameters, we called the function *Invert PROSPECT()*, which applies the iterative optimization algorithm in the *fmincon()* function included in the *pracma* package (v2.4.2). We used the root-mean-squared error (RMSE) merit function, which minimizes the RMSE between the simulated and measured properties. We employed PROSPECT version D over previous versions because of its improved trait parametrization over a larger database and the inclusion of relevant traits such as anthocyanin content (Jacquemoud & Ustin, 2019). We followed the recommendations of Spafford et al. (2021) to obtain prior information on the leaf structure parameter *N* when only *R* is measured by using the integrated function *Get Nprior()* because *a priori N* estimation is expected to improve estimations of leaf traits. We estimated the following leaf traits using PROSPECT-D: *C_ab_*, *C_car_*, *C_ant_*, EWT, and LMA.

#### 2.8.2 Statistical analyses

Since we measured two leaves per sapling, we took the mean of the indices and the model outputs. The statistical analysis was conducted using the statistics software R (v4.3.1) (R Core Team, 2023). We used the linear model function *lm()* of the *stats* package (v3.6.2) to fit the model and conducted Analysis of Variance (ANOVA) with the function *anova()* on the model. The response variables (i.e. the spectral indices and the model outputs) were regressed over the predictor variables height, treatment, genetic cluster, and the interaction treatment × cluster. For the ANOVA, we assumed randomness in the observations, homogeneous variances in each group and normality in the response variable. To ensure we met these assumptions, we conducted a visual inspection of the histogram using *hist()*, residuals versus fits, and Q-Q-plots (see Fig S12) using *qqPlot()*. We discovered that it was necessary to exclude those saplings that exhibited strong wilting (wilting index ≥ 2) or negative values in the normalized difference water index (NDWI *<* 0) to meet model assumptions (n=10) because they showed values strongly deviating from the main population of 180 seedlings, likely because they experienced a much stronger stress response. Square root and logarithmic transformations were not successful in improving the model Q-Q-plots when outliers were included (see Table S6 and Figure S12). These outlier values were, however, still included in the calculation of the log response ratio (LRR) that is robust to such effects. The individuals excluded from the ANOVA belonged to multiple genetic clusters (*n_cluster_*_2_ = 3, *n_cluster_*_3_ = 7) and sites (1 x BES, 1 x ROS, 3 x RSF, 1 x FRM, 2 x HBV, 1 x PLW, 1 x CHL). We ran the analyses with and without outliers and although some p-values and statistical significances for the terms ”height” and ”drought treatment” changed, the main results, i.e., the direction and magnitude of the effect sizes, were not affected (see Table S6 in the Supporting Information). We considered *p* ≤ 0.05 as our significance threshold.

Next, we calculated effect sizes for each spectral index and modelled trait for all individuals, including the 10 individuals that were strongly affected by the drought. Log response ratios (LRR) quantify the log-proportional change of the mean of the treatment and the control group (Lajeunesse, 2016). A strength of this method is that it linearizes its metric, treating the deviations in the numerator and denominator in the same way. To calculate the LRR and subsequently compute the variations of a leaf trait between the clusters, we adapted the equation from Hedges et al. (1999) and Bakbergenuly et al. (2020):

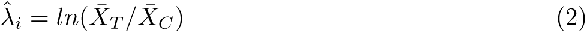

where 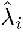 indicates the LRR and 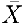 denotes the observed means of each treatment (*T*) and control (*C*) group. Since the logarithm is only defined for positive values and some indices show a series of negative and positive values, we apply an offset transformation by adding a constant *c*, where *c* equals 1 except for ARI, where *c* equals to 3; we inverted MSI for interpretability (see index definitions in Table 2).

The standard error of the LRR *S_E_* was computed as:

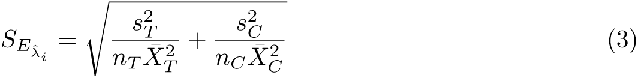

where *s* denotes standard deviations and *n* the sample sizes of the groups. We incorporated the outliers in the computation and visualization of the LRR (but see Fig. S8 for a visualization excluding the outliers).

### 2.9 Sequencing data and kinship matrix

Reads containing adapter sequences and those with low-quality scores were removed from the raw data using Trimmomatic (v0.39) (Bolger et al., 2014). To avoid bias introduced by using a reference genome, we used a reference-free method. This allows capturing all genetic variation among individuals without losing variation that is not present in the reference genome. Trimmed sequencing reads were analysed using the kmerGWAS pipeline (v0.2) (Voichek & Weigel, 2020) according to the manual found on github (https://github.com/voichek/kmersGWAS). The k-mer database was built using KMC (v3) (Kokot et al., 2017) with the default k-mer size and settings (-k 31 –mac 5 -p 0.2), equivalent to the approach in (Schmid et al., 2024). The k-mer table containing the presence/absence pattern of each 31 bp k-mer in the population was then used to derive the kinship matrix. The kinship matrix (Fig. S5) was calculated using efficient mixed-model association (EMMA) (Kang et al., 2008) with a minor allele frequency (MAF) of 0.05. The resulting matrix showing genetic relatedness was visualized with a heatmap and dendrogram with the *ASRgenomics* (v1.1.3) package in R using hierarchical clustering and Euclidean distances (Gezan et al., 2022). We then used PLINK (v1.9) (Purcell et al., 2007; Chang et al., 2015) to convert the k-mer table to a bed format and used PLINK to generate a PCA using the .bed file with the following options in effect: double-id –allow-extra-chr –pca –allow-no-sex–out. Visual inspection of the kinship matrix indicated the presence of three separate genetic clusters. These corresponded well with the findings of other recent studies investigating genetic relatedness based on SNPs among beech trees across Europe (Milesi et al., 2024; Lazic et al., 2024). Note that some discrepancies between our study and others may be due to the used settings (–mac 5) and the MAF being set to 0.05 in this study. This resulted in any k-mers that are private to populations consisting of fewer than five individuals (ITB, ITM, ITE and PLB) being excluded; however, reducing these threshold values also introduces noise and leads to more complex clustering that is difficult to interpret. Based on the kinship matrix and PCA (Fig. S5, Fig. S6), we assigned every sample (i.e., sapling) a genetic cluster and used these three clusters (annotated as cluster 1, cluster 2 and cluster 3) as predictor variables for further analyses. We note that two samples could not be properly processed using the k-mer pipeline, likely due to insufficient quality of their sequences. Consequently, we assigned them to their genetic clusters based on where their half-siblings were positioned in the kinship matrix. PLW 1 4 was assigned to cluster 3 because all half-sibs clustered together, and ESP 8 6 was assigned to cluster 1 because all of its half-sibs were in that cluster. The genetic clusters are visualised geographically in Fig. 1a.

## 3 Results

### 3.1 Soil moisture and stomatal conductance

The soil moisture sensors showed significantly lower moisture for the drought treatment (mean TDT value: 1562) compared to the control group (mean TDT value: 1906), indicating that rain exclusion had resulted in 18.0 (± 10.25) percent drier soils in the drought treatment group (Fig. S2a). Furthermore, we found an inverse relationship between soil moisture and tree height for both control and treatment groups, suggesting that taller saplings tended to have less soil moisture in their pots (*R*^2^ = 0.42, *p* = 0.059 for the treatment group and *R*^2^ = 0.52, *p <* 0.05 for the control group, Fig. S7a).

The drought treatment decreased stomatal conductance (*g_s_*) (*F*_(1,134)_ = 43.0879, *p <* 0.001). As with soil moisture, stomatal conductance was also generally lower for taller saplings (Fig. S7b, (*F*_(1,134)_ = 68.6692, *p <* 0.001). Stomatal conductance did not differ between the genetic clusters (Table 3, *F*_(2,134)_ = 1.5842, *p* = 0.21) and the interaction between genetic cluster and drought treatment was also not significant (Table 3, *F*_(2,134)_ = 0.2493, *p* = 0.78).

**Table 3.** ANOVA table for the linear models with spectral leaf traits as dependent observations and sapling height, treatment, genetic cluster, and treatment × genetic cluster interaction as explanatory factors. *p* ≤ 0.001 (*^′^*∗ ∗ ∗*^′^*), *p* ≤ 0.01 (*^′^*∗∗*^′^*), *p* ≤ 0.05 (*^′^*∗*^′^*).

| Variable | Factor | DF | F value | P value |
| --- | --- | --- | --- | --- |
| NDSR | Height | 1 | 0.0677 | 0.7950 |
|  | Treatment | 1 | 20.2845 | 1.267e-05 *** |
|  | Cluster | 2 | 0.1387 | 0.8706 |
|  | Treatment x Cluster | 2 | 0.6807 | 0.5077 |
|  | Residuals | 163 |  |  |
| CIRE | Height | 1 | 8.7537 | 0.003550 ** |
|  | Treatment | 1 | 15.4489 | 0.000125 *** |
|  | Cluster | 2 | 0.2193 | 0.803280 |
|  | Treatment x Cluster | 2 | 0.6183 | 0.540137 |
|  | Residuals | 163 |  |  |
| CCI | Height | 1 | 1.5056 | 0.2215783 |
|  | Treatment | 1 | 15.6351 | 0.0001143 *** |
|  | Cluster | 2 | 0.2534 | 0.7764515 |
|  | Treatment x Cluster | 2 | 0.3494 | 0.7056659 |
|  | Residuals | 163 |  |  |
| PRI | Height | 1 | 6.5809 | 0.01121 * |
|  | Treatment | 1 | 33.8565 | 3.059e-08 *** |
|  | Cluster | 2 | 0.2516 | 0.77782 |
|  | Treatment x Cluster | 2 | 0.2650 | 0.76757 |
|  | Residuals | 163 |  |  |
| RGR | Height | 1 | 0.0162 | 0.8988 |
|  | Treatment | 1 | 27.7693 | 4.278e-07 *** |
|  | Cluster | 2 | 0.1018 | 0.9033 |
|  | Treatment x Cluster | 2 | 0.6076 | 0.5459 |
|  | Residuals | 163 |  |  |
| ARI | Height | 1 | 12.4887 | 0.0007106 *** |
|  | Treatment | 1 | 4.8022 | 0.0298421 * |
|  | Cluster | 2 | 0.8665 | 0.4223320 |
|  | Treatment x Cluster | 2 | 0.1251 | 0.8825305 |
|  | Residuals | 163 |  |  |
| MSI | Height | 1 | 3.7758 | 0.05372 |
|  | Treatment | 1 | 3.2350 | 0.07393 |
|  | Cluster | 2 | 0.2720 | 0.76219 |
|  | Treatment x Cluster | 2 | 0.6506 | 0.52310 |
|  | Residuals | 163 |  |  |
| NDWI | Height | 1 | 2.6809 | 0.10349 |
|  | Treatment | 1 | 6.0518 | 0.01494 * |
|  | Cluster | 2 | 0.0714 | 0.93116 |
|  | Treatment x Cluster | 2 | 0.0756 | 0.92721 |
|  | Residuals | 163 |  |  |
| WABI | Height | 1 | 3.9269 | 0.0492 * |
|  | Treatment | 1 | 20.7369 | 1.027e-05 *** |
|  | Cluster | 2 | 0.2105 | 0.8104 |
|  | Treatment x Cluster | 2 | 1.0836 | 0.3408 |
|  | Residuals | 163 |  |  |
| NDNI | Height | 1 | 1.6635 | 0.1990 |
|  | Treatment | 1 | 0.6213 | 0.4317 |
|  | Cluster | 2 | 1.0973 | 0.3362 |
|  | Treatment x Cluster | 2 | 0.5186 | 0.5963 |
|  | Residuals | 163 |  |  |
| NDLI | Height | 1 | 0.1488 | 0.70014 |
|  | Treatment | 1 | 4.8308 | 0.02936 * |
|  | Cluster | 2 | 1.3676 | 0.25763 |
|  | Treatment x Cluster | 2 | 0.5083 | 0.60245 |
|  | Residuals | 159 |  |  |
| EWI | Height | 1 | 21.7964 | 6.299e-06 *** |
|  | Treatment | 1 | 1.5499 | 0.2149 |
|  | Cluster | 2 | 0.1373 | 0.8718 |
|  | Treatment x Cluster | 2 | 0.0009 | 0.9991 |
|  | Residuals | 163 |  |  |
| $C_{ab}$ | Height | 1 | 6.9006 | 0.009439 ** |
|  | Treatment | 1 | 21.8055 | 6.272e-06 *** |
|  | Cluster | 2 | 0.0749 | 0.927911 |
|  | Treatment x Cluster | 2 | 0.8407 | 0.433285 |
|  | Residuals | 163 |  |  |
| $C_{car}$ | Height | 1 | 1.4615 | 0.2284 |
|  | Treatment | 1 | 16.2562 | 8.478e-05 *** |
|  | Cluster | 2 | 0.1950 | 0.8230 |
|  | Treatment x Cluster | 2 | 1.1605 | 0.3159 |
|  | Residuals | 163 |  |  |
| $C_{ant}$ | Height | 1 | 3.6239 | 0.05872 |
|  | Treatment | 1 | 0.7938 | 0.37426 |
|  | Cluster | 2 | 0.6886 | 0.50374 |
|  | Treatment x Cluster | 2 | 0.0761 | 0.92673 |
|  | Residuals | 163 |  |  |
| EWT | Height | 1 | 0.8154 | 0.3679 |
|  | Treatment | 1 | 1.7213 | 0.1914 |
|  | Cluster | 2 | 0.4006 | 0.6706 |
|  | Treatment x Cluster | 2 | 0.7447 | 0.4765 |
|  | Residuals | 163 |  |  |
| LMA | Height | 1 | 0.3761 | 0.5406 |
|  | Treatment | 1 | 2.5040 | 0.1155 |
|  | Cluster | 2 | 0.9626 | 0.3840 |
|  | Treatment x Cluster | 2 | 1.0906 | 0.3385 |
|  | Residuals | 163 |  |  |
| $g_s$ | Height | 1 | 68.6692 | 1.066e-13 *** |
|  | Treatment | 1 | 43.0879 | 1.053e-9 *** |
|  | Cluster | 2 | 1.5842 | 0.2089 |
|  | Treatment x Cluster | 2 | 0.2493 | 0.7797 |
|  | Residuals | 134 |  |  |

Although sapling height did not differ significantly between the genetic clusters (*F*_(2,177)_ = 0.223, *p* = 0.8, we found significant relationships between sapling height and stomatal conductance as well as soil moisture, which is why we included sapling height as a covariate in other analyses (Table 3).

### 3.2 Leaf spectral trait variation in response to drought

The leaf spectra showed on average increased reflectance for the drought-exposed saplings (Fig. 2a). This was more pronounced in genetic clusters 2 and 3 (Fig. 2c, d), and less so in genetic cluster 1 (Fig. 2b). The variation among the individuals also increased in the drought treatment, again, especially for genetic clusters 2 and 3.

#### 3.2.1 Leaf pigments

Two out of the three main pigment classes as assessed using spectral indices (Table 2) were significantly reduced in the drought treatment (Table 3, Fig. 2). We observed a negative direction for chlorophyll-related (Fig. 3a) and carotenoidrelated indices (Fig. 3c), but the opposite in anthocyanins derived from indices, which exhibited either a neutral or a positive effect size (ARI and RGR, Fig. 3e).

**Figure 3.**
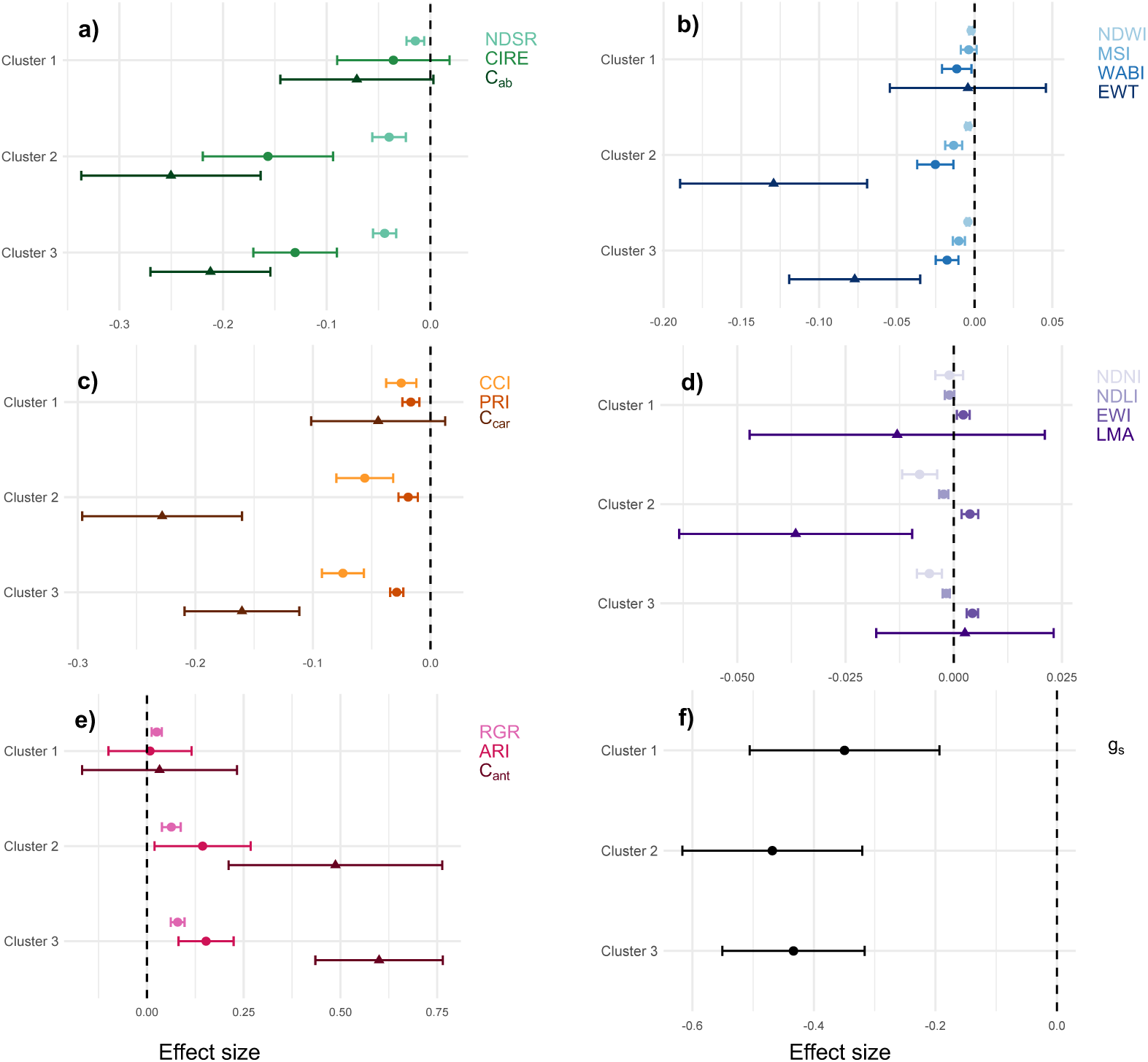
Log response ratios of spectral indices, PROSPECT-D simulated leaf traits and stomatal conductance (mean ± standard error; Table 2 for trait definitions). A negative effect size indicates a negative response to the experimental drought and vice versa. a) Indices related to chlorophylls. NDSR: Normalized Difference Chlorophyll Index, CIRE: Red-edge clorophyll index. *C_ab_*: simulated chlorophyll a+b content. b) Traits related to water content and stress.

In addition, we found a significant reduction in the PROSPECT-D modelled leaf pigments *C_ab_* and *C_car_* but, in line with the results from the spectral indices, a nonsignificant increase in drought-treated saplings for *C_ant_*. The pigment ratio between *C_ab_* and *C_car_* was significantly reduced between the control and the treatment group (*F*_(1,163)_ = 8.4248*, p <* 0.01). CIRE and PRI showed a significant negative association with increasing sapling height, while ARI exhibited a significant positive association with increasing sapling height, indicating more anthocyanin in taller trees.

Genetic cluster did not have a significant effect on any of the leaf pigments, nor did the treatment × genetic cluster interaction (Table 3). There was thus significant variation between the control and treatment groups, but not between genetic clusters and their response to drought, as assessed by leaf spectroscopy. Despite the absence of significant effects in the ANOVA, genetic cluster 1, which is mostly composed of Spanish provenances, showed a lower magnitude of drought response in several traits compared to the other genetic clusters (Fig. 3). The LRRs were all close to zero and the standard errors overlapped.

In contrast, the other two genetic clusters had LRRs deviating further from zero, indicating a stronger response. Furthermore, spectral indices responded differently and did not always match the response of the corresponding modelled trait. For example, the LRR for NDSR was consistently less negative and variable than the LRR for simulated *C_ab_*. However, the LRR of CIRE was comparable to the LRR of *C_car_*, with overlapping standard errors. The LRRs for the modelled *C_car_* were more negative and more variable than the LRRs for either carotenoid index (CCI or PRI), i.e., the carotenoid indices did not follow the drought response of modelled carotenoids. The carotenoid indices showed distinct patterns: CCI exhibited a more negative LRR with a larger standard error than PRI. Likewise, LRRs for modelled *C_ant_* were consistently higher and more variable than those for either anthocyanin index (RGR or ARI), though the standard error of the ARI LRR overlapped with that of *C_ant_* in cluster 2. The LRRs of RGR and ARI were generally consistent across clusters, with ARI being more variable.

#### 3.2.2 Leaf water potential and water content

Drought-stressed saplings had lower NDWI and water potential index (WABI), but the moisture stress index (MSI) and modelled EWT were not affected (Table 3, and see Table 2 for indices). Again, we did not find a statistically significant difference between the genetic clusters and their interaction with the drought treatment. We found a negative drought response in these metrics of water content and water potential for all three genetic clusters (Fig. 3b). In line with the results of the leaf pigments, we observed only a slightly negative response in EWT in genetic cluster 1, revealing no water content changes between the control and the treatment saplings, and the MSI was lower in genetic cluster 1 as well (Fig. 3b). As for the pigments, for genetic cluster 1, the standard errors of the LRRs for modelled water-related traits and related indices overlapped and LRRs were all close to zero. In contrast, for clusters 2 and 3, LRRs for modeled EWT were consistently more negative and more variable than for any of the water-related indices, although modelled EWT was not significantly affected by drought due to its variability. NDWI, while significantly affected by drought, had a small magnitude of LRR, leaving WABI, with larger LRRs and a significant change with drought, the clearest water-related indicator of sapling drought response in this dataset.

#### 3.2.3 Other structural and biochemical leaf traits

The drought treatment had a significant effect on lignin content, as assessed with the spectral index NDLI. In contrast, nitrogen content, epicuticular wax (EWI), and LMA were not significantly affected by the drought treatment as assessed by spectral indices and modeling (Fig. 3a, Table 3, and see Table 2 for indices). Again, there was no significant interaction between treatment and genetic cluster. EWI increased with drought in all three genetic clusters, suggesting an increase in epicuticular wax on the leaves, while modelled nitrogen and lignin decreased. For modelled LMA, we found a negative response to drought, although it was not significant. Sapling height was correlated with EWI, but not with any of the other structural leaf spectral traits (Table 3).

### 3.3 Validation for PROSPECT-D inversion output

We conducted chromatographic separation for 20 leaf samples, and identified main compounds based on retention times and standard spectra. We regressed all the measured traits to the simulated traits from the PROSPECT-D inversion (Fig. 4). We found a strong and significant relationship in both chlorophylls and carotenoids (chlorophyll (*C_ab_*) (*R*^2^ = 0.73, *p <* 0.001), carotenoids (*C_car_*) (*R*^2^ = 0.74*, p <* 0.001), Fig. 4a, b) and a less strong but still significant positive correlation for EWT and LMA (EWT (*R*^2^ = 0.69, *p <* 0.001), LMA (*R*^2^ = 0.51, *p <* 0.001), Fig. 4c, d). For the pigments, we also found a clear separation of leaf samples from the treatment and control pool, with the drought-treated group exhibiting a smaller pigment concentration per area (Fig. 4). However, there was systemic overestimation in both simulated *C_ab_*and *C_car_* (*β*_1_ *>* 1).

**Figure 4.**
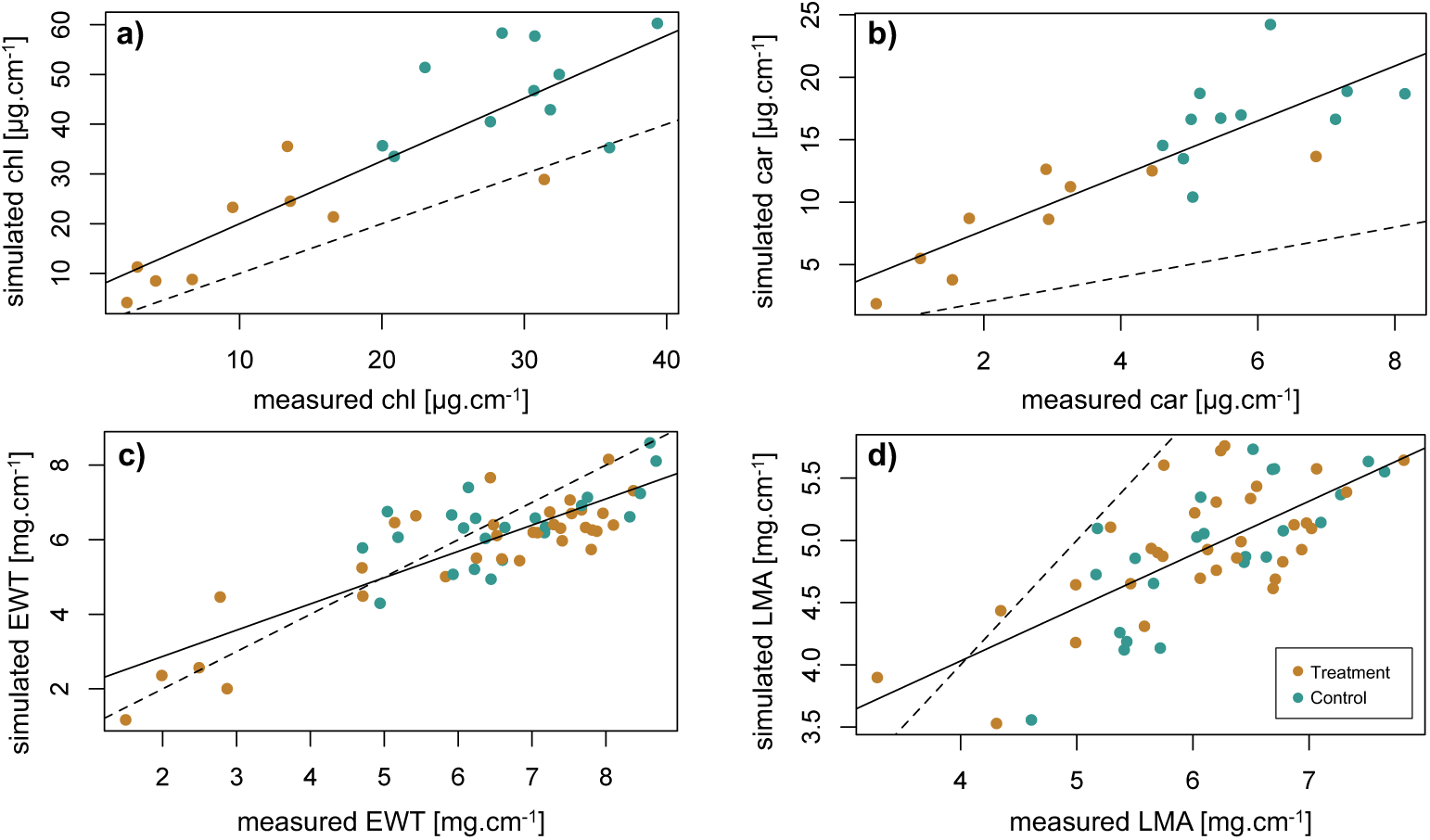
Measured leaf pigments (n=20) and leaf functional traits (n=56) regressed against PROSPECT-D inverse modelled traits. Dashed lines indicate *x* = *y*. a) Chlorophyll (*C_ab_*) (*R*^2^ = 0.73). b) Carotenoids (*C_car_*) (*R*^2^ = 0.74). c) EWT (*R*^2^ = 0.69). d) LMA (*R*^2^ = 0.51).

## 4 Discussion

### 4.1 Influence of drought treatment on spectrally derived traits related to pigments

The drought treatment altered the relative abundance of photosyntheticallyactive pigments estimated from spectral data. Leaf chlorophyll content is well known to be positively correlated with photosynthetic rate (Fleischer, 1935; Emerson, 1929). We observed a significant decrease in chlorophyll-related indices such as NDWI: Normalized Difference Water Index. MSI: moisture stress. Note that a more negative response reflects greater moisture stress. WABI: Water Balance Index (water potential). EWT: Equivalent Water Thickness. c) Indices related to carotenoids. CCI: Chlorophyll-Carotenoid Index. PRI: Photochemical Reflectance Index. *C_car_*: simulated carotenoid content. d) Leaf structural traits. NDNI: Normalized Difference Nitrogen Index. NDLI: Normalized Difference Lignin Index. EWI: Epicuticular Wax Index. LMA: Leaf Mass per Area. e) Traits related to anthocyanins. RGR: Red:Green Ratio. ARI: Anthocyanin Reflectance Index. *C_ant_*: simulated anthocyanin content. f) Stomatal conductance (*g_s_*). Note that the complete dataset was used for all shown LRRs.

NDSR and CIRE, corresponding to a decrease in PROSPECT-D simulated *C_ab_* in drought-treated saplings. Previous studies found a similar decrease in chlorophyll content per area in drought-stressed *F. sylvatica* with liquid chromatography or chlorophyll measurements (Tognetti et al., 1995; Galĺe & Feller, 2007; Arend et al., 2016). Galĺe and Feller (2007), for instance, found that the activity of PSII during drought stress is down-regulated, which is consistent with the reduction of chlorophyll in severely drought stressed leaves.

Carotenoid content was also significantly lower in drought-treated saplings compared to the control group. This is in line with previous findings (Galĺe & Feller, 2007; Junaid et al., 2023). However, to compensate for water stress, the photosynthetic apparatus is known to change carotenoid composition rather than total content. One may observe an increased ratio of xantophyll cycle pigments (VAZ), which are a division of the carotenoids, to chlorophyll content during water limitation (Demmig-Adams & Adams, 1992; Merzlyak et al., 1999; Biswal, 1995). VAZ contribute to dissipate excess energy in the apparatus during drought situations and are thus regarded as key pigments for photo-protection in *F. sylvatica* (Munńe-Bosch & Alegre, 2000). This is consistent with our findings where the PROSPECT-D modelled ratio *C_chl_/C_car_* declines in drought-treated saplings. Based on these observations, we infer that when plants were water stressed, the content of both chlorophylls and carotenoids declined correspondingly. However, the proportion between them changes differently. In agreement, we found a decline in PRI and CCI, which are spectral indices related to the chlorophyll and carotenoid ratio and linked to photosynthetic capacity (Filella et al., 2009). PRI and CCI both use the 531 nm band, which is indicative of the xanthophyll cycle epoxidation state transition (J. Gamon et al., 1992). Both were significantly decreased in drought-treated saplings.

Anthocyanins are thought to be involved in many plant protective functions (Gould, 2004), (Landi et al., 2015), as they mediate stressors (i.e., ROS) by antioxidant activity and light attenuation reducing excess energy, similar to carotenoids (Feild et al., 2001; Landi et al., 2015; Steyn et al., 2002). However, whether anthocyanins play a key role for plants, in particular beech, under water stress is still unknown, and there is little literature on anthocyanin accumulation in beech except for the copper varieties, which bear a mutated gene preventing them from degrading anthocyanins and are bred for ornamental purposes. We observed a significant increase of the anthocyanin-related indices RGR and ARI in drought-treated saplings, and also an increase of PROSPECT-D modelled *C_ant_*. This suggests a potential accumulation of anthocyanins under drought stress. However, the modelled *C_ant_* indicated a very low amount of *C_ant_* compared to the possible range of variation in PROSPECT-D (Jacquemoud & Ustin, 2019). Consistently, we only detected anthocyanins in a copper beech sample and not in the saplings used for the drought experiment, although our laboratory analysis was not optimized to be sensitive for anthocyanins, and we do not know the limit of detection. It is likely that anthocyanin accumulation in the leaves of our experimental saplings in response to drought was low in comparison to anthocyanin concentrations that have been measured in other plants and other conditions. Gitelson et al. (2009) successfully used ARI in maple (*Acer platanoides*) and dogwood (*Cornus alba*), and the index was also able to predict anthocyanin content in lettuce (*Lactuca sativa*)(C. Kim & Van Iersel, 2023) and grapevines (Steele et al., 2009). However, the applicability of ARI in beech has not yet been verified, and we therefore do not draw strong conclusions based on this index. Similarly, the RGR has so far, to our knowledge, not yet been applied or validated in *F. sylvatica* exposed to drought or other environmental stress. We thus presume that anthocyanin content was relatively low in our samples during the first drought period, and that we were unable to adequately quantify its change.

### 4.2 Influence of drought treatment on water content and leaf structural and biochemical traits

Moisture stress and water content were captured with MSI, which was higher in drought-treated saplings, and NDWI, which was lower in drought-treated saplings, as expected. Many studies concerning hydraulic function in leaves observed a similar increase in MSI, suggesting MSI to be a good indicator of altered plant water status (D. M. Kim et al., 2015; Zhang & Zhou, 2019). One of the most crucial counteracting mechanisms of plants to water deficiency is the rapid regulation of stomatal conductance to prevent plant dehydration. The closure of stomatal pores allow a decrease of gas exchange through reduced water vapor transpiration, and thus preserving water resources (Aranda et al., 2015; Wankmüller & Carminati, 2022; Pflug et al., 2018). As a moderately anisohydric species, *F. sylvatica* optimizes stomatal conductance under drought conditions (Peuke et al., 2002; Tardieu & Simonneau, 1998). As EWT was found to be similar in control and drought-treated saplings, this suggests that water content was preserved in the measured leaves. Another measure for hydraulic functioning and water dynamics throughout a drought is the leaf water potential (Ψ), the energy state of water in a system reflecting the water’s ability to move. Ψ varies in response to climate and soil conditions, and is strongly influenced by seasonal and diurnal changes (Bartlett et al., 2012; Czajkowski & Bolte, 2006; Dietrich et al., 2019), which resembles the anisohydric behaviour of *F. sylvatica* (Roman et al., 2015). WABI, which was significantly decreased in the drought-treated saplings, relates independent changes in photo-protective pigments involved in non-photochemical quenching (NPQ) (531 nm) and water absoprtion bands (around 1500 nm) which indicate stress-induced changes by Ψ (Rapaport et al., 2015). Thus, we infer that Ψ was indeed decreased in drought-treated saplings. The discrepancy between changes in Ψ and no changes in water content suggests that the saplings may be able to maintain relatively stable water content through rapid adjustments such as stomatal regulations while simultaneously varying their Ψ.

LMA can be understood as the leaf-level cost of light interception (Grime, 2001). An alternative way to define LMA is the accumulation per area of different classes of compounds, such as lignin or proteins (Poorter & Villar, 1997). We derived LMA through PROSPECT-D inversion and found no effect of the drought treatment. This was as expected, as sudden withholding of water and a short drought period may not be enough to modify leaf structure (Poorter et al., 2009). Similarly, we also did not find any drought-induced differences in modelled nitrogen content. However, it is likely that a longer drought duration may have led to an increased LMA (Poorter et al., 2009) and altered leaf nitrogen.

At the interface of the leaf towards the atmosphere, epicuticular waxes serve as a protective barrier against ultraviolet light exposure and uncontrolled water diffusion (Speckert et al., 2023). Consistent with our expectations, we observed a slight increase of EWI in drought-treated saplings, although this was not statistically significant. Epicuticular waxes are directly exposed to the environment and thus expected to adjust to abiotic stressors such as drought with the aim to conserve water and resist cellular dehydration in leaves (Shepherd & Wynne Griffiths, 2006). Studies using black pine (*Pinus nigra*) (Kreyling et al., 2014), grassland plant communities (Srivastava & Wiesenberg, 2018), and several shrubs and coniferous trees (Ofiti et al., 2023) increase renewal rates of waxes, which lead to a reduction of drought stress. However, the biosynthesis of waxes in *F. sylvatica* depends strongly on exposure to sun (Speckert et al., 2023). Deposition of wax as a response to water stress can occur swiftly within a few days (Bengtson et al., 1978). Our findings can thus be potentially explained by alternative water conservation mechanisms such as the aforementioned stomatal regulation to cope with the imposed water stress in *F. sylvatica*, or by changing the composition of the waxes rather than varying the concentration (Sachse et al., 2009).

As a major effect of our experimental conditions, we found that taller saplings generally had lower soil moisture in both control and treatment groups, potentially due to larger belowground biomass and increased water needs. We thus cannot exclude the possibility that taller saplings on average experienced a more severe drought. We addressed this limitation by controlling for sapling height within our block design and by adding sapling height as a factor in the statistical analyses.

We did observe a few significant (but not very strong) associations between sapling height and leaf traits, which would indicate that the responses to drought depended at least in part on sapling height. Along with the experimental conditions, foliar water uptake through permeable cuticles in *F. sylvatica* may have influenced the actual drought conditions of the saplings (Schreel et al., 2020).

### 4.3 Intraspecific variation in drought stress responses across genetic clusters

There are many studies on the role of intraspecific phenotypic variation in tree responses to drought (e.g. *F. sylvatica*; (Dounavi et al., 2016; Cocozza et al., 2016; Gonźalez De Andŕes et al., 2021; Leuschner, 2020; Carsjens et al., 2014; Thom et al., 2023; Baudis et al., 2014; F. Wang et al., 2021)), with the aim to improve decisionmaking in forest management and enhance forest resilience to climate change. We hypothesized that there is intraspecific variation in such drought responses depending on the sapling’s genetic background. Indeed, we demonstrated a variety of physiological responses to decreased water availability but, contrary to our expectations, we did not find that the genetic cluster strongly influenced the drought response.

However, there were indications of variation depending on the genetic cluster, particularly in genetic cluster 1, which showed a more resistant response in multiple traits. Thus, provenances from the Iberian Peninsula showed increased resistance to the imposed drought. This is perhaps not surprising, as this genetic cluster originates from a region that is characterized by low mean annual precipitation (638 mm). In contrast, the two other genetic clusters are from regions with more precipitation (815.25 mm and 913.2 mm annual rainfall, respectively). Although mean annual precipitation is generally used to define climatic niche, the number of consecutive dry days (CDD, defined as consecutive days with daily precipitation below 1 mm) is more directly comparable to our experimental design. The region (Araǵon) where the seeds of the trees from cluster 1 originated experienced on average 11.8 CDD in June and July from 1980 to 2010 (based on data provided by the Copernicus Climate Change Service, https://climate-adapt.eea.europa.eu/en/metadata/indicators/consecutive-dry-days). This is compared to the slightly lower CCD (on average 9.2 for cluster 2 and 10.8 for cluster 3) for the regions from which seed collections of the other two genetic clusters originated. However, conditions at the origin sites can deviate substantially from their regional averages. Furthermore, there are large variations among regions associated with a single cluster. Notably, seedlings belonging to cluster 3 originate from regions such as Sicily that experience up to 25 CDD in summer, as well as from regions that experience only 5 CDD on average (e.g. in Romania).

In sum, average climate information is not sufficient to indicate whether our findings are a signal of local adaptation. Drought response is a complex and environmentally contingent trait, and this study does not indicate whether genetically encoded adaptations to drought correspond with the identified genetic clusters. Rather, we conclude that the response of these seedlings to experimental drought varies along a suite of leaf functional traits and that this variation partly corresponds with geography of seed origin and overall genomic similarity. This suggests, but does not yet test, the hypothesis that seedlings belonging to genetic cluster 1 are evolutionary better adapted to withstand drought.

Other studies on intraspecific variation found performance differences in different populations of *F. sylvatica* in ecophysiological stress relevant to water stress, such as photosynthetic rate and stomatal conductance (Cocozza et al., 2016; Gonźalez De Andŕes et al., 2021; Śanchez-Gómez et al., 2013). However, phenotypic variation between populations or genetic clusters could often be masked by high within-population variation.Schmeddes et al. (2023) demonstrated the greatest phenotypic variation among *F. sylvatica* within the progeny of a single mother tree. Together, these and our findings indicate significant potential for both individual acclimation to climatic variability and population-level adaptation to drought.

### 4.4 Leaf spectroscopy can capture leaf phenotypic variation in drought response

Leaf spectroscopy offers a non-destructive, repeatable, and rapid measurement method for leaf traits compared to traditional methods and is thus a promising method in ecological research (Helsen et al., 2021; Messier et al., 2010; Siefert et al., 2015), with low measurement uncertainties at the leaf level (Petibon et al., 2021).

We want to highlight that despite cross-species assessment of leaf traits (i.e., EWT or LMA) (Asner et al., 2014), it remains unclear whether spectrally-derived traits models at the leaf level are accurate enough to reliably capture trait differences within species (Feilhauer et al., 2018; Girard et al., 2020). Yet, despite such uncertainties, we demonstrated that applying spectral indices to high resolution spectral data is a simple method for deriving leaf trait variation in response to an experimental drought, making it a valuable tool in ecological experiments, especially with high replication or experimental units. However, it is crucial to note that spectral indices do not necessarily provide mechanistic explanations: Spectral indices hardly disclose the leaf physiological processes and metabolic systems underpinning the drought responses (Rapaport et al., 2015; Curran, 1989). A different approach is the inclusion of relevant physical processes. The inversion of the PROSPECT RTM simulates leaf traits based on the physical processes of the interaction of electromagnetic radiation with leaves (Jacquemoud & Ustin, 2019). We performed a subsequent extraction and HPLC analysis proposed by Petibon and Wiesenberg (2022) to establish a chromatographic profile of complex photosynthetic pigments of drought-treated and control saplings to estimate chlorophyll and carotenoid concentrations, and also measured leaf mass per area and water content directly for a set of validation samples. Our validation data show a strong relationship between the simulated and extracted traits for chlorophyll and carotenoid content, and a moderate relationship for EWT and LMA. This finding aligns with Spafford et al. (2021), who found similar relationships in their dataset when *N* is estimated and only reflectance is used. The robustness of the retrieval of leaf traits may also depend on the availability of transmittance T data, which can be retrieved with an integrating sphere measurement device, for instance (Petibon et al., 2021). Both R and T combined carry the most information for PROSPECT inversion (Spafford et al., 2021); however, the combined measurement of reflectance and transmittance takes at least twice as long per leaf and is a destructive measurement, requiring leaves to be removed for measurement against a fixed integrating sphere (Laliberté & Soffer, 2018). We note that for the assessment of absolute pigment concentration, other techniques are required, such as the extraction methods that we performed on a subset of leaves. However, in an experiment with dozens, hundreds, or even thousands of individuals, as can be the case in ecological studies of species-level, intraspecific and genetic variation, and multiple measurement time points, leaf traits estimated from rapidly acquired spectral data may not only be more practical but also sufficient – or even advantageous, being better able to capture change and support sufficient replication. While the challenge is to meaningfully interpret spectral data, an increasing number of studies, including ours, support this (Meireles et al., 2020; Cavender-Bares et al., 2016; C. Li et al., 2023; Czyż et al., 2023).

## 5 Conclusions

Due to the increasing duration and frequency of droughts, there is growing concern that *F. sylvatica* may no longer be adapted to these novel environmental conditions, which challenges the persistence of European beech forests. Thus, understanding the ecophysiological processes that occur in response to increased drought stress is key. We detected signatures of changes in leaf traits such as photosynthetically active pigments and water potential in drought-treated saplings. However, we did not detect significant changes in signatures of leaf structural components such as nitrogen and epicuticular wax. Leaf spectroscopy offered a rapid and nondestructive way to simultaneously assess a suite of relevant traits from individual leaves, and we found the inversion of PROSPECT-D to be a robust approach to quantify changes in leaf traits in European beech saplings exposed to experimental drought. This approach could potentially be upscaled to canopy-level or airborne (imaging) spectroscopy to support drought response assessments for entire forests that are currently based on multi-band sensors (Sturm et al., 2022; Helfenstein et al., 2024). In particular, our results could provide valuable insights for newer satellite and airborne missions that use imaging spectroscopy with high spectral resolution by informing them about spectral indices and modelled constituents that are effective in assessing tree drought responses. We present a unique experimental study acting as a small-scale validation step.

Intraspecific variation in drought responses will likely be an important determinant of beech forest persistence and evolution. With saplings representing three broad genetic clusters from different geographic regions across the species range, we did not find significant variation in the directionality of the drought responses based on the genetic cluster. However, we detected significant variation of the three genetic clusters in the magnitude of their drought response. Together, this is consistent with a developing consensus that beech populations harbour both local plasticity and geographic differentiation in their response to drought, and that the species has the potential to persist in future drier environmental conditions (Cocozza et al., 2016; 1. F. Wang et al., 2021).

## Acknowledgments

This work was supported by the NOMIS grant “Remotely Sensing Ecological Genomics”. For technical assistance we thank Linda Binkert, Katja Pfister, Mike Werfeli, Marius Vögtli, Maarten Eppinga, Nicole Manser, Cheng Li and Matthias Furler. We thank the Hauenstein garden center in Rafz for sapling care and support.

## Conflict of interest statement

The authors declare no conflict of interest.

## Open Research

Data are publicly accessible on Data Dryad (https://doi.org/10.5061/dryad.cc2fqz6fm) and the code on Zenodo(10.5281/zenodo.14040529). Whole-genome sequences are

available on ENA (accession ID PRJEB77559).

## Author contributions

**Dave Kurath:** Conceptualization, Investigation, Methodology, Formal analysis, Data curation, Visualization, Writing original draft. **Sofia J. van Moorsel:** Conceptualization, Formal analysis, Project administration, Supervision, Investigation, Writing original draft. **Jolanda Klaver:** Conceptualization, Investigation, Methodology, Data curation. **Tis Voortman:** Investigation, Data curation, Writing review & editing. **Barbara Siegfried:** Investigation, Methodology, Writing review & editing. **Yves-Alain Brügger:**Investigation, Methodology. **Abobakr Moradi:** Investigation, Methodology. **Ewa A. Czyz:** Investigation, Writing review & editing. **Marylaure de La Harpe:** Investigation. **Guido L.B. Wiesenberg:** Methodology, Writing review & editing. **Michael E. Schaepman:** Funding acquisition. **Meredith C. Schuman:** Conceptualization, Project administration, Supervision, Resources, Writing review & editing.

## Notes

### Competing Interest Statement

The authors have declared no competing interest.

### Summary of Updates

Some minor modifications in response to the reviewers. For example, more information on the used spectral indices.

