## Supporting Information for "Leaf spectroscopy reveals drought response variation in *Fagus sylvatica* saplings from across the species’ range"

[2]Current address: Syngenta Crop Protection AG, Schaffhauserstrasse, 4334 Münchwilen, Switzerland

[3]Current address: University of Wisconsin-Madison, Department of Forest and Wildlife Ecology, 1630, Linden Drive, Russell Labs, Madison, WI 53706, United States

[4]Current address: Canton of Graubünden, Office for Nature and Environment, Ringstrasse 10, 7000 Chur, Switzerland

### Contents of this file

1. Figure S1. Overview block design and height distribution.
2. Figure S2. Local air temperature, soil temperature and precipitation.
3. Figure S3. Leaf area calculation in ImageJ.
4. Figure S4. Chromatograms of "Long Red", treatment, and control leaf.
5. Figure S5. Kinship matrix showing genetic relatedness among the 180 saplings.
6. Figure S6. PCA for the 180 saplings.
7. Figure S7. Height of the seedlings measured before the drought period against mean soil moisture values expressed as raw TDT values for treatment and control group.
8. Figure S8. Log response ratios excluding the outliers for comparison.
9. Supplementary methods: Seed collections.

10. Supplementary methods: Validation of the PROSPECT-D inversion.
11. Figure S9. Calibration curves from standards for the determination of pigment concentration.
12. Figure S10. Pigment standards spectra with absorption maxima used for the identification of pigments.
13. Figure S11. Spectra of pheophytin a and b sampled from a chromatogram when identifying unknown pigments.
14. Figure S12. Q-Q-plots for spectral traits with and without outlier removal.
15. Table S2. Summary seed collection.
16. Table S3. Timeline of irrigation during the drought period.
17. Table S4. ASD FieldSpec 4 instrument configurations.
18. Table S5. Gradient of used eluents in the sequential pigment extraction.
19. Table S6. ANOVA table including outliers for comparison.

**Additional Supporting Information (Files uploaded separately)**

1. Table S1. Detailed overview of seed collection and sites.

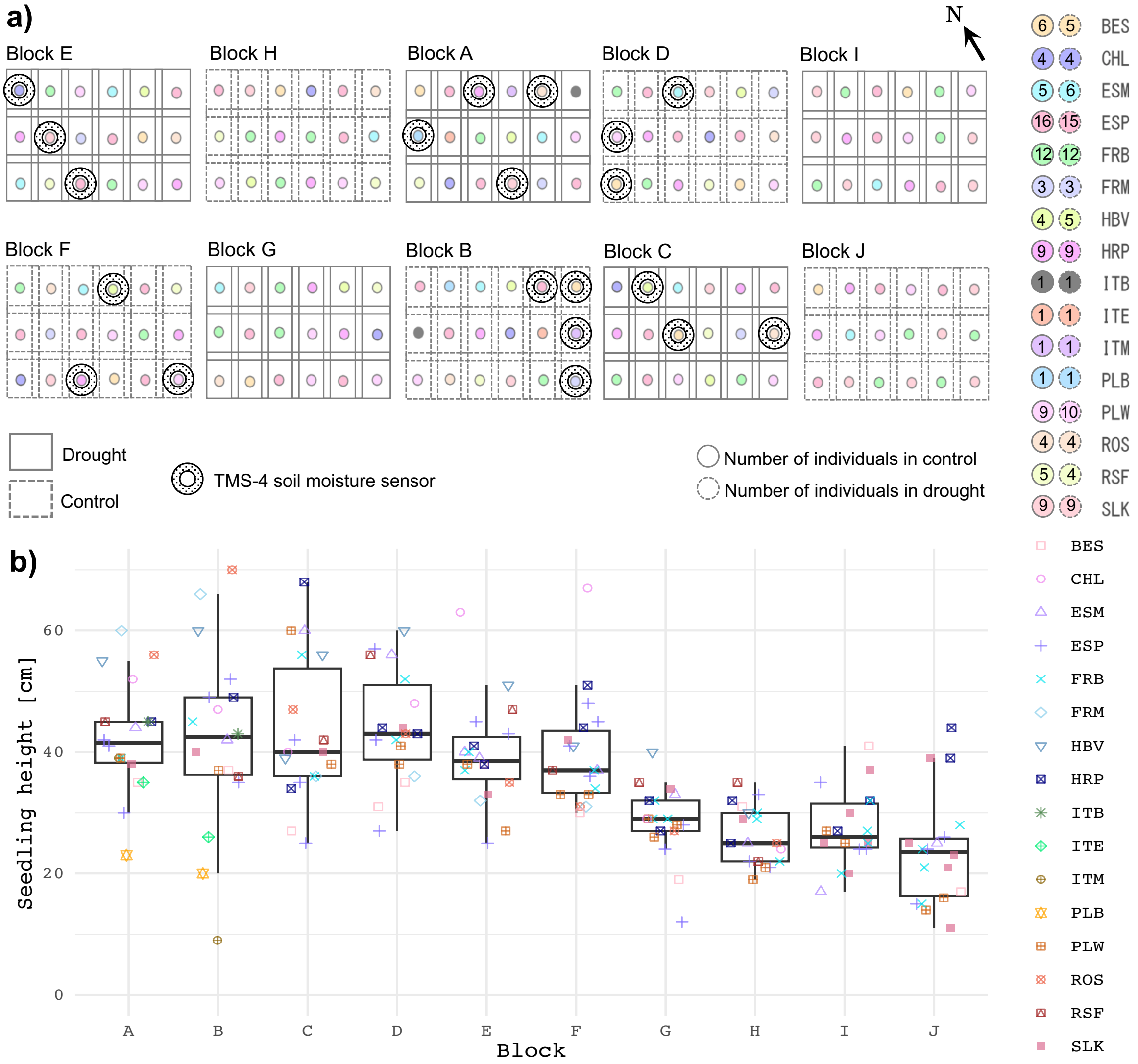


**Supplementary Figure S1.** a) Randomized representative block design with provenances for the common garden experiment and locations of TMS-4 sensors measuring soil moisture in a subset of the saplings throughout the experiment. North arrow indicates north direction. b) Sapling height distribution for each block. We attempted to have on average saplings of similar height assigned to the drought and the control treatments.

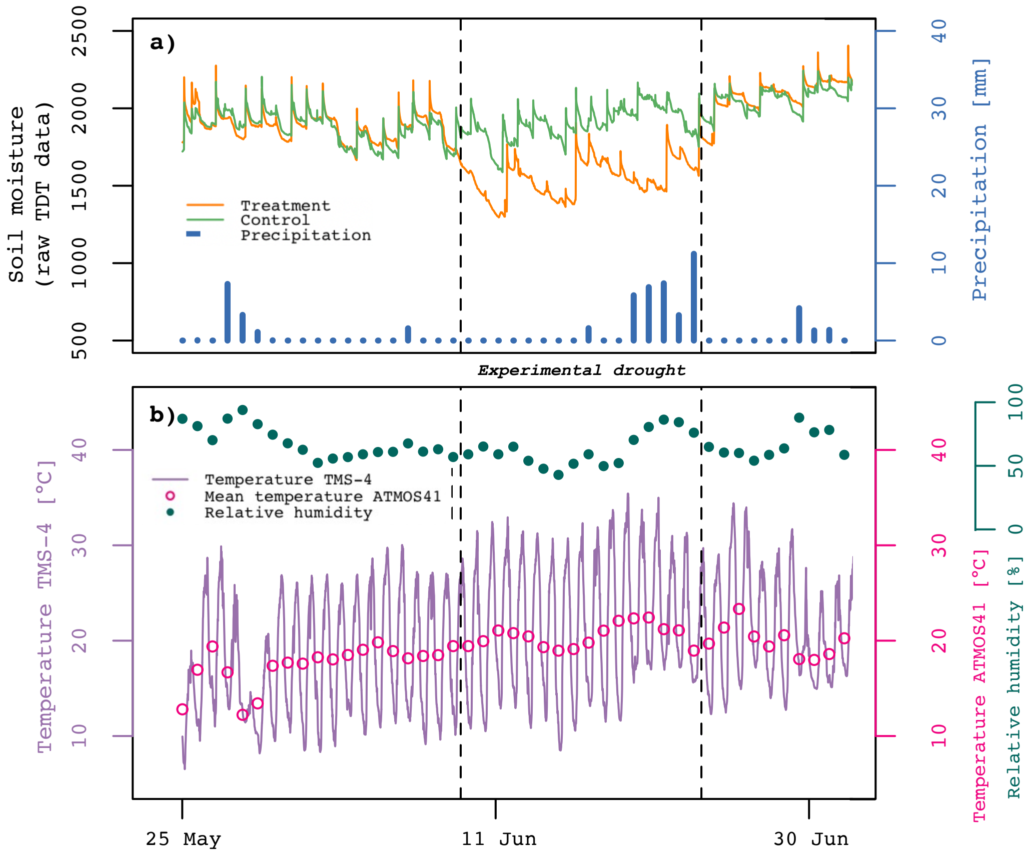


**Supplementary Figure S2.** Local air temperature, precipitation and relative humidity were measured throughout the experiment with ATMOS 41. a) Soil moisture was measured by TMS-4 probes, and raw time-domain transmission (TDT) values decreased in the treatment group. There were 6 days with precipitation accumulating a total of 36 mm of rain. b) During the drought period, mean temperature was 21.6° C (ranging from 8.9° C to 35.1° C) and the local air temperature measured by ATMOS 41 was 20.5° C. This discrepancy could have been due to the black rain covers absorbing the sunlight and thus warming up the soil temperature. Mean relative humidity was 63%.

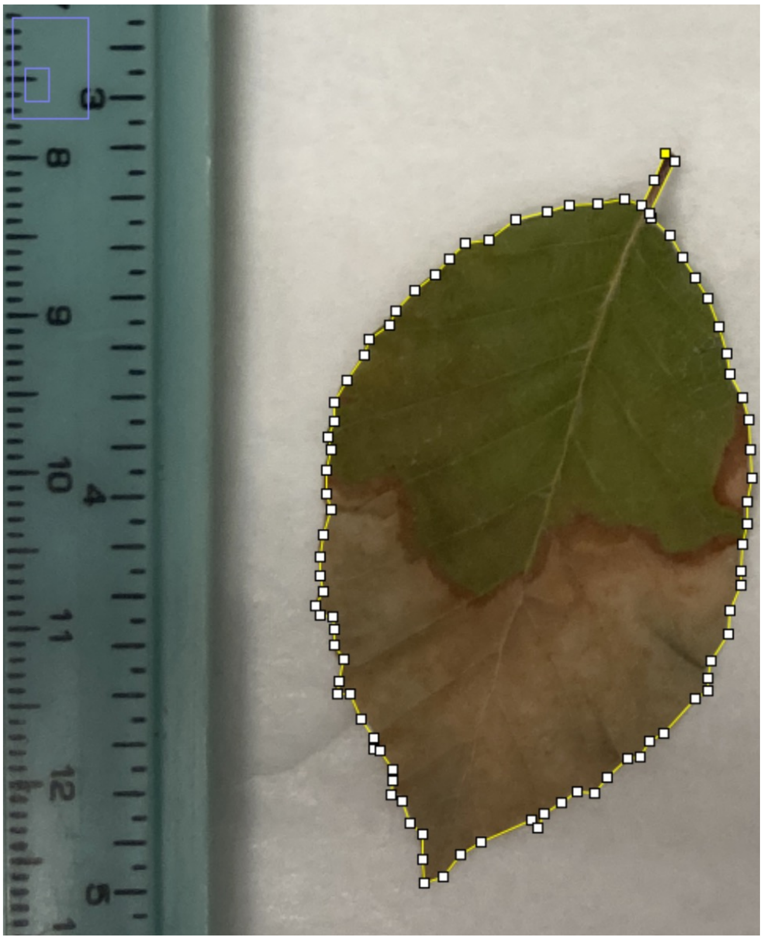

**Supplementary Figure S3.** Leaf area estimation in ImageJ (v1.38) (Schneider et al., 2012) using an above parallel photographed image of the leaf sample with a ruler with which we determined the real distance of a pixel. Using the point tool, we determined the region to calculate the area with the known distance.

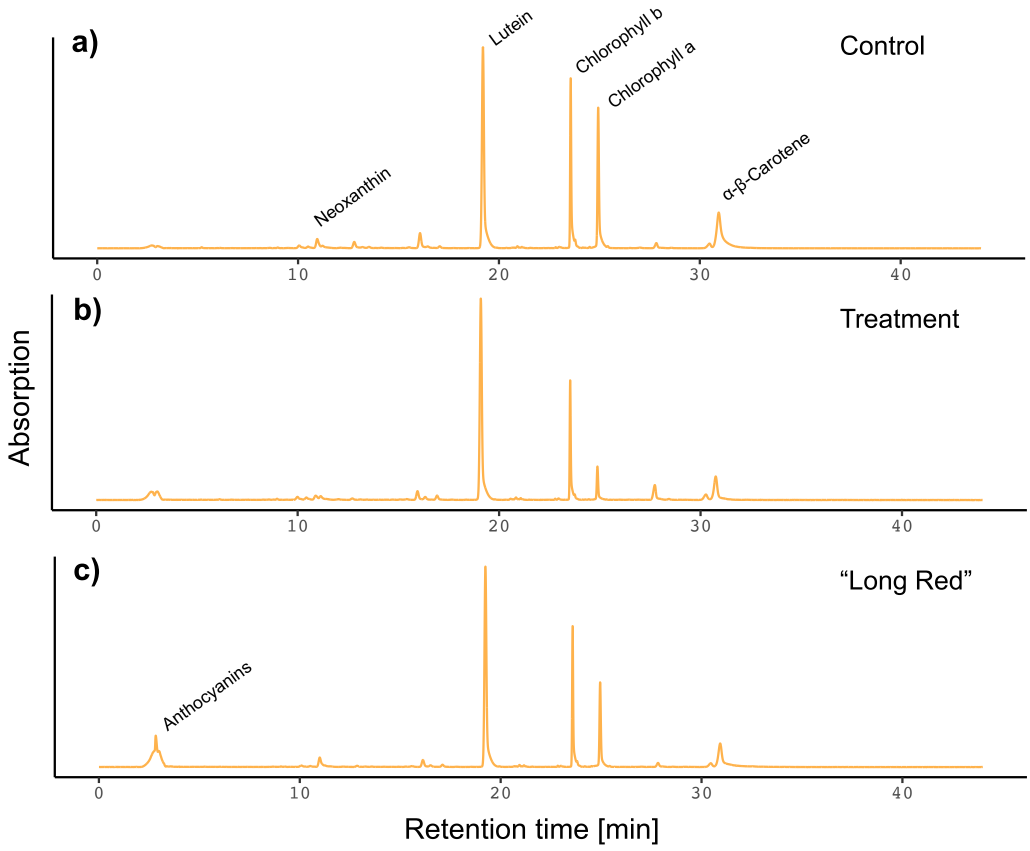


**Supplementary Figure S4.** Example chromatograms with wavelength = 450 nm of three leaves: a) Control leaf, b) Drought treatment leaf, c) ”Long Red” leaf. The rest of the leaves exhibited a similar pattern, lacking a strong peak for anthocyanins, except for ”Long Red”, which showed anthocyanin absorption.

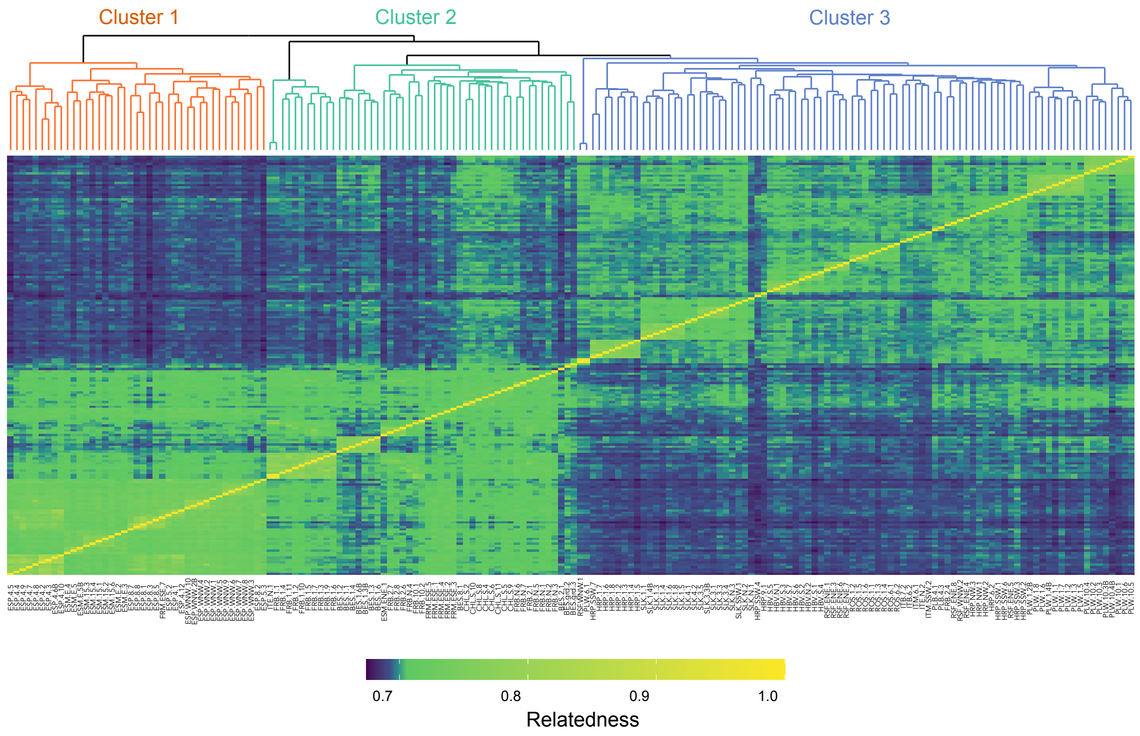


**Supplementary Figure S5.** The kinship matrix, showing genetic relatedness among the 180 studied beech saplings based on k-mers (31 bp). Yellow: high relatedness, blue: low relatedness. The matrix was visualized using hierarchical clustering and Euclidean distances. Visual inspection of the matrix indicated the presence of three separate genetic clusters, which we annotated as clusters 1, 2 and 3. See also Table S2 for an overview of the samples included in this analysis (except for one individual each from the collections PLW_1 and ESP_8).

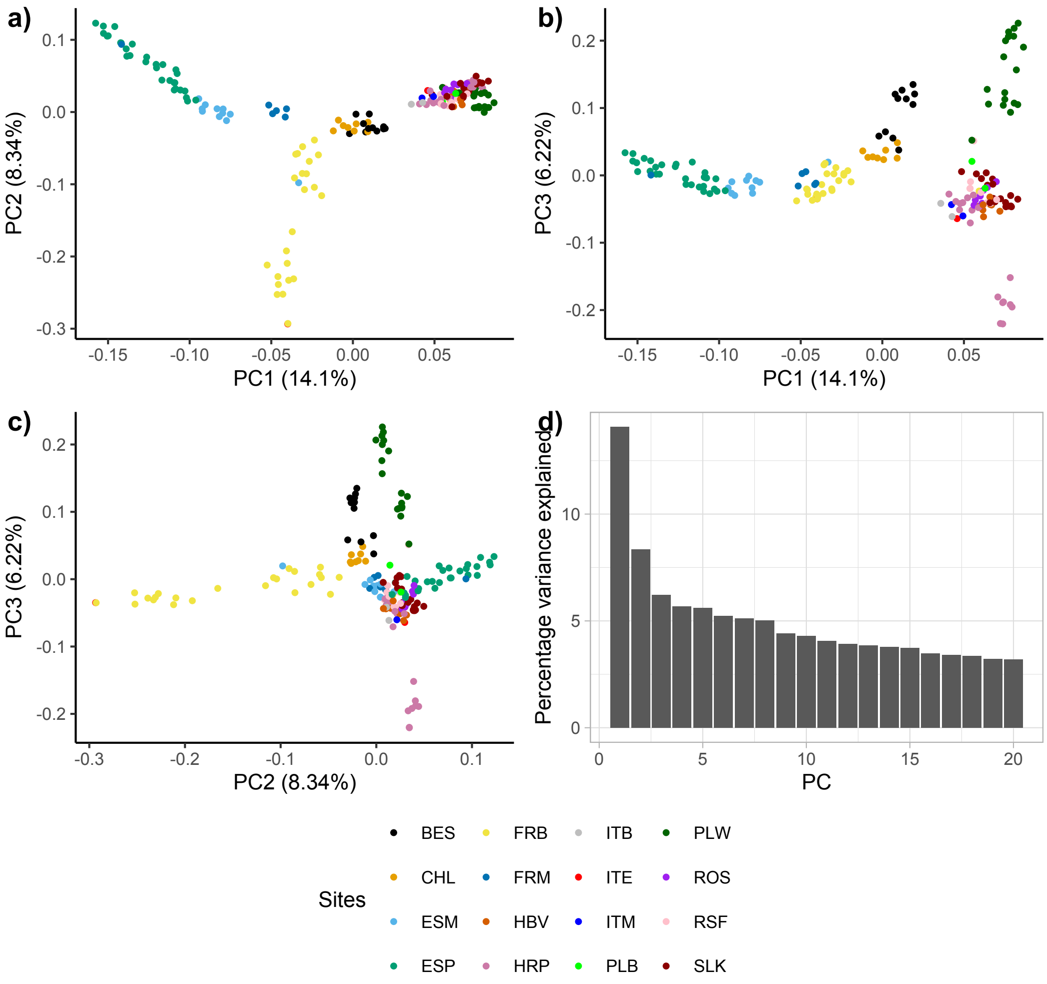


**Supplementary Figure S6.** Principal Component Analysis (PCA) color-coded by site to further illustrate the population genetic structure of the used beech saplings. a) PC1 and PC2 derived from the k-mer table. b) PC1 and PC3. c) PC2 and PC3. d) Percentage explained variation for the first 20 principal components. See also Table S2 for an overview of all samples included in the PCA (except for one individual each from the collections PLW_1 and ESP_8).

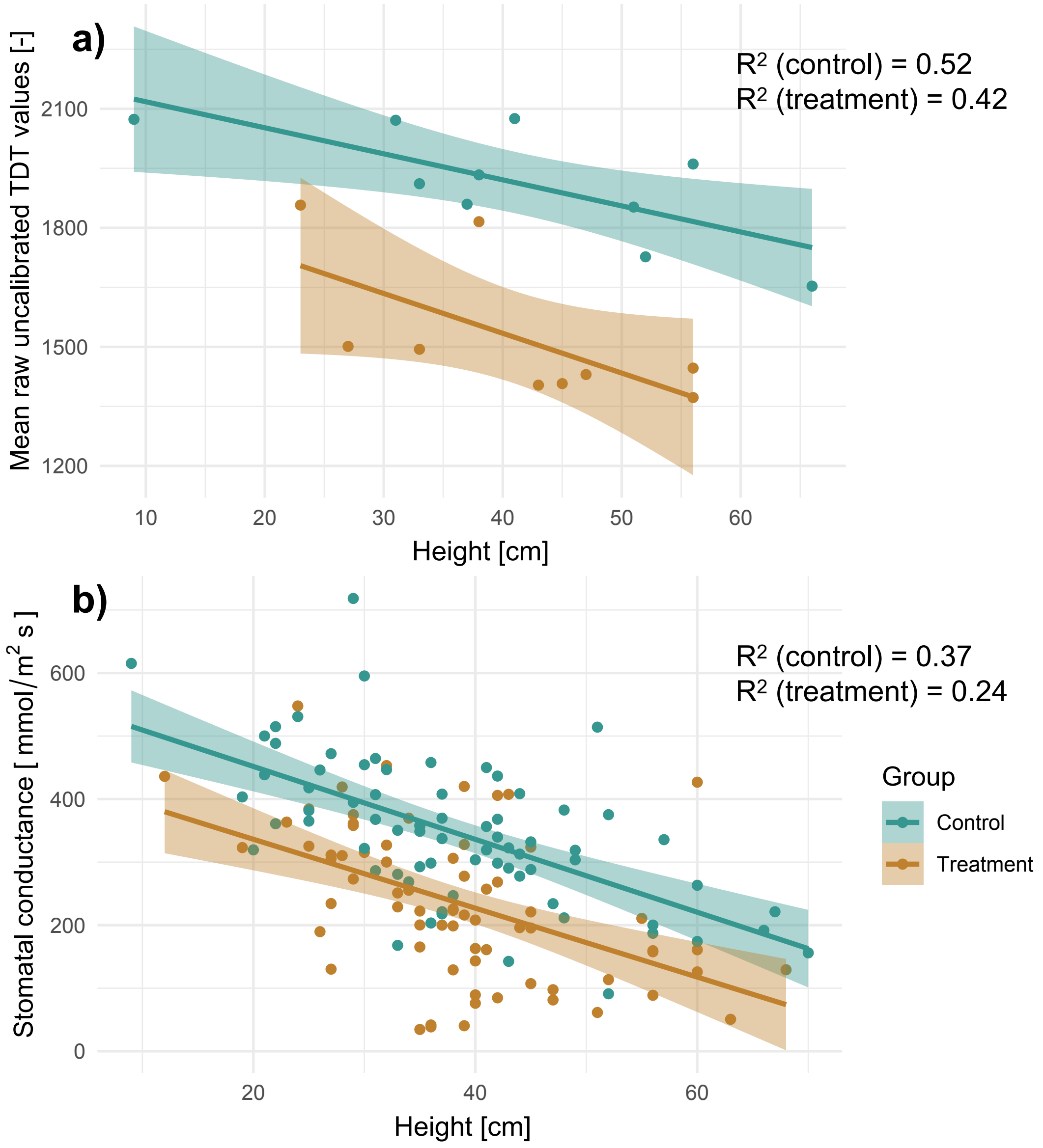


**Supplementary Figure S7.** a) Height of the saplings measured before the experimental drought period (18.04.2023) against mean soil moisture values expressed as raw TDT values for treatment and control group (n=20 soil moisture sensors, 10 per treatment). There was a negative association between soil moisture and sapling height in both treatment ($R^2 = 0.42$, $p = 0.059$) and control group ($R^2 = 0.52$, $p < 0.05$). b) Height of the saplings against stomatal conductance measured during the experimental drought. In the drought treatment, stomatal conductance was significantly reduced across all saplings. Furthermore, there was a negative association between stomatal conductance and sapling height for both treatment ($R^2 = 0.24$, $p < 0.05$) and control groups ($R^2 = 0.37$, $p < 0.05$).

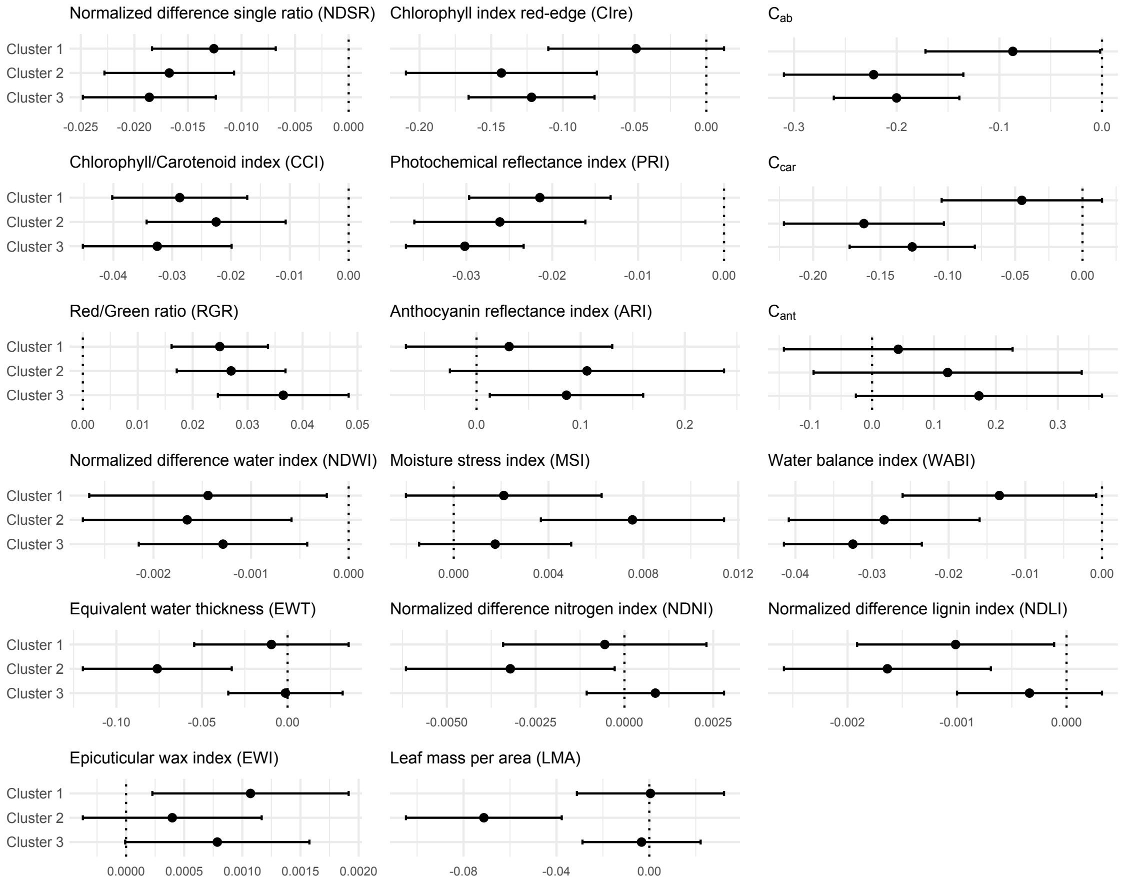


**Supplementary Figure S8.** Log response ratios excluding the ten outliers.

**Supplementary methods: seed collections**

Seeds were collected in 2020 between August 7$^{th}$ and October 1$^{st}$ at 24 different locations in Europe from well-protected sites, thus having a relatively high probability of harbouring populations representing natural migration more than introductions for forestry. Collections were conducted earlier at warmer sites with earlier phenology and later at colder sites with later phenology, as much as possible, and were made opportunistically alongside a field campaign for another project (Czyż et al., 2023) as 2020 was a mast year – thus collection times were not optimized for seed ripeness. European beech was the dominant tree species at all sites. The majority of seed collections were sampled directly from branches of trees located in an area of 10 x 200 meters. However, we also conducted bulk collections from the ground not more than 1 km from a core area of trees used for leaf spectroscopy measurements in Czyż et al. (2023).

Seeds were placed in labelled metal mesh envelopes upon collection; however, for the first collections in August, they were placed in labelled plastic bags which were packed upright and handled carefully (closing during transport, opened when possible), and transferred to metal meshes upon arrival at the University of Zurich (UZH). At UZH, seeds in labelled mesh envelopes were placed in racks upright at ca. 30 °angles covering the bottom of plastic boxes with lids and custom gas inlets and outlets, and dried under a gentle nitrogen flow (ca. 0.1-0.2 bar pressure divided to 5 boxes) until the box reached ca. 10% RH at RT as judged by an off-the-shelf ambient RH and temperature sensor. Once dried, the seeds were transferred to labelled paper envelopes and preserved in sealed plastic bags at 4° C to 5° C (one plastic bag per site) for a maximum of six months, until February 2021. This nitrogen drying and storage procedure was done to reduce or prevent moulding, suffocate and eliminate seed-boring insects (several collections were found to host seed-boring larvae), and thereby better preserve seeds; *F. sylvatica* seeds were previously reported to have longer viability in cold storage (4 °C or -20 °C) with 7.8–11.5%

RH (León-Lobos & Ellis, 2002). The RH of stored seeds was estimated by measuring the RH of these bags at RT and displayed a wide range from 10-40% that was positively correlated with the estimated RH of the seeds upon arrival at UZH between August and October 2020.

An initial test set of seeds from six collections representing six different sites across the sampling range was brought to a nursery on December 2nd, 2020 (47°35'48.9"N, 9°11'40.5"E, Kressibucher AG, Berg Thurgau, Switzerland), where the quality was evaluated by the head of the nursery (M. Kressibucher), and sets of seeds of higher to lower quality were sown in comparison to the nursery's own fresh (not dried) September 2020 collection. Germination of our tested collections was slower and with lower success rates as expected, with seeds collected in September rated as higher quality by the nursery based on size, estimated weight, and appearance. We subsequently brought 94 seed collections representing all 24 sites and, within sites, preferring collections with larger numbers of higher-quality seeds, to the same nursery on February 25th, 2021. There, they were imbibed in water for 24 hours, and then sown in 5 x 5 x 20 cm pots in flats (max 4 seeds/pot) with mixed humus and wood chips and no additional fertilizers. Seeds rated as higher quality by the nursery and thus more likely to germinate were preferentially sown within each collection. Individual seed collections were kept separate and labelled throughout. A wire grid was installed over the pots and the flats were put outside and exposed to the environment. Damage to the dicotyledons was mitigated by bringing them inside a closed shelter when overnight temperatures fell below freezing. Seedlings germinated by July 2021, with seedlings emerging for 43 of the 94 collections and an overall seedling emergence rate of 3.5% (7.9% for the 43 collections in which seedlings emerged). A summary of the collections which yielded seedlings for the common garden is provided in Table ??.

In November 2021, saplings were transported to the Hauenstein garden center in Rafz, Zurich, Switzerland (47°36'36"N, 8°32'35"E). The plants were transplanted into tall 4 L pots filled with an equally proportioned mixture of peat-free potting soil and low-organic matter sandy soil. Each pot was individually labeled with two identical labels corresponding to the collection ID and a unique seedling number; one label was buried in the soil, and the other placed visibly in the pot beside the sapling. The potting soil was enriched with Osmocote Exact Standard NPK (Mg) 12-14M (ICL Group, Tel Aviv-Jaffa, Israel) with micro-nutrients as fertilizer, containing 15% total nitrogen (N), 8% phosphorus pentoxide ($P_2O_5$), 11% potassium oxide ($K_2O$) and 2% total magnesium oxide (MgO). During the winter, the saplings were in a winter tunnel, where temperatures were maintained at circa 7° C. The saplings were watered by an overhead irrigation system as needed. Furthermore, Multikraft BB Start (Multikraft, Pichl/Wels, Austria) was used to treat the soil with ectomycorrhizal fungi. Once it was determined that the saplings were no longer at risk of frost damage, the pots were moved outside the winter tunnel (12 May 2022). We controlled aphid infestation by repeatedly spraying NeemAzal-T/S (3ml/L in water, Andermatt Biogarten, Grossdietwil, Switzerland) directly onto the leaves.

### Supplementary methods: validation of the PROSPECT-D inversion

To assess the results of PROSPECT-D model inversion, we conducted a validation process with a dataset based on an established pigment extraction protocol by Petibon and Wiesenberg (2022). They developed a validated methodology for sequential extraction and subsequent high-performance liquid chromatography analysis (HPLC) of pigments in *F. sylvatica* among other European deciduous species. In this supplementary section, we detail the approach to validate the PROSPECT-D output for both optical subdomains and whole spectra for each leaf trait and assess their accuracy.

#### *Sample selection for pigment extraction*

We collected leaves (n=56) across the control and treatment group in June before and during the drought simulation, from which we used a subset (n=20) for the pigment extraction. The leaf samples were stored on dry ice in non-transparent plastic bags immediately after their spectral measurement and transported to a deep-freezer at -80° C to prevent degradation of the pigments.

To select a suitable range of samples for validation, we first calculated the three indices RGR, NDSR and PRI, which are proxies for relative anthocyanin, chlorophyll, and carotenoid content, respectively, for all the collected samples. Secondly, we clustered the calculated indices based on k-means in python (v3.9.7) from the *sklearn* library (v1.3.2) from Pedregosa et al. (2012). We took a total of 20 samples representing the full range of the pigments for the subsequent extraction.

To estimate pigment concentrations, leaf mass per area is needed. Therefore, we measured the fresh weight of the selected leaves for $fw$ using a semi-micro-balance (MSU125P, Sartorius, Göttingen, Germany, 0.015 mg repeatability) and area $A_{\mathrm{leaf}}$ using the point tool and a known distance in the image analysis program ImageJ (v1.38) (Schneider et al., 2012). The leaves were then freeze-dried overnight using a lyophilizer Alpha 2-4 LD plus (Martin Christ, Osterode am Harz, Germany) and weighed again on the same balance for their dry weight $dw$.

##### *0.0.1 Sampling leaf functional traits*

To validate PROSPECT-D modelled LMA and EWT, in addition to the selected samples for pigment extraction, we took additional samples (n=56) for their $fw$ and $A_{\text{leaf}}$. These fresh leaves were weighed in the field and then stored in paper bags after taking photographs and put into an oven at 70° C for 72 h, after which we weighed the dry leaves for $dw$. $A_{\text{leaf}}$ was estimated based on the photographs using the point tool and a size standard (ruler) in ImageJ (v1.38) (Schneider et al., 2012).

To calculate LMA, we applied the following equation adapted from Féret et al. (2019), where $dw$ indicates oven-dry weight and $A_{\text{leaf}}$ the one-sided leaf area. The result is specified in $mg.cm^{-2}$:

$$\text{leaf mass per area} = \frac{dw}{A_{\text{leaf}}} \tag{1}$$

We calculated EWT specified in $mg.cm^{-2}$ using the following equation by Féret et al. (2019), where $fw$ refers to fresh leaf weight:

$$\text{equivalent water thickness} = \frac{fw - dw}{A_{\text{leaf}}} \tag{2}$$

**Sequential pigment extraction**

For the extraction we used the following high purity solvents: Acetone ($\geq$ 99.9%, UltraGrade Carl Roth, Karlsruhe, Germany), n-hexane ($\geq$ 95%, UltraGrade Carl Roth,

Karlsruhe, Germany) and Isopropanol ($\geq$ 99.9%, HPLC Grade, Sigma-Aldrich, St.Louis, USA). Ultrapure water was taken from a Q-POD MilliQ A10 system (Merck, Darmstadt, Germany).

All procedures were conducted under subdued lighting to avoid pigment degradation by light exposure. We prepared the dried leaf samples by carefully removing the mid-vein and manually grounding the leaf in liquid nitrogen to homogenized leaf powder with a mortar and a pestle. We then filled each amber glass vials with 17-22 mg of ground leaf powder. The vials and the solvents were stored in the freezer at -20° C for later use.

We performed a sequential extraction with the following solvents in this order:

1. acetone:water (85:15, $v/v$)
2. acetone
3. isopropanol:n-hexane (2:1, $v/v$)

We added $\approx$0.8 ml of cooled solvent to the leaf powder vial and then agitated the solution with a vortex stirrer. We left the solution to settle before filtering it on a 1 $\mu m$ glass-fiber filter (Macherey-Nagel, Dueren, Germany) and transferring it to a 10 ml amber vial, which was kept cool on a plate-cooling block. We repeated this procedure three times for each solvent. We then concentrated the eluate, first under a gentle stream of nitrogen (Sample concentrator SBHconc/1, Stuart, Fairham, UK) and secondly with a concentrator plus centrifuge (Vaudaux-Eppendorf, Basel, Switzerland) under vacuum. The extracts were then transferred to amber 1.5 ml HPLC vials with inserts, evaporated to dryness and stored at -20° C until HPLC analysis.

**HPLC analysis**

We used the following reagents for the preparation of the eluents: Methanol (99.9%, gradient grade, Carl Roth, Karlsruhe, Germany), ethyl acetate ($\geq$ 99.8%, LicChrosolv,

Merck, Darmstadt, Germany) and formic acid ($\geq$ 98%, Sigma-Aldrich, Buchs, Switzerland). Ultrapure water was taken from a Q-POD MilliQ A10 system (Merck, Darmstadt, Germany).

The chromatographic separation was performed on an Agilent 1290 Infinity UHPLC system (Agilent Technologies, Santa Clara, USA) with integrated binary pump (Agilent G4220A) and autosampler (Agilent G4226A) with tray cooler kept at 4° C. The column oven was equipped with a InfinityLab Poroshell 120$SB{-}C_{18}$ (4.6 x 150 mm, particle size of 2.7 $\mu m$, Agilent) column and kept at 10° C. The detection was done with a diode array detector (DAD) (Agilent VL+G1315C).

For the HPLC analysis, we used the following eluents:

- Eluent A: water:formic acid (500:1, $v/v$, $pH$ 2.5)
- Eluent B: methanol:ethyl acetate (68:32, $v/v$)

Directly before the HPLC measurement, we redissolved the samples in an exact volume of 50 $\mu l$ of isopropanol:n-hexane (2:1, $v/v$) and 250 $\mu l$ of acetone:water (85:15, $v/v$). Injection volume per sample was 15 $\mu l$ and the flow rate for the chromatographic separation was 0.5 ml per minute. The total run time of the chromatographic separation was 44 min. The gradient for eluent A and B can be found in Table S5.

**Identification and quantification of pigment concentration**

Analytical standards were purchased for the following seven pigments (all Sigma Aldrich, St.Louis, USA): Chlorophyll $a$ ($\geq$ 95.0%) and chlorophyll $b$ ($\geq$ 95.0%), $\alpha$-carotene ($\geq$ 97.0%), $\beta$-carotene ($\geq$ 95.0%), lutein ($\geq$ 96.0%), neoxanthin ($\geq$ 97.0%) and zeaxanthin ($\geq$ 96.0%). Seven standard solutions in the concentration range 1 - 200ug/ml were prepared for each compound in ethyl acetate ($\alpha$-carotene) or acetone (other pigments) and analysed by HPLC-DAD. The derived calibration curves were visualized using R (Fig. S8).

We identified the main pigments based on the absorption spectra and retention time (RT) of the analytical standards (Fig. S9). In addition, we identified two peaks with a significant contribution to the total absorbance as pheophytin *a* and pheophytin *b*, which are demetallised chlorophyll derivatives. To this end, we prepared solutions containing pheophytin *a* and *b* by adding formic acid to standard solutions of chlorophyll *a* and *b* and compared the retention times and absorption spectra (Fig. S10). We also compared the spectra of the unknown peaks with pheophytin spectra reported in literature (Petrovic et al., 2012). All other peaks with absorption > 5 mAU were categorised as either derivatives or unknown compounds on the basis of their spectra according to the decision tree provided by Petibon and Wiesenberg (2022).

Compounds for which standard material was purchased were then quantified based on their peak area using the Agilent OpenLAB CDS ChemStation software at a specific wavelength and the calibration curves shown in Fig. S8. As we observed co-elution for $\alpha$-carotene and $\beta$-carotene, we summed the area of both peaks and applied the calibration curve of $\alpha$-carotene. The concentration of each pigment in the original sample ($C_{\text{pigment}}$, specified in $mg.cm^{-2}$) was then calculated according to the following equation where $A_{\text{peak}}$ is the area of the integrated peak, a and b are the regression coefficients of the calibration curve, $m_g$ is the previously determined ground leaf weight (mg), $dw$ is the dry weight of the leaf (g) and $A_{\text{leaf}}$ is the leaf area (g). The factor 0.3 represents the volume of solvent (ml) in which the sample was dissolved prior to the HPLC analysis.

$$C_{\text{pigment}} = \left(\frac{A_{\text{peak}} - b}{a}\right) \times 0.3 \times m_g^{-1} \times dw \times A_{\text{leaf}}^{-1} \quad (3)$$

The calculated concentrations were summarised as $C_{ab}$ (chlorophyll $a$ and $b$, pheophytin $a$ and $b$) and $C_{car}$ ($\alpha$- and $\beta$-carotene, lutein, neoxanthin, zeaxanthin) which were then used to validate leaf traits from the PROSPECT-D inversion.

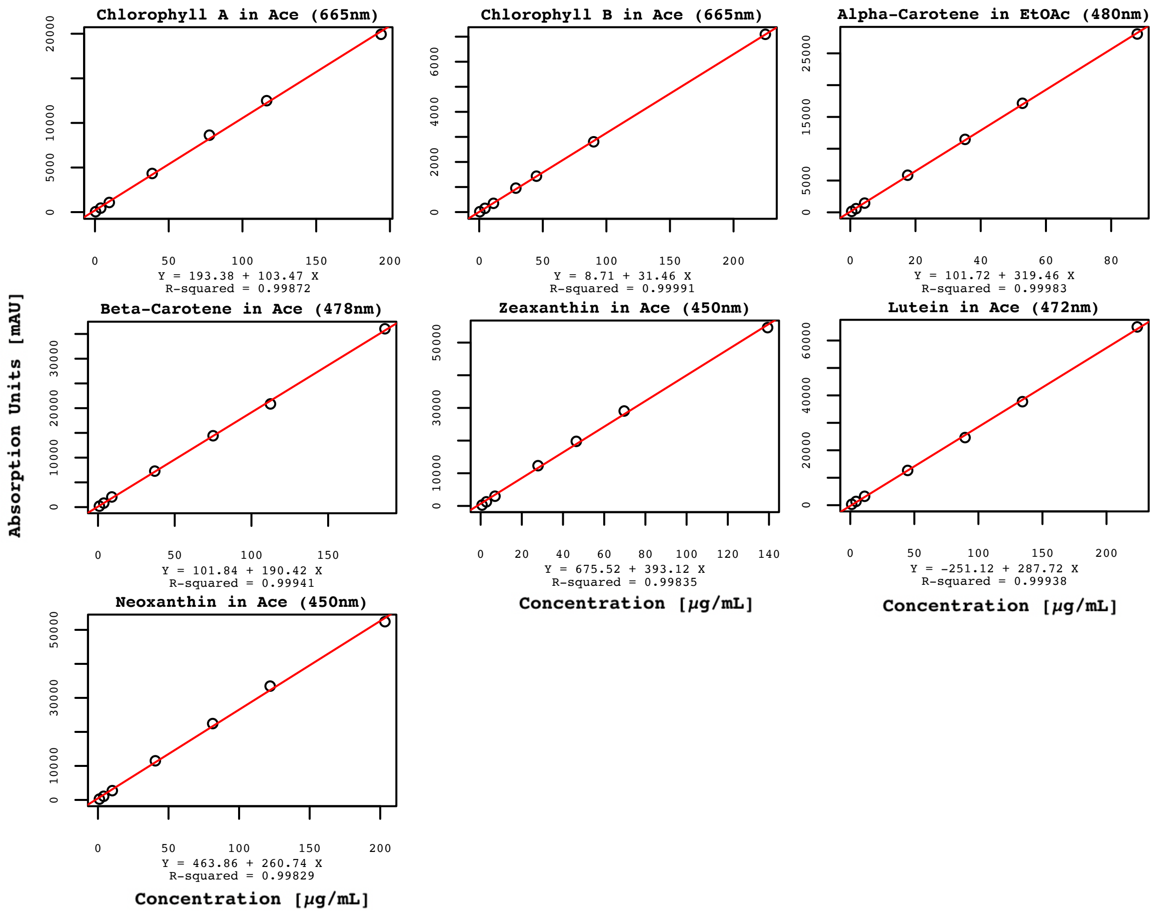


**Supplementary Figure S9.** Calibration curves for the determination of pigment concentration, including absorption wavelength, regression function (lm function in R) and accuracy. Figure created in R v4.3.1

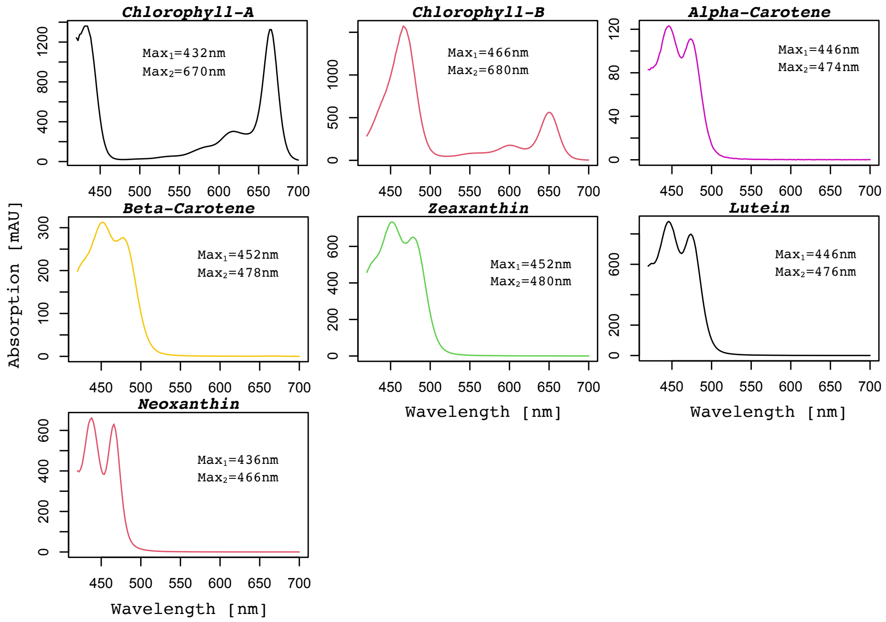


**Supplementary Figure S10.** Pigment standards spectra with absorption maxima used for the identification of pigments. Spectra recorded in ethylacetat (a-carotene) and acetone (other pigments). Figure created in R v4.3.1

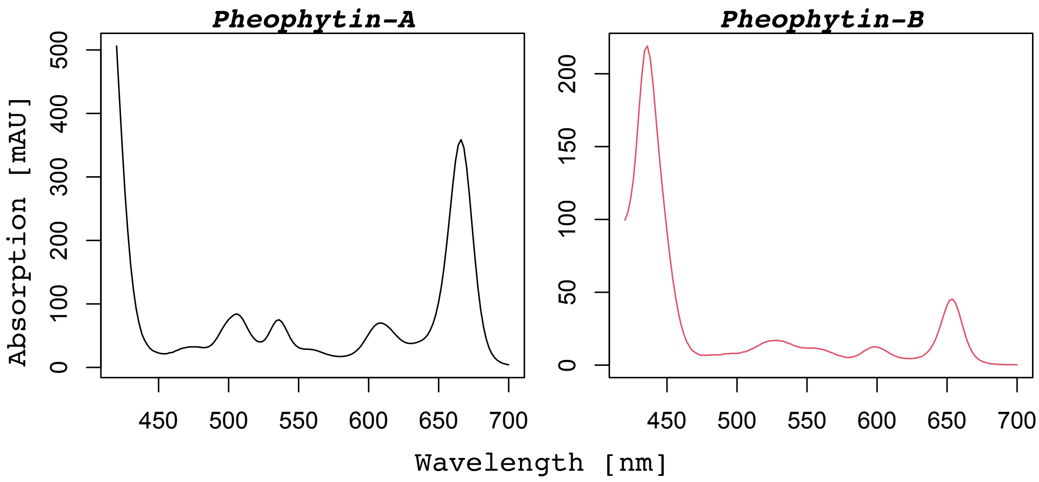


**Supplementary Figure S11.** Spectra of pheophytin a and b sampled from a chromatogram when identifying unknown pigments. Figure created in R v4.3.1.

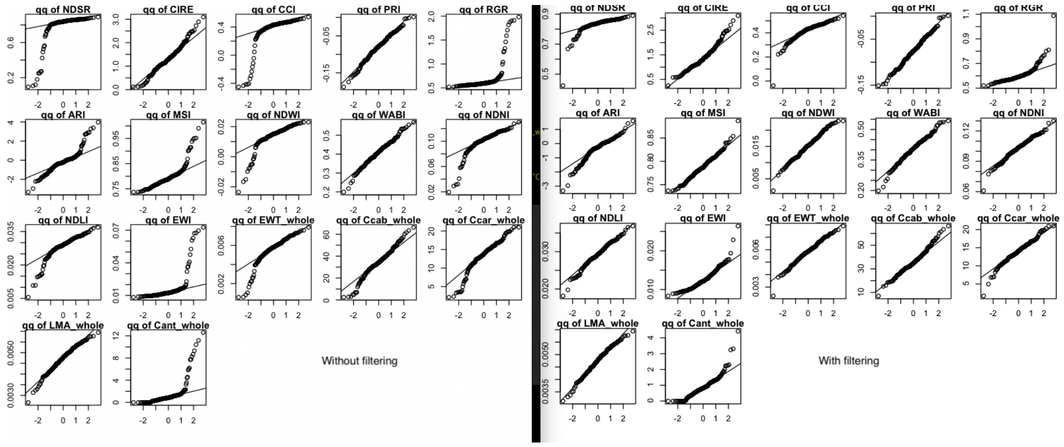


**Supplementary Figure S12.** Q-Q-plots for spectral traits including the 10 outliers (left) vs. without the 10 outliers (right). The ANOVAs presented in Table S6 are based on the data shown on the left, the main analyses based on the filtered data.

**Supplementary Table S2.** Summary of seed collection per site and number of individuals per collection. Collection ID refers to the specific ID for each site and collection. The first three letters refer to the abbreviation of the site. The number refers to the specific collection (if seeds were collected from a top-of-canopy branch, this is a specific mother tree). Bulk collections from the ground are indicated with their exposition. W: West, N: North, E: East, S: South. BES_grd is an exception because the site was so flat that there was no exposition. ROS is another exception, where only ground sampling was conducted, but each collected seed batch received a number. Uneven numbers of individuals per collection are due to uneven germination rates.

| **Site** | **Collection ID** | **No of individuals** |
|---|---|---|
| BES | BES_1 | 7 |
| | BES_2 | 1 |
| | BES_8 | 1 |
| | BES_grd | 2 |
| CHL | CHL_S | 8 |
| ESM | ESM_15 | 5 |
| | ESM_E | 6 |
| ESP | ESP_4 | 11 |
| | ESP_5 | 2 |
| | ESP_8 | 7 |
| | ESP_WNW | 11 |
| FRB | FRB_1 | 10 |
| | FRB_2 | 5 |
| | FRB_10 | 2 |
| | FRB_N | 7 |
| FRM | FRM_ESE | 6 |
| HBV | HBV_N | 2 |
| | HBV_S | 7 |
| HRP | HRP_1 | 7 |
| | HRP_6 | 1 |
| | HRP_9 | 1 |
| | HRP_NW | 3 |
| | HRP_SSW | 6 |
| ITB | ITB_6 | 1 |
| | ITB_7 | 1 |
| ITE | ITE_N | 2 |
| ITM | ITM_6 | 1 |
| | ITM_SSW | 1 |
| PLB | PLB_4 | 2 |
| PLW | PLW_1 | 10 |
| | PLW_6 | 1 |
| | PLW_10 | 8 |
| ROS | ROS_1 | 6 |
| | ROS_6 | 2 |
| RSF | RSF_ENE | 7 |
| | RSF_WNW | 2 |
| SLK | SLK_1 | 7 |
| | SLK_3 | 5 |
| | SLK_4 | 2 |
| | SLK_N | 3 |
| | SLK_SSW | 1 |
| **TOTAL** | | **180** |

**Supplementary Table S3.** Timeline of irrigation during the drought experiment. Ad-hoc irrigation refers to watering saplings that exhibited signs of severe wilting. This was done to prevent mortality of those individuals. When no amount in mL is indicated, the saplings were watered *ad libidum.*

| Date | Action |
|---|---|
| 09.06.23 | Start of drought treatment |
| 10.06.23 | Irrigation of control saplings |
| 11.06.23 | Irrigation of control saplings |
| 13.06.23 | Irrigation of all saplings: 600 mL |
| 14.06.23 | Irrigation of control saplings, |
| | Ad-hoc irrigation of E1 (CHL S 10): 600 mL |
| 15.06.23 | Irrigation of control saplings |
| 16.06.23 | Ad-hoc irrigation of the following drought-treated saplings: |
| | E1 (CHL S 10): 600 mL, |
| | C18 (PLW 01 6): 600 mL, |
| | C11 (FRM ESE 4): 300 mL, |
| | E5 (HBV S 6): 300 mL |
| 17.06.23 | Irrigation of control saplings, |
| | Ad-hoc irrigation of the following drought-treated saplings: |
| | A1 (BES 1 2): 600 mL, |
| | A5 (ROS 1 3): 600 mL, |
| | E5 (HBV S 6): 300 mL, |
| | E1 (CHL S 10): 300 mL, |
| | C17 (FRB 2 4): 300 mL, |
| | C11 (FRM ESE 4): 300 mL |
| 18.06.23 | Irrigation of all saplings: 600 mL |
| 19.06.23 | Irrigation of control saplings |
| 20.06.23 | Irrigation of control saplings |
| 22.06.23 | Irrigation of control saplings |
| 24.06.23 | Irrigation of all saplings |

**Supplementary Table S4.** Instrument configurations for ASD FieldSpec 4 during spectra acquisition.

| Type | Configuration | Type | Configuration |
|---|---|---|---|
| Foreoptic | Bare Fiber | Integration Time | 8.5 ms |
| Spectrum | 10 | SWIR1 Gain | 500 |
| Dark Current | 100 | SWIR1 Offset | 2048 |
| White Reference | 10 | SWIR2 Gain | 500 |
| Scan Type | AB Even | SWIR2 Offset | 2048 |

**Supplementary Table S5.** Gradient used for HPLC analysis.

| Run time (min) | Eluent A (%) | Eluent B (%) |
|---|---|---|
| 0 | 20 | 80 |
| 15 | 10 | 90 |
| 20 | 0 | 100 |
| 41 | 20 | 80 |
| 44 | 20 | 80 |

**Supplementary Table S6.** ANOVA table including the 10 outliers for the linear models with spectral leaf traits as dependent observations and sapling height, treatment, genetic cluster, and treatment $\times$ genetic cluster interaction as explanatory factors. These models are presented here only for comparison because they violate model assumptions and are, therefore, not valid (see also Q-Q-plots in Figure S12). This illustrates the approaches we tried to incorporate the outliers in the main analysis. $p \leq 0.001\,('***')$, $p \leq 0.01\,('**')$, $p \leq 0.05\,('*')$.

| **Variable** | **Factor** | **DF** | **F value** | **P value** |
|---|---|---|---|---|
| NDSR | Height | 1 | 8.2468 | 0.004593 ** |
| | Treatment | 1 | 23.1697 | 3.218e-06 *** |
| | Cluster | 2 | 1.0318 | 0.358553 |
| | Treatment x Cluster | 2 | 1.2606 | 0.286086 |
| | Residuals | 173 | | |
| CIRE | Height | 1 | 17.4915 | 4.579e-05 ** |
| | Treatment | 1 | 27.0693 | 5.507e-07 *** |
| | Cluster | 2 | 0.0217 | 0.9786 |
| | Treatment x Cluster | 2 | 1.2723 | 0.2828 |
| | Residuals | 173 | | |
| CCI | Height | 1 | 6.2774 | 0.01315 * |
| | Treatment | 1 | 23.4178 | 2.872e-06 *** |
| | Cluster | 2 | 1.1795 | 0.30988 |
| | Treatment x Cluster | 2 | 1.1323 | 0.32468 |
| | Residuals | 173 | | |
| PRI | Height | 1 | 17.9106 | 3.750e-05 *** |
| | Treatment | 1 | 49.4107 | 4.599e-11 *** |
| | Cluster | 2 | 0.5747 | 0.5640 |
| | Treatment x Cluster | 2 | 0.9724 | 0.3802 |
| | Residuals | 173 | | |
| RGR | Height | 1 | 9.9159 | 0.001931 ** |
| | Treatment | 1 | 21.8024 | 6.04e-06 *** |
| | Cluster | 2 | 1.1241 | 0.327299 |
| | Treatment x Cluster | 2 | 1.3025 | 0.274495 |
| | Residuals | 173 | | |
| ARI | Height | 1 | 22.7064 | 3.981e-06 *** |
| | Treatment | 1 | 16.4337 | 7.604e-05 *** |
| | Cluster | 2 | 0.5891 | 0.5560 |
| | Treatment x Cluster | 2 | 0.9715 | 0.3806 |
| | Residuals | 173 | | |
| MSI | Height | 1 | 1.6108 | 0.2060819 |
| | Treatment | 1 | 14.8100 | 0.0001671 *** |
| | Cluster | 2 | 0.9737 | 0.3797449 |
| | Treatment x Cluster | 2 | 0.8242 | 0.4403154 |
| | Residuals | 173 | | |
| NDWI | Height | 1 | 2.2486 | 0.1356 |
| | Treatment | 1 | 18.3898 | 2.986e-05 * |
| | Cluster | 2 | 0.4094 | 0.6647 |
| | Treatment x Cluster | 2 | 0.5306 | 0.5892 |
| | Residuals | 173 | | |
| WABI | Height | 1 | 0.2626 | 0.609018 |
| | Treatment | 1 | 7.1423 | 0.008249 ** |
| | Cluster | 2 | 1.0703 | 0.345181 |
| | Treatment x Cluster | 2 | 0.1262 | 0.881550 |
| | Residuals | 173 | | |
| NDNI | Height | 1 | 2.0242 | 0.156608 |
| | Treatment | 1 | 9.2463 | 0.002727 ** |
| | Cluster | 2 | 1.8683 | 0.157490 |
| | Treatment x Cluster | 2 | 0.5076 | 0.602824 |
| | Residuals | 173 | | |
| NDLI | Height | 1 | 6.6164 | 0.0109440 |
| | Treatment | 1 | 15.5805 | 0.0001148 *** |
| | Cluster | 2 | 2.2456 | 0.1089461 |
| | Treatment x Cluster | 2 | 0.2554 | 0.7748615 |
| | Residuals | 173 | | |
| EWI | Height | 1 | 2.7964 | 0.0962818 |
| | Treatment | 1 | 14.1466 | 0.0002312 *** |
| | Cluster | 2 | 0.8660 | 0.4224438 |
| | Treatment x Cluster | 2 | 0.7414 | 0.4779563 |
| | Residuals | 173 | | |
| $C_{ab}$ | Height | 1 | 16.4336 | 7.604e-05 *** |
| | Treatment | 1 | 36.1301 | 1.065e-08 *** |
| | Cluster | 2 | 0.1108 | 0.895 |
| | Treatment x Cluster | 2 | 1.6581 | 0.1935 |
| | Residuals | 173 | | |
| $C_{car}$ | Height | 1 | 6.8164 | 0.009825 ** |
| | Treatment | 1 | 29.6606 | 1.744e-07 *** |
| | Cluster | 2 | 0.1918 | 0.825611 |
| | Treatment x Cluster | 2 | 1.7771 | 0.172206 |
| | Residuals | 173 | | |
| $C_{ant}$ | Height | 1 | 12.8128 | 0.0004469 *** |
| | Treatment | 1 | 12.9743 | 0.0004125 *** |
| | Cluster | 2 | 1.0264 | 0.3604797 |
| | Treatment x Cluster | 2 | 1.1885 | 0.3071447 |
| | Residuals | 173 | | |
| EWT | Height | 1 | 2.4460 | 0.1196485 |
| | Treatment | 1 | 11.6029 | 0.0008188 *** |
| | Cluster | 2 | 1.1637 | 0.3147635 |
| | Treatment x Cluster | 2 | 0.8946 | 0.4106566 |
| | Residuals | 173 | | |
| LMA | Height | 1 | 0.0111 | 0.91635 |
| | Treatment | 1 | 3.6394 | 0.05808 |
| | Cluster | 2 | 0.8831 | 0.41537 |
| | Treatment x Cluster | 2 | 1.1645 | 0.31451 |
| | Residuals | 173 | | |
