## Supplemental Table S1 for "Leaf spectroscopy reveals drought response variation in *Fagus sylvatica* saplings from across the species’ range"

**Table S1.** Overview of all 24 sampling locations across Europe. ID refers to the specific ID for each site and collection. The first three letters refer to the abbreviation of the site. The number refers to the specific collection (if seeds were collected from a top-of-canopy branch, this is a specific mother tree). Bulk collections from the ground are indicated with their exposition. W: West, N: North, E: East, S: South. BES is an exception because it was so flat that there was no exposition. ROS is another exception, where only ground sampling was conducted. Transect means that we selected two transects of five trees each, in parallel to each other, and the trees within transects were ca. 50 m apart. Normally the transects started near the top of a hill and went straight down the hill. Collections that are part of the present study are highlighted in bold.

| **Country** | **Locality** | **Date** | **Exposition** | **Substrate** | **Geology** | **ID** | **Longitude** | **Latitude** | **Elevation** | **Transect** | **Seeds_long** | **Seeds_lat** | **Seeds provenance** | **Comments** |
|---|---|---|---|---|---|---|---|---|---|---|---|---|---|---|
| Italy | Bracciano | 11.08.2020 | E | igneous | Vulcanic | ITB_01 | 12.15492814 | 42.17390453 | 605.2874654 | 1 | 12.15492814 | 42.17390453 | branch | ok, big seeds |
| Italy | Bracciano | 11.08.2020 | E | igneous | Vulcanic | ITB_02 | 12.15541119 | 42.17373173 | 599.9394719 | 1 | 12.15541119 | 42.17373173 | branch | 2 bags, ok, big seeds |
| Italy | Bracciano | 11.08.2020 | E | igneous | Vulcanic | ITB_03 | 12.15604266 | 42.17375268 | 583.7859593 | 1 | 12.15604266 | 42.17375268 | branch | ok |
| Italy | Bracciano | 11.08.2020 | E | igneous | Vulcanic | ITB_04 | 12.15644968 | 42.17355018 | 569.7990876 | 1 | na | na | na | no seeds |
| Italy | Bracciano | 11.08.2020 | E | igneous | Vulcanic | ITB_05 | 12.15695463 | 42.17328708 | 553.5141462 | 1 | 12.15695463 | 42.17328708 | branch | ok |
| **Italy** | **Bracciano** | **11.08.2020** | **E** | **igneous** | **Vulcanic** | **ITB_06** | **12.15731961** | **42.17368886** | **561.5835** | **2** | **12.15731961** | **42.17368886** | **branch** | ok |
| Italy | Bracciano | 21.08.2020 | E | igneous | Vulcanic | ITB_06_b | 12.15731961 | 42.17368886 | 561.5834972 | 2 | na | na | na | mixed with ITB_06 |
| **Italy** | **Bracciano** | **21.08.2020** | **E** | **igneous** | **Vulcanic** | **ITB_07** | **12.15686013** | **42.17386617** | **582.78728** | **2** | **12.15686013** | **42.17386617** | **branch** | Ok 2 bags |
| Italy | Bracciano | 21.08.2020 | E | igneous | Vulcanic | ITB_08 | 12.15648486 | 42.17411176 | 580.5586719 | 2 | 12.15648486 | 42.17411176 | branch | ok (big seeds) |
| Italy | Bracciano | 21.08.2020 | E | igneous | Vulcanic | ITB_09 | 12.15600583 | 42.17447731 | 599.2955041 | 2 | 12.15600583 | 42.17447731 | branch | ok |
| Italy | Bracciano | 21.08.2020 | E | igneous | Vulcanic | ITB_10 | 12.15528532 | 42.17455818 | 600.3009508 | 2 | 12.15528532 | 42.17455818 | branch | removed worms and perfored seeds |
| Italy | Bracciano | 21.08.2020 | N | igneous | Vulcanic | ITB_11 | 12.15532432 | 42.17537485 | 594.4725105 | 3 | na | na | na | |
| Italy | Bracciano | 21.08.2020 | N | igneous | Vulcanic | ITB_12 | 12.1555291 | 42.17584766 | 593.6521125 | 3 | na | na | na | |
| Italy | Bracciano | 21.08.2020 | N | igneous | Vulcanic | ITB_13 | 12.15573793 | 42.17633867 | 569.9936766 | 3 | na | na | na | |
| Italy | Bracciano | 21.08.2020 | E | igneous | Vulcanic | ITB_E | na | na | na | 1,2 | 12.15492814 | 42.17390453 | ground | few seeds on the ground, probably sterile |

| | | | | | | | | | | | | | | |
|---|---|---|---|---|---|---|---|---|---|---|---|---|---|---|
| Italy | Bracciano | 21.08.2020 | N | igneous | Vulcanic | ITB_N | na | na | na | 3 | 12.15573793 | 42.17633867 | ground | ok |
| Italy | Monte_Vulture | 13.08.2020 | SW | igneous | Vulcanic | ITV_01 | 15.6318338 | 40.94931997 | 1188.509402 | 1 | 15.6318338 | 40.94931997 | branch | ok, molt before N2 drying, slight EtOH wash, looks ok after drying |
| Italy | Monte_Vulture | 13.08.2020 | SW | igneous | Vulcanic | ITV_02 | 15.63176112 | 40.94900288 | 1168.326493 | 1 | 15.63176112 | 40.94900288 | branch | ok, molt before N2 drying, slight EtOH wash, looks ok after drying |
| Italy | Monte_Vulture | 13.08.2020 | SW | igneous | Vulcanic | ITV_03 | 15.63156098 | 40.94859742 | 1140.273498 | 1 | 15.63156098 | 40.94859742 | branch | ok, molt before N2 drying, slight EtOH wash, looks ok after drying |
| Italy | Monte_Vulture | 13.08.2020 | SW | igneous | Vulcanic | ITV_04 | 15.63109713 | 40.94823472 | 1122.126009 | 1 | 15.63109713 | 40.94823472 | branch | ok, molt before N2 drying, slight EtOH wash, looks ok after drying |
| Italy | Monte_Vulture | 13.08.2020 | SW | igneous | Vulcanic | ITV_05 | 15.63093358 | 40.9479578 | 1102.356274 | 1 | 15.63093358 | 40.9479578 | branch | ok, molt before N2 drying, slight EtOH wash, looks ok after drying |
| Italy | Monte_Vulture | 13.08.2020 | SW | igneous | Vulcanic | ITV_06 | 15.63049608 | 40.9481467 | 1087.424622 | 2 | 15.63049608 | 40.9481467 | branch | ok, molt before N2 drying, slight EtOH wash, looks ok after drying |
| Italy | Monte_Vulture | 13.08.2020 | SW | igneous | Vulcanic | ITV_07 | 15.63056491 | 40.94850306 | 1105.632427 | 2 | 15.63056491 | 40.94850306 | branch | ok |
| Italy | Monte_Vulture | 13.08.2020 | SW | igneous | Vulcanic | ITV_08 | 15.63076344 | 40.94876348 | 1128.523265 | 2 | 15.63076344 | 40.94876348 | branch | dried, slightly molted inside |
| Italy | Monte_Vulture | 13.08.2020 | SW | igneous | Vulcanic | ITV_09 | 15.63124338 | 40.94907127 | 1152.693826 | 2 | 15.63124338 | 40.94907127 | branch | ok, molt before N2 drying, slight EtOH wash, looks ok after drying |
| Italy | Monte_Vulture | 13.08.2020 | SW | igneous | Vulcanic | ITV_10 | 15.63139589 | 40.9494184 | 1173.536309 | 2 | na | na | na | no seeds |
| Italy | Monte_Vulture | 13.08.2020 | SW | igneous | Vulcanic | ITV_SW | na | na | na | 1,2 | 15.6318338 | 40.94931997 | ground | molt before N2 drying, slight EtOH wash, looks ok after drying |
| Italy | Monte_Pierfaone | 14.08.2020 | E | sedimentary | Limestone | ITF_01 | 15.752594 | 40.50605401 | 1763.121942 | 1 | 15.752594 | 40.50605401 | branch | molt before N2 drying, slight EtOH wash, looks ok after drying |
| Italy | Monte_Pierfaone | 14.08.2020 | E | sedimentary | Limestone | ITF_02 | 15.75294968 | 40.50599479 | 1752.613089 | 1 | 15.75294968 | 40.50599479 | branch | no seeds |
| Italy | Monte_Pierfaone | 14.08.2020 | E | sedimentary | Limestone | ITF_03 | 15.75323036 | 40.5059187 | 1741.731355 | 1 | 15.75323036 | 40.5059187 | ground | lots of seeds!, molt before N2 drying, slight EtOH wash, looks ok after drying |

| | | | | | | | | | | | | | | |
|---|---|---|---|---|---|---|---|---|---|---|---|---|---|---|
| Italy | Monte_Pierfaone | 14.08.2020 | E | sedimentary | Limestone | ITF_04 | 15.75360669 | 40.50589432 | 1727.689657 | 1 | 15.75360669 | 40.50589432 | branch | ok |
| Italy | Monte_Pierfaone | 14.08.2020 | E | sedimentary | Limestone | ITF_05 | 15.7539473 | 40.50577554 | 1713.754214 | 1 | 15.7539473 | 40.50577554 | branch | ok, molt before N2 drying, slight EtOH wash, looks ok after drying |
| Italy | Monte_Pierfaone | 14.08.2020 | E | sedimentary | Limestone | ITF_06 | 15.75395551 | 40.50549512 | 1711.475528 | 2 | 15.75395551 | 40.50549512 | branch | ok |
| Italy | Monte_Pierfaone | 14.08.2020 | E | sedimentary | Limestone | ITF_07 | 15.75361634 | 40.50542736 | 1731.249925 | 2 | 15.75361634 | 40.50542736 | branch | ok |
| Italy | Monte_Pierfaone | 14.08.2020 | E | sedimentary | Limestone | ITF_08 | 15.75340126 | 40.50566795 | 1741.667312 | 2 | 15.75340126 | 40.50566795 | branch | ok |
| Italy | Monte_Pierfaone | 14.08.2020 | E | sedimentary | Limestone | ITF_09 | 15.75310496 | 40.50563622 | 1753.894459 | 2 | 15.75310496 | 40.50563622 | branch | ok, few seeds |
| Italy | Monte_Pierfaone | 14.08.2020 | E | sedimentary | Limestone | ITF_10 | 15.75267489 | 40.50570816 | 1766.330063 | 2 | 15.75267489 | 40.50570816 | branch | ok |
| Italy | Monte_Pierfaone | 14.08.2020 | E | sedimentary | Limestone | ITF_E | na | na | na | 1,2 | na | na | na | |
| Italy | Madonie_Park | 16.08.2020 | S | sedimentary | Limestone | ITM_01 | 14.0420053 | 37.85853487 | 1774.67017 | 1 | 14.0420053 | 37.85853487 | branch | ok, few seeds |
| Italy | Madonie_Park | 16.08.2020 | S | sedimentary | Limestone | ITM_02 | 14.04206055 | 37.85829899 | 1754.442462 | 1 | 14.04206055 | 37.85829899 | branch | ok, seeds small, probably sterile |
| Italy | Madonie_Park | 16.08.2020 | S | sedimentary | Limestone | ITM_03 | 14.04183043 | 37.8577452 | 1696.556724 | 1 | 14.04183043 | 37.8577452 | branch | ok |
| Italy | Madonie_Park | 16.08.2020 | S | sedimentary | Limestone | ITM_04 | 14.04185205 | 37.85735751 | 1665.466805 | 1 | 14.04185205 | 37.85735751 | branch | ok |
| Italy | Madonie_Park | 16.08.2020 | S | sedimentary | Limestone | ITM_05 | 14.04158261 | 37.85684177 | 1642.51447 | 1 | 14.04158261 | 37.85684177 | branch | ok, few seeds (5) |
| **Italy** | **Madonie_Park** | **16.08.2020** | **S** | **sedimentary** | **Limestone** | **ITM_06** | **14.04091599** | **37.85692687** | **1654.5689** | **2** | **14.04091599** | **37.85692687** | **branch** | **ok** |
| Italy | Madonie_Park | 16.08.2020 | S | sedimentary | Limestone | ITM_07 | 14.04119879 | 37.85733116 | 1685.49996 | 2 | 14.04119879 | 37.85733116 | branch | ok |
| Italy | Madonie_Park | 16.08.2020 | S | sedimentary | Limestone | ITM_08 | 14.04137 | 37.85781334 | 1707.904344 | 2 | 14.04137 | 37.85781334 | branch | ok |
| Italy | Madonie_Park | 16.08.2020 | S | sedimentary | Limestone | ITM_09 | 14.04165567 | 37.85834292 | 1753.064007 | 2 | 14.04165567 | 37.85834292 | branch | ok |
| Italy | Madonie_Park | 16.08.2020 | S | sedimentary | Limestone | ITM_10 | 14.04165452 | 37.85857566 | 1775.380355 | 2 | 14.04165452 | 37.85857566 | branch | ok |
| Italy | Madonie_Park | 16.08.2020 | N | sedimentary | Limestone | ITM_11 | 14.03964185 | 37.85162929 | 1720.852091 | 3 | 14.03964185 | 37.85162929 | branch | ok, seeds very small |
| Italy | Madonie_Park | 16.08.2020 | N | sedimentary | Limestone | ITM_12 | 14.03941431 | 37.85189959 | 1703.002317 | 3 | na | na | na | no seeds |

| | | | | | | | | | | | | | | |
|---|---|---|---|---|---|---|---|---|---|---|---|---|---|---|
| Italy | Madonie_Park | 16.08.2020 | N | sedimentary | Limestone | ITM_13 | 14.03937156 | 37.85237362 | 1679.329646 | 3 | na | na | na | no seeds |
| Italy | Madonie_Park | 16.08.2020 | N | sedimentary | Limestone | ITM_14 | 14.03926096 | 37.85301995 | 1634.98262 | 3 | na | na | na | no seeds |
| Italy | Madonie_Park | 16.08.2020 | N | sedimentary | Limestone | ITM_15 | 14.03912813 | 37.85378796 | 1589.420353 | 3 | na | na | na | no seeds |
| Italy | Madonie_Park | 16.08.2020 | N | sedimentary | Limestone | ITM_N | | na | na | 3 | 14.03912813 | 37.85378796 | ground | Ok, 2 bags, 1:24, 2:147 |
| Italy | Madonie_Park | 16.08.2020 | S | sedimentary | Limestone | ITM_S | | na | na | 1,2 | 14.0420053 | 37.85853487 | ground | Ok 2 bags, 1:180, 2:74 |
| Italy | Etna | 17.08.2020 | SSW | igneous | Vulcanic | ITE_01 | 15.04391952 | 37.70975374 | 1776.686524 | 1 | na | na | na | no seeds |
| Italy | Etna | 17.08.2020 | SSW | igneous | Vulcanic | ITE_02 | 15.04386 | 37.70945 | 1742.502 | 1 | na | na | na | no seeds |
| Italy | Etna | 17.08.2020 | SSW | igneous | Vulcanic | ITE_03 | 15.04411173 | 37.70906338 | 1713.989925 | 1 | 15.04411173 | 37.70906338 | branch | ok |
| Italy | Etna | 17.08.2020 | SSW | igneous | Vulcanic | ITE_04 | 15.04452962 | 37.7083089 | 1681.197843 | 1 | na | na | na | no seeds |
| Italy | Etna | 17.08.2020 | SSW | igneous | Vulcanic | ITE_05 | 15.04475287 | 37.70770489 | 1651.760876 | 1 | na | na | na | no seeds |
| Italy | Etna | 17.08.2020 | SSW | igneous | Vulcanic | ITE_06 | 15.04535067 | 37.70773567 | 1664.887808 | 2 | 15.04535067 | 37.70773567 | branch | ok |
| Italy | Etna | 17.08.2020 | SSW | igneous | Vulcanic | ITE_07 | 15.04551924 | 37.70844587 | 1714.785074 | 2 | 15.04551924 | 37.70844587 | branch | ok |
| Italy | Etna | 17.08.2020 | SSW | igneous | Vulcanic | ITE_08 | 15.04558455 | 37.7088537 | 1736.17355 | 2 | 15.04558455 | 37.7088537 | branch | very small seeds |
| Italy | Etna | 17.08.2020 | SSW | igneous | Vulcanic | ITE_09 | 15.04587706 | 37.70904804 | 1761.749932 | 2 | 15.04587706 | 37.70904804 | branch | ok |
| Italy | Etna | 17.08.2020 | SSW | igneous | Vulcanic | ITE_10 | 15.04598005 | 37.70938723 | 1790.477565 | 2 | 15.04598005 | 37.70938723 | branch | ok, 4 seeds |
| Italy | Etna | 17.08.2020 | N | igneous | Vulcanic | ITE_11 | 15.04697124 | 37.70930558 | 1787.626139 | 2 | 15.04697124 | 37.70930558 | branch | ok |
| Italy | Etna | 17.08.2020 | N | igneous | Vulcanic | ITE_12 | 15.04708746 | 37.70949728 | 1775.792441 | 3 | 15.04708746 | 37.70949728 | branch | ok, small seeds probably sterile |
| Italy | Etna | 17.08.2020 | N | igneous | Vulcanic | ITE_13 | 15.04704834 | 37.70966354 | 1761.097808 | 3 | 15.04704834 | 37.70966354 | na | no seeds |
| Italy | Etna | 17.08.2020 | N | igneous | Vulcanic | ITE_14 | 15.04710868 | 37.70980333 | 1748.505879 | 3 | 15.04710868 | 37.70980333 | na | no seeds |
| Italy | Etna | 17.08.2020 | N | igneous | Vulcanic | ITE_15 | 15.04712616 | 37.71007955 | 1728.714175 | 3 | 15.04712616 | 37.71007955 | na | no seeds |

| | | | | | | | | | | | | | | |
|---|---|---|---|---|---|---|---|---|---|---|---|---|---|---|
| Italy | Etna | 17.08.2020 | S | igneous | Vulcanic | ITE_16_big tree | 15.046667 | 37.705 | 1513 | 3 | 15.046667 | 37.705 | branch | ok |
| Italy | Etna | 17.08.2020 | SSW | igneous | Vulcanic | ITE_SSW | | na | na | 1,2 | 15.04391952 | 37.70975374 | ground | removed dead worms and perfored seeds, ok |
| **Italy** | **Etna** | **17.08.2020** | **N** | **igneous** | **Vulcanic** | **ITE_N** | | **na** | **na** | **3** | **15.04598005** | **37.70938723** | **ground** | **ok** |
| Italy | Pollino | 19.08.2020 | SSE | sedimentary | Limestone | ITP_01 | 16.10892709 | 39.90561078 | 1854.77149 | 1 | 16.10892709 | 39.90561078 | branch | ok, . Seeds seems empty/sterile... |
| Italy | Pollino | 19.08.2020 | SSE | sedimentary | Limestone | ITP_02 | 16.10933653 | 39.90529879 | 1811.082456 | 1 | 16.10933653 | 39.90529879 | branch | ok, . Seeds seems empty/sterile... |
| Italy | Pollino | 19.08.2020 | SSE | sedimentary | Limestone | ITP_03 | 16.10937041 | 39.90495197 | 1796.870765 | 1 | 16.10937041 | 39.90495197 | branch | ok, . Seeds seems empty/sterile... |
| Italy | Pollino | 19.08.2020 | SSE | sedimentary | Limestone | ITP_04 | 16.10973178 | 39.9046188 | 1754.493876 | 1 | 16.10973178 | 39.9046188 | branch | ok, . Seeds seems empty/sterile... |
| Italy | Pollino | 19.08.2020 | SSE | sedimentary | Limestone | ITP_05 | 16.10993086 | 39.90428101 | 1738.56295 | 1 | 16.10993086 | 39.90428101 | branch | ok, . Seeds seems empty/sterile... |
| Italy | Pollino | 19.08.2020 | SSE | sedimentary | Limestone | ITP_06 | 16.11072789 | 39.90444785 | 1746.842366 | 2 | 16.11072789 | 39.90444785 | branch | ok, . Seeds seems empty/sterile... |
| Italy | Pollino | 19.08.2020 | SSE | sedimentary | Limestone | ITP_07 | 16.1103739 | 39.90479061 | 1765.035138 | 2 | 16.1103739 | 39.90479061 | branch | ok, . Seeds seems empty/sterile... |
| Italy | Pollino | 19.08.2020 | SSE | sedimentary | Limestone | ITP_08 | 16.11017805 | 39.905112 | 1801.865942 | 2 | 16.11017805 | 39.905112 | branch | ok, . Seeds seems empty/sterile... |
| Italy | Pollino | 19.08.2020 | SSE | sedimentary | Limestone | ITP_09 | 16.10974045 | 39.90546272 | 1827.181867 | 2 | 16.10974045 | 39.90546272 | branch | ok, . Seeds seems empty/sterile... |
| Italy | Pollino | 19.08.2020 | SSE | sedimentary | Limestone | ITP_10 | 16.109378 | 39.90585892 | 1859.098358 | 2 | 16.109378 | 39.90585892 | branch | ok, . Seeds seems empty/sterile... |
| Italy | Pollino | 19.08.2020 | NW | sedimentary | Limestone | ITP_11 | 16.11518795 | 39.9087096 | 1800.335228 | 3 | na | na | na | no seeds |
| Italy | Pollino | 19.08.2020 | NW | sedimentary | Limestone | ITP_12 | 16.115337 | 39.90910506 | 1766.446752 | 3 | na | na | na | no seeds |
| Italy | Pollino | 19.08.2020 | NW | sedimentary | Limestone | ITP_13 | 16.11509333 | 39.90960356 | 1729.953427 | 3 | na | na | na | no seeds |
| Italy | Pollino | 19.08.2020 | NW | sedimentary | Limestone | ITP_14 | 16.11501292 | 39.91005302 | 1698.478396 | 3 | na | na | na | no seeds |
| Italy | Pollino | 19.08.2020 | NW | sedimentary | Limestone | ITP_15 | 16.11491275 | 39.91050293 | 1665.885949 | 3 | na | na | na | no seeds |
| Italy | Pollino | 19.08.2020 | SSE | sedimentary | Limestone | ITP_SSE | na | na | na | 1,2 | 16.10892709 | 39.90561078 | ground | seeds seems ok, not too small |
| Italy | Pollino | 19.08.2020 | NW | sedimentary | Limestone | ITP_NW | na | na | na | 3 | 16.11491275 | 39.91050293 | ground | molt: ok, a few holes. Seeds seems empty/sterile... |

| | | | | | | | | | | | | | | |
|---|---|---|---|---|---|---|---|---|---|---|---|---|---|---|
| France, Corsica | Vizzavone | 22.08.2020 | ESE | metamorphic | Granitic | FRV_01 | 9.096328834 | 42.11532561 | 1608.426101 | 1 | 9.096328834 | 42.11532561 | branch | ok |
| France, Corsica | Vizzavone | 22.08.2020 | ESE | metamorphic | Granitic | FRV_02 | 9.096507528 | 42.11472579 | 1588.457374 | 1 | 9.096507528 | 42.11472579 | branch | ok |
| France, Corsica | Vizzavone | 22.08.2020 | ESE | metamorphic | Granitic | FRV_03 | 9.096958307 | 42.11452514 | 1569.129235 | 1 | 9.096958307 | 42.11452514 | branch | ok |
| France, Corsica | Vizzavone | 22.08.2020 | ESE | metamorphic | Granitic | FRV_04 | 9.097617207 | 42.11451165 | 1536.516633 | 1 | 9.097617207 | 42.11451165 | branch | ok |
| France, Corsica | Vizzavone | 22.08.2020 | ESE | metamorphic | Granitic | FRV_05 | 9.098088454 | 42.11426565 | 1505.985574 | 1 | 9.098088454 | 42.11426565 | branch | ok |
| France, Corsica | Vizzavone | 22.08.2020 | ESE | metamorphic | Granitic | FRV_06 | 9.098352362 | 42.11474059 | 1498.500003 | 2 | 9.098352362 | 42.11474059 | branch | ok |
| France, Corsica | Vizzavone | 22.08.2020 | ESE | metamorphic | Granitic | FRV_07 | 9.097913402 | 42.11484606 | 1529.597964 | 2 | 9.097913402 | 42.11484606 | branch | ok |
| France, Corsica | Vizzavone | 22.08.2020 | ESE | metamorphic | Granitic | FRV_08 | 9.097161127 | 42.11491338 | 1562.833116 | 2 | 9.097161127 | 42.11491338 | branch | ok |
| France, Corsica | Vizzavone | 22.08.2020 | ESE | metamorphic | Granitic | FRV_09 | 9.096698498 | 42.11503012 | 1583.614427 | 2 | 9.096698498 | 42.11503012 | branch | ok |
| France, Corsica | Vizzavone | 22.08.2020 | ESE | metamorphic | Granitic | FRV_10 | 9.096114289 | 42.11501567 | 1615.442693 | 2 | 9.096114289 | 42.11501567 | branch | ok, small seeds |
| France, Corsica | Vizzavone | 22.08.2020 | NW | metamorphic | Granitic | FRV_11 | 9.120030838 | 42.11133168 | 1258.948868 | 3 | na | na | na | no seeds |
| France, Corsica | Vizzavone | 22.08.2020 | NW | metamorphic | Granitic | FRV_12 | 9.11999376 | 42.11153211 | 1255.947621 | 3 | na | na | na | no seeds |
| France, Corsica | Vizzavone | 22.08.2020 | NW | metamorphic | Granitic | FRV_13 | 9.119749308 | 42.11164969 | 1242.009524 | 3 | na | na | na | no seeds |
| France, Corsica | Vizzavone | 22.08.2020 | NW | metamorphic | Granitic | FRV_14 | 9.119437585 | 42.11168862 | 1240.295245 | 3 | na | na | na | no seeds |
| France, Corsica | Vizzavone | 22.08.2020 | NW | metamorphic | Granitic | FRV_15 | 9.119135952 | 42.11224587 | 1218.103011 | 3 | na | na | na | no seeds |
| France, Corsica | Vizzavone | 25.09.2020 | NW | metamorphic | Granitic | FRV_16 | 9.127903 | 42.108972 | 1532 | 3 | na | na | na | no seeds |
| France, Corsica | Vizzavone | 25.09.2020 | NW | metamorphic | Granitic | FRV_17 | 9.126409 | 42.109134 | 1486 | 3 | na | na | na | no seeds |
| France, Corsica | Vizzavone | 25.09.2020 | NW | metamorphic | Granitic | FRV_18 | 9.12417 | 42.111334 | 1322 | 3 | na | na | na | no seeds |
| France, Corsica | Vizzavone | 22.08.2020 | ESE | metamorphic | Granitic | FRV_ESE | na | na | na | 1,2 | 9.096328834 | 42.11532561 | ground | ok |
| France, Corsica | Vizzavone | 22.08.2020 | NW | metamorphic | Granitic | FRV_NW | na | na | na | 3 | 9.119749308 | 42.11164969 | ground | ok |

| | | | | | | | | | | | | | | |
|---|---|---|---|---|---|---|---|---|---|---|---|---|---|---|
| France, Corsica | Coscione | 23.08.2020 | W | metamorphic | Granitic | FRC_01 | 9.163708428 | 41.88339437 | 1563.926202 | 1 | 9.163708428 | 41.88339437 | branch | ok |
| France, Corsica | Coscione | 23.08.2020 | W | metamorphic | Granitic | FRC_02 | 9.163166143 | 41.88341074 | 1547.348902 | 1 | 9.163166143 | 41.88341074 | branch | ok |
| France, Corsica | Coscione | 23.08.2020 | W | metamorphic | Granitic | FRC_03 | 9.162150352 | 41.88291837 | 1533.103186 | 1 | 9.162150352 | 41.88291837 | branch | ok, worms |
| France, Corsica | Coscione | 23.08.2020 | W | metamorphic | Granitic | FRC_04 | 9.161205038 | 41.88251363 | 1504.292604 | 1 | 9.161205038 | 41.88251363 | branch | ok |
| France, Corsica | Coscione | 23.08.2020 | W | metamorphic | Granitic | FRC_05 | 9.15959138 | 41.88236223 | 1480.515672 | 1 | 9.15959138 | 41.88236223 | branch | ok |
| France, Corsica | Coscione | 23.08.2020 | W | metamorphic | Granitic | FRC_06 | 9.160147567 | 41.8813926 | 1502.986575 | 2 | 9.160147567 | 41.8813926 | branch | ok |
| France, Corsica | Coscione | 23.08.2020 | W | metamorphic | Granitic | FRC_07 | 9.161711385 | 41.8814562 | 1507.547961 | 2 | 9.161711385 | 41.8814562 | branch | ok |
| France, Corsica | Coscione | 23.08.2020 | W | metamorphic | Granitic | FRC_08 | 9.162291957 | 41.88213963 | 1523.45474 | 2 | 9.162291957 | 41.88213963 | branch | ok |
| France, Corsica | Coscione | 23.08.2020 | W | metamorphic | Granitic | FRC_09 | 9.163306615 | 41.88224539 | 1550.295522 | 2 | 9.163306615 | 41.88224539 | branch | ok |
| France, Corsica | Coscione | 23.08.2020 | W | metamorphic | Granitic | FRC_10 | 9.163958861 | 41.88248677 | 1557.885638 | 2 | 9.163958861 | 41.88248677 | branch | ok |
| France, Corsica | Coscione | 23.08.2020 | NE | metamorphic | Granitic | FRC_11 | 9.147851232 | 41.8594289 | 1590.777198 | 3 | 9.147851232 | 41.8594289 | branch | ok |
| France, Corsica | Coscione | 23.08.2020 | NE | metamorphic | Granitic | FRC_12 | 9.148475627 | 41.86014745 | 1572.362758 | 3 | 9.148475627 | 41.86014745 | branch | ok |
| France, Corsica | Coscione | 23.08.2020 | NE | metamorphic | Granitic | FRC_13 | 9.148789799 | 41.86100352 | 1554.540869 | 3 | 9.148789799 | 41.86100352 | branch | ok |
| France, Corsica | Coscione | 23.08.2020 | NE | metamorphic | Granitic | FRC_14 | 9.149483052 | 41.86145321 | 1546.455114 | 3 | 9.149483052 | 41.86145321 | branch | 1:20, 2:21 |
| France, Corsica | Coscione | 23.08.2020 | NE | metamorphic | Granitic | FRC_15 | 9.150290186 | 41.86250466 | 1519.444455 | 3 | na | na | na | no seeds |
| France, Corsica | Coscione | 23.08.2020 | NE | metamorphic | Granitic | FRC_16_BONZAI | 9.150103402 | 41.86234633 | 1525.732756 | 3 | na | na | na | no seeds |
| France, Corsica | Coscione | 23.08.2020 | NE | metamorphic | Granitic | FRC_17 | 9.148959296 | 41.86285305 | 1538.666374 | 3 | 9.148959296 | 41.86285305 | | ok |
| France, Corsica | Coscione | 23.08.2020 | W | metamorphic | Granitic | FRC_W | na | na | na | 1,2 | 9.163708428 | 41.88339437 | ground | ok |
| France, Corsica | Coscione | 23.08.2020 | NE | metamorphic | Granitic | FRC_NE | na | na | na | 3 | 9.150290186 | 41.86250466 | ground | ok |
| France | Massane | 25.08.2020 | W | metamorphic | Granitic | FRM_01 | 3.033057225 | 42.49099419 | 749.5002085 | 1 | 3.033057225 | 42.49099419 | branch | ok, probably a few fruitful seeds, lots of sterile |

| | | | | | | | | | | | | | | |
|---|---|---|---|---|---|---|---|---|---|---|---|---|---|---|
| France | Massane | 25.08.2020 | W | metamorphic | Granitic | FRM_02 | 3.032626318 | 42.49135526 | 739.8556256 | 1 | 3.032626318 | 42.49135526 | branch | ok |
| France | Massane | 25.08.2020 | W | metamorphic | Granitic | FRM_03 | 3.032250691 | 42.49163773 | 725.9732119 | 1 | 3.032250691 | 42.49163773 | branch | ok |
| France | Massane | 25.08.2020 | W | metamorphic | Granitic | FRM_04 | 3.031798357 | 42.49196744 | 709.5962851 | 1 | 3.031798357 | 42.49196744 | branch | ok |
| France | Massane | 25.08.2020 | W | metamorphic | Granitic | FRM_05 | 3.03104821 | 42.49207094 | 686.7737505 | 1 | 3.03104821 | 42.49207094 | branch | ok, small seeds, sterile ? |
| France | Massane | 25.08.2020 | W | metamorphic | Granitic | FRM_06 | 3.030726322 | 42.49155225 | 687.2459704 | 2 | 3.030726322 | 42.49155225 | branch | ok, small seeds |
| France | Massane | 25.08.2020 | W | metamorphic | Granitic | FRM_07 | 3.03132578 | 42.49141249 | 708.2248961 | 2 | 3.03132578 | 42.49141249 | branch | ok, big seeds |
| France | Massane | 25.08.2020 | W | metamorphic | Granitic | FRM_08 | 3.031786119 | 42.49130619 | 721.3994377 | 2 | 3.031786119 | 42.49130619 | branch | ok |
| France | Massane | 25.08.2020 | W | metamorphic | Granitic | FRM_09 | 3.03220431 | 42.49086752 | 737.1061813 | 2 | 3.03220431 | 42.49086752 | branch | few seeds |
| France | Massane | 25.08.2020 | W | metamorphic | Granitic | FRM_10 | 3.032624127 | 42.49068069 | 753.6441117 | 2 | 3.032624127 | 42.49068069 | branch | ok |
| France | Massane | 25.08.2020 | E | metamorphic | Granitic | FRM_11 | 3 | 42.482778 | 1036 | 3 | 3 | 42.482778 | branch | |
| France | Massane | 25.08.2020 | E | metamorphic | Granitic | FRM_12 | 3.001111 | 42.481944 | 1022 | 3 | 3.001111 | 42.481944 | branch | |
| France | Massane | 25.08.2020 | E | metamorphic | Granitic | FRM_13 | 3.0025 | 42.48 | 969 | 3 | 3.0025 | 42.48 | branch | |
| France | Massane | 25.08.2020 | E | metamorphic | Granitic | FRM_14 | 3.007222 | 42.478333 | 853 | 3 | 3.007222 | 42.478333 | branch | |
| France | Massane | 25.08.2020 | E | metamorphic | Granitic | FRM_15 | 3 | 42.476944 | 1135 | 3 | 3 | 42.476944 | branch | |
| France | Massane | 25.08.2020 | W | metamorphic | Granitic | FRM_W | na | na | na | 3 | 3.033057225 | 42.49099419 | ground | |
| France | Massane | 25.08.2020 | E | metamorphic | Granitic | FRM_E | na | na | na | 1,2 | 3 | 42.482778 | ground | 765 |
| **Belgium** | **Sonian Forest** | **02.09.2020** | **flat** | **sedimentary** | **loess** | **BES_01** | **4.422123583** | **50.7501374** | **170.10455** | **1** | **4.422123583** | **50.7501374** | **branch** | |
| **Belgium** | **Sonian Forest** | **02.09.2020** | **flat** | **sedimentary** | **loess** | **BES_02** | **4.422256538** | **50.75038771** | **175.29383** | **1** | **4.422256538** | **50.75038771** | **branch** | |
| Belgium | Sonian Forest | 02.09.2020 | flat | sedimentary | loess | BES_03 | 4.422530964 | 50.75070214 | 169.1378242 | 1 | 4.422530964 | 50.75070214 | branch | |
| Belgium | Sonian Forest | 02.09.2020 | flat | sedimentary | loess | BES_04 | 4.422941646 | 50.75089137 | 162.4417067 | 1 | na | na | branch | |

| | | | | | | | | | | | | | | |
|---|---|---|---|---|---|---|---|---|---|---|---|---|---|---|
| Belgium | Sonian Forest | 02.09.2020 | flat | sedimentary | loess | BES_05 | 4.423024729 | 50.75112158 | 172.1650822 | 1 | 4.423024729 | 50.75112158 | branch | |
| Belgium | Sonian Forest | 02.09.2020 | flat | sedimentary | loess | BES_06 | 4.423573676 | 50.75105209 | 162.1296136 | 2 | 4.423573676 | 50.75105209 | branch | |
| Belgium | Sonian Forest | 02.09.2020 | flat | sedimentary | loess | BES_07 | 4.423491177 | 50.75071713 | 166.3464125 | 2 | 4.423491177 | 50.75071713 | branch | |
| **Belgium** | **Sonian Forest** | **02.09.2020** | **flat** | **sedimentary** | **loess** | **BES_08** | **4.423215882** | **50.75041341** | **167.08023** | **2** | **4.423215882** | **50.75041341** | **branch** | |
| Belgium | Sonian Forest | 02.09.2020 | flat | sedimentary | loess | BES_09 | 4.42273793 | 50.7501941 | 166.463497 | 2 | 4.42273793 | 50.7501941 | branch | |
| Belgium | Sonian Forest | 02.09.2020 | flat | sedimentary | loess | BES_10 | 4.422420639 | 50.74993303 | 169.0199269 | 2 | 4.422420639 | 50.74993303 | branch | |
| **Belgium** | **Sonian Forest** | **02.09.2020** | **flat** | **sedimentary** | **loess** | **BES_grd** | **na** | **na** | **na** | **1,2** | **4.422123583** | **50.7501374** | **ground** | 609 |
| Sweden | Malmö | 04.09.2020 | flat | metamorphic | caledonian nappes | SEB_01 | 13.20930751 | 55.56091927 | 91.76296316 | 1 | na | na | branch | no seeds |
| Sweden | Malmö | 04.09.2020 | flat | metamorphic | caledonian nappes | SEB_02 | 13.20865943 | 55.56060229 | 95.44023491 | 1 | na | na | branch | no seeds |
| Sweden | Malmö | 04.09.2020 | flat | metamorphic | caledonian nappes | SEB_03 | 13.20833359 | 55.56020516 | 88.82494699 | 1 | na | na | branch | no seeds |
| Sweden | Malmö | 04.09.2020 | flat | metamorphic | caledonian nappes | SEB_04 | 13.20795147 | 55.55993736 | 94.63894446 | 1 | na | na | branch | no seeds |
| Sweden | Malmö | 04.09.2020 | flat | metamorphic | caledonian nappes | SEB_05 | 13.20774352 | 55.55964482 | 86.57670204 | 1 | na | na | branch | no seeds |
| Sweden | Malmö | 04.09.2020 | flat | metamorphic | caledonian nappes | SEB_06 | 13.20818602 | 55.55942178 | 94.72793124 | 2 | na | na | branch | no seeds |
| Sweden | Malmö | 04.09.2020 | flat | metamorphic | caledonian nappes | SEB_07 | 13.20859552 | 55.5598027 | 94.72141959 | 2 | na | na | branch | no seeds |
| Sweden | Malmö | 04.09.2020 | flat | metamorphic | caledonian nappes | SEB_08 | 13.20906383 | 55.55996646 | 93.72183188 | 2 | na | na | branch | no seeds |
| Sweden | Malmö | 04.09.2020 | flat | metamorphic | caledonian nappes | SEB_09 | 13.20939628 | 55.5603182 | 88.19539188 | 2 | 13.20939628 | 55.5603182 | branch | no seeds |
| Sweden | Malmö | 04.09.2020 | flat | metamorphic | caledonian nappes | SEB_10 | 13.20997208 | 55.56070473 | 92.43263259 | 2 | na | na | branch | no seeds |
| Poland | Kadzidlowo | 07.09.2020 | flat | sedimentary | East-european platform | PLM_01 | 21.47170551 | 53.71405171 | 169.8009657 | 1 | na | na | branch | no seeds |
| Poland | Kadzidlowo | 07.09.2020 | flat | sedimentary | East-european platform | PLM_02 | 21.4718094 | 53.71367398 | 172.7805885 | 1 | na | na | branch | no seeds |
| Poland | Kadzidlowo | 07.09.2020 | flat | sedimentary | East-european platform | PLM_03 | 21.4717324 | 53.7133508 | 171.7477877 | 1 | na | na | branch | no seeds |

| | | | | | | | | | | | | | | |
|---|---|---|---|---|---|---|---|---|---|---|---|---|---|---|
| Poland | Kadzidlowo | 07.09.2020 | flat | sedimentary | East-european platform | PLM_04 | 21.47176808 | 53.71300627 | 169.7461168 | 1 | na | na | branch | no seeds |
| Poland | Kadzidlowo | 07.09.2020 | flat | sedimentary | East-european platform | PLM_05 | 21.47201855 | 53.71285478 | 165.9057071 | 1 | na | na | branch | no seeds |
| Poland | Kadzidlowo | 07.09.2020 | flat | sedimentary | East-european platform | PLM_06 | 21.47097659 | 53.71277368 | 167.3369344 | 2 | na | na | branch | no seeds |
| Poland | Kadzidlowo | 07.09.2020 | flat | sedimentary | East-european platform | PLM_07 | 21.47127103 | 53.7130549 | 169.8159037 | 2 | na | na | branch | no seeds |
| Poland | Kadzidlowo | 07.09.2020 | flat | sedimentary | East-european platform | PLM_08 | 21.47111585 | 53.7133799 | 168.3473073 | 2 | na | na | branch | no seeds |
| Poland | Kadzidlowo | 07.09.2020 | flat | sedimentary | East-european platform | PLM_09 | 21.47086902 | 53.71368987 | 169.5236763 | 2 | na | na | branch | no seeds |
| Poland | Kadzidlowo | 07.09.2020 | flat | sedimentary | East-european platform | PLM_10 | 21.4709102 | 53.71399885 | 169.345465 | 2 | na | na | branch | no seeds |
| **Poland** | **Wisla** | **09.09.2020** | **SSE** | **sedimentary** | **Carpathians flysch** | **PLW_01** | **18.85985191** | **49.67003094** | **695.75275** | **1** | **18.85985191** | **49.67003094** | **branch** | |
| Poland | Wisla | 09.09.2020 | SSE | sedimentary | Carpathians flysch | PLW_02 | 18.86027615 | 49.66978107 | 675.0584766 | 1 | 18.86027615 | 49.66978107 | branch | |
| Poland | Wisla | 09.09.2020 | SSE | sedimentary | Carpathians flysch | PLW_03 | 18.86062352 | 49.66955655 | 661.3067069 | 1 | 18.86062352 | 49.66955655 | branch | big seeds |
| Poland | Wisla | 09.09.2020 | SSE | sedimentary | Carpathians flysch | PLW_04 | 18.86104255 | 49.66941559 | 651.0725499 | 1 | 18.86104255 | 49.66941559 | branch | |
| Poland | Wisla | 09.09.2020 | SSE | sedimentary | Carpathians flysch | PLW_05 | 18.86170572 | 49.6692049 | 632.5416134 | 1 | 18.86170572 | 49.6692049 | branch | |
| **Poland** | **Wisla** | **09.09.2020** | **SSE** | **sedimentary** | **Carpathians flysch** | **PLW_06** | **18.86127057** | **49.66882205** | **627.35656** | **2** | **18.86127057** | **49.66882205** | **branch** | |
| Poland | Wisla | 09.09.2020 | SSE | sedimentary | Carpathians flysch | PLW_07 | 18.86051421 | 49.66910247 | 650.1195651 | 2 | 18.86051421 | 49.66910247 | branch | |
| Poland | Wisla | 09.09.2020 | SSE | sedimentary | Carpathians flysch | PLW_08 | 18.85992867 | 49.66917952 | 668.8233877 | 2 | 18.85992867 | 49.66917952 | branch | |
| Poland | Wisla | 09.09.2020 | SSE | sedimentary | Carpathians flysch | PLW_09 | 18.85970732 | 49.66949034 | 677.1596989 | 2 | 18.85970732 | 49.66949034 | branch | |
| **Poland** | **Wisla** | **09.09.2020** | **SSE** | **sedimentary** | **Carpathians flysch** | **PLW_10** | **18.85941897** | **49.66975466** | **682.61658** | **2** | **18.85941897** | **49.66975466** | **branch** | |
| Poland | Wisla | 09.09.2020 | NW | sedimentary | Carpathians flysch | PLW_11 | 18.85905431 | 49.67275023 | 725.5251821 | 3 | na | na | na | |
| Poland | Wisla | 09.09.2020 | NW | sedimentary | Carpathians flysch | PLW_12 | 18.85878324 | 49.67294866 | 714.053684 | 3 | na | na | na | |
| Poland | Wisla | 09.09.2020 | NW | sedimentary | Carpathians flysch | PLW_13 | 18.85858576 | 49.67331891 | 691.7239361 | 3 | na | na | na | |

| | | | | | | | | | | | | | | |
|---|---|---|---|---|---|---|---|---|---|---|---|---|---|---|
| Poland | Wisla | 09.09.2020 | NW | sedimentary | Carpathians flysch | PLW_14 | 18.85822875 | 49.67354343 | 679.1010348 | 3 | na | na | na | |
| Poland | Wisla | 09.09.2020 | NW | sedimentary | Carpathians flysch | PLW_15 | 18.8579211 | 49.67371805 | 667.7390798 | 3 | na | na | na | |
| Poland | Wisla | 09.09.2020 | SSE | sedimentary | Carpathians flysch | PLW_SSE | na | na | na | 1,2 | 18.85985191 | 49.67003094 | ground | |
| Poland | Wisla | 09.09.2020 | NW | sedimentary | Carpathians flysch | PLW_NW | na | na | na | 3 | 18.85905431 | 49.67275023 | ground | 611 |
| Poland | Biezczadski | 11.09.2020 | SE | sedimentary | Carpathians flysch | PLB_01 | 22.47208492 | 49.23328919 | 923.48184 | 1 | 22.47208492 | 49.23328919 | branch | |
| Poland | Biezczadski | 11.09.2020 | SE | sedimentary | Carpathians flysch | PLB_02 | 22.47221463 | 49.23293161 | 912.0408089 | 1 | 22.47221463 | 49.23293161 | branch | |
| Poland | Biezczadski | 11.09.2020 | SE | sedimentary | Carpathians flysch | PLB_03 | 22.4724451 | 49.23261253 | 887.6551623 | 1 | 22.4724451 | 49.23261253 | branch | |
| **Poland** | **Biezczadski** | **11.09.2020** | **SE** | **sedimentary** | **Carpathians flysch** | **PLB_04** | **22.47283517** | **49.23225533** | **873.91864** | **1** | **22.47283517** | **49.23225533** | **branch** | |
| Poland | Biezczadski | 11.09.2020 | SE | sedimentary | Carpathians flysch | PLB_05 | 22.47301723 | 49.23192029 | 861.163327 | 1 | 22.47301723 | 49.23192029 | branch | |
| Poland | Biezczadski | 11.09.2020 | SE | sedimentary | Carpathians flysch | PLB_06 | 22.47213011 | 49.23163146 | 867.0188052 | 2 | 22.47213011 | 49.23163146 | branch | |
| Poland | Biezczadski | 11.09.2020 | SE | sedimentary | Carpathians flysch | PLB_07 | 22.47201569 | 49.23204701 | 879.5781065 | 2 | 22.47201569 | 49.23204701 | branch | |
| Poland | Biezczadski | 11.09.2020 | SE | sedimentary | Carpathians flysch | PLB_08 | 22.47188009 | 49.23241376 | 886.6440423 | 2 | 22.47188009 | 49.23241376 | branch | |
| Poland | Biezczadski | 11.09.2020 | SE | sedimentary | Carpathians flysch | PLB_09 | 22.4717562 | 49.23278079 | 911.5822176 | 2 | 22.4717562 | 49.23278079 | branch | |
| Poland | Biezczadski | 11.09.2020 | SE | sedimentary | Carpathians flysch | PLB_10 | 22.47153733 | 49.23310548 | 925.4096746 | 2 | 22.47153733 | 49.23310548 | branch | |
| Poland | Biezczadski | 11.09.2020 | NNE | sedimentary | Carpathians flysch | PLB_11 | 22.46951313 | 49.23570834 | 915.2783625 | 3 | na | na | na | |
| Poland | Biezczadski | 11.09.2020 | NNE | sedimentary | Carpathians flysch | PLB_12 | 22.46961156 | 49.23614547 | 905.9917173 | 3 | na | na | na | |
| Poland | Biezczadski | 11.09.2020 | NNE | sedimentary | Carpathians flysch | PLB_13 | 22.46956881 | 49.2364755 | 893.8050889 | 3 | na | na | na | |
| Poland | Biezczadski | 11.09.2020 | NNE | sedimentary | Carpathians flysch | PLB_14 | 22.46994738 | 49.23696244 | 884.0412758 | 3 | na | na | na | |
| Poland | Biezczadski | 11.09.2020 | NNE | sedimentary | Carpathians flysch | PLB_15 | 22.47020007 | 49.23738823 | 868.3189598 | 3 | na | na | na | |
| Poland | Biezczadski | 11.09.2020 | SE | sedimentary | Carpathians flysch | PLB_SE | na | na | na | 1,2 | 22.47208492 | 49.23328919 | ground | |

| | | | | | | | | | | | | | | |
|---|---|---|---|---|---|---|---|---|---|---|---|---|---|---|
| Poland | Biezczadski | 11.09.2020 | NNE | sedimentary | Carpathians flysch | PLB_NNE | na | na | na | 3 | 22.46951313 | 49.23570834 | ground | 675 |
| Serbia | Fruska Gora National Park | 13.09.2020 | ENE | sedimentary | mix of rocks | RSF_01 | 19.63482653 | 45.13654185 | 492.5040128 | 1 | 19.63482653 | 45.13654185 | na | |
| Serbia | Fruska Gora National Park | 13.09.2020 | ENE | sedimentary | mix of rocks | RSF_02 | 19.63538953 | 45.13657403 | 478.3534321 | 1 | 19.63538953 | 45.13657403 | na | |
| Serbia | Fruska Gora National Park | 13.09.2020 | ENE | sedimentary | mix of rocks | RSF_03 | 19.63587213 | 45.13668031 | 472.464589 | 1 | 19.63587213 | 45.13668031 | na | |
| Serbia | Fruska Gora National Park | 13.09.2020 | ENE | sedimentary | mix of rocks | RSF_04 | 19.63618841 | 45.13676005 | 468.1992674 | 1 | 19.63618841 | 45.13676005 | na | |
| Serbia | Fruska Gora National Park | 13.09.2020 | ENE | sedimentary | mix of rocks | RSF_05 | 19.63671871 | 45.13705731 | 476.1973773 | 1 | 19.63671871 | 45.13705731 | na | |
| Serbia | Fruska Gora National Park | 13.09.2020 | ENE | sedimentary | mix of rocks | RSF_06 | 19.63678304 | 45.13651052 | 461.544009 | 2 | 19.63678304 | 45.13651052 | na | |
| Serbia | Fruska Gora National Park | 13.09.2020 | ENE | sedimentary | mix of rocks | RSF_07 | 19.63648893 | 45.13639953 | 463.4207375 | 2 | 19.63648893 | 45.13639953 | na | |
| Serbia | Fruska Gora National Park | 13.09.2020 | ENE | sedimentary | mix of rocks | RSF_08 | 19.63606837 | 45.13626744 | 462.9684918 | 2 | 19.63606837 | 45.13626744 | na | |
| Serbia | Fruska Gora National Park | 13.09.2020 | ENE | sedimentary | mix of rocks | RSF_09 | 19.63559965 | 45.13621033 | 477.0605037 | 2 | 19.63559965 | 45.13621033 | na | |
| Serbia | Fruska Gora National Park | 13.09.2020 | ENE | sedimentary | mix of rocks | RSF_10 | 19.63506853 | 45.13621029 | 485.0042542 | 2 | 19.63506853 | 45.13621029 | na | |
| Serbia | Fruska Gora National Park | 13.09.2020 | WNW | sedimentary | mix of rocks | RSF_11 | 19.71057975 | 45.14930718 | 537.7608939 | 3 | na | na | na | |
| Serbia | Fruska Gora National Park | 13.09.2020 | WNW | sedimentary | mix of rocks | RSF_12 | 19.71038487 | 45.14917641 | 534.7512587 | 3 | na | na | na | |
| Serbia | Fruska Gora National Park | 13.09.2020 | WNW | sedimentary | mix of rocks | RSF_13 | 19.71000801 | 45.14915893 | 525.2410294 | 3 | na | na | na | |
| Serbia | Fruska Gora National Park | 13.09.2020 | WNW | sedimentary | mix of rocks | RSF_14 | 19.70966054 | 45.14915696 | 513.2808913 | 3 | na | na | na | |
| Serbia | Fruska Gora National Park | 13.09.2020 | WNW | sedimentary | mix of rocks | RSF_15 | 19.7092886 | 45.14862556 | 501.3966623 | 3 | na | na | na | |
| **Serbia** | **Fruska Gora National Park** | **13.09.2020** | **ENE** | **sedimentary** | **mix of rocks** | **RSF_ENE** | **na** | **na** | **na** | **1,2** | **19.63482653** | **45.13654185** | **ground** | |
| **Serbia** | **Fruska Gora National Park** | **13.09.2020** | **WNW** | **sedimentary** | **mix of rocks** | **RSF_WNW** | **na** | **na** | **na** | **3** | **19.71057975** | **45.14930718** | **ground** | 334 |
| Serbia | Boranja (Krupanj) | 14.09.2020 | SSW | igneous | mix of rocks, most probably grandiorite | RSB_01 | 19.29131561 | 44.34339648 | 949.1395217 | 1 | 19.29131561 | 44.34339648 | branch | |

| | | | | | | | | | | | | | | |
|---|---|---|---|---|---|---|---|---|---|---|---|---|---|---|
| Serbia | Boranja (Krupanj) | 14.09. 2020 | SSW | igneous | mix of rocks, most probably grandiorite | RSB_02 | 19.29151693 | 44.34310184 | 943.5992437 | 1 | 19.29151693 | 44.34310184 | na | |
| Serbia | Boranja (Krupanj) | 14.09. 2020 | SSW | igneous | mix of rocks, most probably grandiorite | RSB_03 | 19.2919704 | 44.34263983 | 935.1040147 | 1 | 19.2919704 | 44.34263983 | na | |
| Serbia | Boranja (Krupanj) | 14.09. 2020 | SSW | igneous | mix of rocks, most probably grandiorite | RSB_04 | 19.29287248 | 44.34229797 | 931.3270378 | 1 | 19.29287248 | 44.34229797 | na | |
| Serbia | Boranja (Krupanj) | 14.09. 2020 | SSW | igneous | mix of rocks, most probably grandiorite | RSB_05 | 19.29303964 | 44.34124457 | 917.1317398 | 1 | 19.29303964 | 44.34124457 | na | |
| Serbia | Boranja (Krupanj) | 14.09. 2020 | N | igneous | mix of rocks, most probably grandiorite | RSB_11 | 19.28911324 | 44.3456453 | 929.1945682 | 3 | na | na | na | |
| Serbia | Boranja (Krupanj) | 14.09. 2020 | N | igneous | mix of rocks, most probably grandiorite | RSB_12 | 19.28958764 | 44.34586939 | 910.8325276 | 3 | na | na | na | |
| Serbia | Boranja (Krupanj) | 14.09. 2020 | N | igneous | mix of rocks, most probably grandiorite | RSB_13 | 19.2899141 | 44.34616297 | 906.0842672 | 3 | na | na | na | |
| Serbia | Boranja (Krupanj) | 14.09. 2020 | N | igneous | mix of rocks, most probably grandiorite | RSB_14 | 19.29019987 | 44.3464266 | 894.8894403 | 3 | na | na | na | |
| Serbia | Boranja (Krupanj) | 14.09. 2020 | N | igneous | mix of rocks, most probably grandiorite | RSB_15 | 19.29061211 | 44.34654285 | 896.1397471 | 3 | na | na | na | |
| Serbia | Boranja (Krupanj) | 14.09. 2020 | SSW | igneous | mix of rocks, most probably grandiorite | RSB_SSW | na | na | na | 1,2 | 19.29131561 | 44.34339648 | ground | |
| Serbia | Boranja (Krupanj) | 14.09. 2020 | N | igneous | mix of rocks, most probably grandiorite | RSB_N | na | na | na | 3 | 19.28911324 | 44.3456453 | ground | 235 |
| **Croatia** | **Packlenica National Park** | **16.09. 2020** | **SSW** | **sedimentary** | **Limestone** | **HRP_01** | **15.46785701** | **44.36057508** | **948.39624** | **1** | **15.46785701** | **44.36057508** | **branch** | |
| Croatia | Packlenica National Park | 16.09. 2020 | SSW | sedimentary | Limestone | HRP_02 | 15.46754478 | 44.36044549 | 930.3754062 | 1 | 15.46754478 | 44.36044549 | branch | |
| Croatia | Packlenica National Park | 16.09. 2020 | SSW | sedimentary | Limestone | HRP_03 | 15.46732508 | 44.36030579 | 920.499139 | 1 | 15.46732508 | 44.36030579 | branch | |
| Croatia | Packlenica National Park | 16.09. 2020 | SSW | sedimentary | Limestone | HRP_04 | 15.4669794 | 44.36008582 | 907.337508 | 1 | na | na | na | |

| | | | | | | | | | | | | | | |
|---|---|---|---|---|---|---|---|---|---|---|---|---|---|---|
| Croatia | Packlenica National Park | 16.09.2020 | SSW | sedimentary | Limestone | HRP_05 | 15.46697699 | 44.35979473 | 896.082493 | 1 | 15.46697699 | 44.35979473 | branch | |
| **Croatia** | **Packlenica National Park** | **16.09.2020** | **SSW** | **sedimentary** | **Limestone** | **HRP_06** | **15.46636723** | **44.36003712** | **906.6371** | **2** | **15.46636723** | **44.36003712** | **branch** | |
| Croatia | Packlenica National Park | 16.09.2020 | SSW | sedimentary | Limestone | HRP_07 | 15.46665544 | 44.36027456 | 915.3190982 | 2 | 15.46665544 | 44.36027456 | branch | small seeds |
| Croatia | Packlenica National Park | 16.09.2020 | SSW | sedimentary | Limestone | HRP_08 | 15.46698037 | 44.36042796 | 920.7731053 | 2 | 15.46698037 | 44.36042796 | branch | |
| **Croatia** | **Packlenica National Park** | **16.09.2020** | **SSW** | **sedimentary** | **Limestone** | **HRP_09** | **15.46720064** | **44.36064567** | **935.44614** | **2** | **15.46720064** | **44.36064567** | **branch** | |
| Croatia | Packlenica National Park | 16.09.2020 | SSW | sedimentary | Limestone | HRP_10 | 15.46752463 | 44.36080653 | 954.1537552 | 2 | 15.46752463 | 44.36080653 | branch | |
| Croatia | Packlenica National Park | 16.09.2020 | NO | sedimentary | Limestone | HRP_11 | 15.451667 | 44.361667 | 1131 | 3 | na | na | na | |
| Croatia | Packlenica National Park | 16.09.2020 | NO | sedimentary | Limestone | HRP_12 | na | na | na | 3 | na | na | na | |
| Croatia | Packlenica National Park | 16.09.2020 | NO | sedimentary | Limestone | HRP_13 | na | na | na | 3 | na | na | na | |
| Croatia | Packlenica National Park | 16.09.2020 | NO | sedimentary | Limestone | HRP_14 | na | na | na | 3 | na | na | na | |
| Croatia | Packlenica National Park | 16.09.2020 | NO | sedimentary | Limestone | HRP_15 | 15.446389 | 44.359444 | 1042 | 3 | na | na | na | |
| **Croatia** | **Packlenica National Park** | **16.09.2020** | **SSW** | **sedimentary** | **Limestone** | **HRP_SSW** | **na** | **na** | **na** | **1,2** | **15.46785701** | **44.36057508** | **ground** | 1:254 ; 2:190, 3 : 142 |
| **Croatia** | **Packlenica National Park** | **16.09.2020** | **NW** | **sedimentary** | **Limestone** | **HRP_NW** | **na** | **na** | **na** | **3** | **15.451667** | **44.361667** | **ground** | 1:79, 2 : 61, 3:98 |
| **Slovenia** | **Pragozd Krokar** | **17/09/2020** | **SSW** | **metamorphic** | **granitic** | **SLK_01** | **14.7638186** | **45.54328222** | **1204.9473** | **1** | **14.7638186** | **45.54328222** | **branch** | |
| Slovenia | Pragozd Krokar | 17/09/2020 | SSW | metamorphic | granitic | SLK_02 | 14.76345172 | 45.54307761 | 1194.752256 | 1 | 14.76345172 | 45.54307761 | branch | |
| **Slovenia** | **Pragozd Krokar** | **17/09/2020** | **SSW** | **metamorphic** | **granitic** | **SLK_03** | **14.76314259** | **45.54285462** | **1176.6923** | **1** | **14.76314259** | **45.54285462** | **branch** | |
| **Slovenia** | **Pragozd Krokar** | **17/09/2020** | **SSW** | **metamorphic** | **granitic** | **SLK_04** | **14.76272354** | **45.54261057** | **1163.8018** | **1** | **14.76272354** | **45.54261057** | **branch** | |
| Slovenia | Pragozd Krokar | 17/09/2020 | SSW | metamorphic | granitic | SLK_05 | 14.76250257 | 45.5423958 | 1152.771051 | 1 | na | na | na | |
| Slovenia | Pragozd Krokar | 17/09/2020 | SSW | metamorphic | granitic | SLK_06 | 14.76289874 | 45.54169118 | 1144.881295 | 2 | na | na | na | |
| Slovenia | Pragozd Krokar | 17/09/2020 | SSW | metamorphic | granitic | SLK_07 | 14.76318769 | 45.54217249 | 1162.451409 | 2 | 14.76318769 | 45.54217249 | branch | |

| | | | | | | | | | | | | | | |
|---|---|---|---|---|---|---|---|---|---|---|---|---|---|---|
| Slovenia | Pragozd Krokar | 17/09/2020 | SSW | metamorphic | granitic | SLK_08 | 14.76350777 | 45.54251845 | 1178.743179 | 2 | na | na | na | |
| Slovenia | Pragozd Krokar | 17/09/2020 | SSW | metamorphic | granitic | SLK_09 | 14.76392273 | 45.54275884 | 1195.08684 | 2 | na | na | na | |
| Slovenia | Pragozd Krokar | 17/09/2020 | SSW | metamorphic | granitic | SLK_10 | 14.76421608 | 45.54296045 | 1204.448937 | 2 | na | na | na | |
| Slovenia | Pragozd Krokar | 17/09/2020 | N | metamorphic | granitic | SLK_11 | 14.76048824 | 45.54062226 | 1183.793233 | 3 | na | na | na | |
| Slovenia | Pragozd Krokar | 17/09/2020 | N | metamorphic | granitic | SLK_12 | 14.75990027 | 45.54061439 | 1172.914656 | 3 | na | na | na | |
| Slovenia | Pragozd Krokar | 17/09/2020 | N | metamorphic | granitic | SLK_13 | 14.75938615 | 45.54070179 | 1156.551952 | 3 | 14.75938615 | 45.54070179 | branch | |
| Slovenia | Pragozd Krokar | 17/09/2020 | N | metamorphic | granitic | SLK_14 | 14.75889624 | 45.54080173 | 1149.410774 | 3 | na | na | na | |
| Slovenia | Pragozd Krokar | 17/09/2020 | N | metamorphic | granitic | SLK_15 | 14.75835862 | 45.54108536 | 1133.023242 | 3 | na | na | na | |
| **Slovenia** | **Pragozd Krokar** | **17/09/2020** | **SSW** | **metamorphic** | **granitic** | **SLK_SSW** | **na** | **na** | **na** | **1,2** | **14.7638186** | **45.54328222** | **ground** | |
| **Slovenia** | **Pragozd Krokar** | **17/09/2020** | **N** | **metamorphic** | **granitic** | **SLK_N** | **na** | **na** | **na** | **3** | **14.76048824** | **45.54062226** | **ground** | 1:33, 2:66 |
| **France** | **Marseille, Sainte Beaume** | **19.09.2020** | **N** | **sedimentary** | **Limestone** | **FRB_01** | **5.769793877** | **43.32763295** | **983.49288** | **1** | **5.769793877** | **43.32763295** | **branch** | |
| **France** | **Marseille, Sainte Beaume** | **19.09.2020** | **N** | **sedimentary** | **Limestone** | **FRB_02** | **5.769730773** | **43.32783486** | **970.38706** | **1** | **5.769730773** | **43.32783486** | **branch** | |
| France | Marseille, Sainte Beaume | 19.09.2020 | N | sedimentary | Limestone | FRB_03 | 5.76970889 | 43.32812121 | 959.1279055 | 1 | 5.76970889 | 43.32812121 | branch | |
| France | Marseille, Sainte Beaume | 19.09.2020 | N | sedimentary | Limestone | FRB_04 | 5.76935447 | 43.3287071 | 925.8814279 | 1 | 5.76935447 | 43.3287071 | branch | |
| France | Marseille, Sainte Beaume | 19.09.2020 | N | sedimentary | Limestone | FRB_05 | 5.769559332 | 43.3290381 | 912.5796585 | 1 | 5.769559332 | 43.3290381 | branch | |
| France | Marseille, Sainte Beaume | 19.09.2020 | N | sedimentary | Limestone | FRB_06 | 5.768960865 | 43.32896242 | 916.6167834 | 2 | 5.768960865 | 43.32896242 | na | |
| France | Marseille, Sainte Beaume | 19.09.2020 | N | sedimentary | Limestone | FRB_07 | 5.769144038 | 43.32845835 | 929.8066721 | 2 | 5.769144038 | 43.32845835 | na | |
| France | Marseille, Sainte Beaume | 19.09.2020 | N | sedimentary | Limestone | FRB_08 | 5.769314344 | 43.32819553 | 951.7905967 | 2 | 5.769314344 | 43.32819553 | branch | |
| France | Marseille, Sainte Beaume | 19.09.2020 | N | sedimentary | Limestone | FRB_09 | 5.769430038 | 43.32792498 | 963.8012571 | 2 | 5.769430038 | 43.32792498 | branch | |
| **France** | **Marseille, Sainte Beaume** | **19.09.2020** | **N** | **sedimentary** | **Limestone** | **FRB_10** | **5.769419644** | **43.32764624** | **987.39628** | **2** | **5.769419644** | **43.32764624** | **branch** | |

| | | | | | | | | | | | | | | |
|---|---|---|---|---|---|---|---|---|---|---|---|---|---|---|
| **France** | **Marseille, Sainte Beaume** | **19.09.2020** | **N** | **sedimentary** | **Limestone** | **FRB_N** | **na** | **na** | **na** | **1,2** | **5.769793877** | **43.32763295** | **ground** | 1:377, 2:234 |
| Spain | Zaragoza, Moncayo | 23.09.2020 | ENE | metamorphic | granitic | ESM_01 | -1.810833752 | 41.79030042 | 1498.674982 | 1 | na | na | na | |
| Spain | Zaragoza, Moncayo | 23.09.2020 | ENE | metamorphic | granitic | ESM_02 | -1.810502515 | 41.79063551 | 1483.259334 | 1 | -1.810502515 | 41.79063551 | branch | |
| Spain | Zaragoza, Moncayo | 23.09.2020 | ENE | metamorphic | granitic | ESM_03 | -1.810316124 | 41.79090823 | 1471.16076 | 1 | -1.810316124 | 41.79090823 | branch | |
| Spain | Zaragoza, Moncayo | 23.09.2020 | ENE | metamorphic | granitic | ESM_04 | -1.810153304 | 41.79123054 | 1466.750086 | 1 | na | na | na | |
| Spain | Zaragoza, Moncayo | 23.09.2020 | ENE | metamorphic | granitic | ESM_05 | -1.809920186 | 41.79167497 | 1454.589409 | 1 | na | na | na | |
| Spain | Zaragoza, Moncayo | 23.09.2020 | ENE | metamorphic | granitic | ESM_06 | -1.809421828 | 41.79152357 | 1457.26309 | 2 | na | na | na | |
| Spain | Zaragoza, Moncayo | 23.09.2020 | ENE | metamorphic | granitic | ESM_07 | -1.809748843 | 41.79116093 | 1473.758964 | 2 | -1.809748843 | 41.79116093 | branch | |
| Spain | Zaragoza, Moncayo | 23.09.2020 | ENE | metamorphic | granitic | ESM_08 | -1.810001618 | 41.7908955 | 1484.465986 | 2 | na | na | na | |
| Spain | Zaragoza, Moncayo | 23.09.2020 | ENE | metamorphic | granitic | ESM_09 | -1.810114784 | 41.79056531 | 1494.405285 | 2 | na | na | na | |
| Spain | Zaragoza, Moncayo | 23.09.2020 | ENE | metamorphic | granitic | ESM_10 | -1.810424107 | 41.7901764 | 1503.711673 | 2 | na | na | na | |
| Spain | Zaragoza, Moncayo | 23.09.2020 | NNW | metamorphic | granitic | ESM_11 | -1.8348 | 41.796517 | 1746 | 3 | -1.8348 | 41.796517 | branch | |
| Spain | Zaragoza, Moncayo | 23.09.2020 | NNW | metamorphic | granitic | ESM_12 | -1.8363 | 41.797083 | 1734 | 3 | -1.8363 | 41.797083 | branch | |
| Spain | Zaragoza, Moncayo | 23.09.2020 | NNW | metamorphic | granitic | ESM_13 | -1.835833 | 41.796767 | 1740 | 3 | na | na | na | |

| | | | | | | | | | | | | | | |
|---|---|---|---|---|---|---|---|---|---|---|---|---|---|---|
| Spain | Zaragoza, Moncayo | 23.09. 2020 | NN W | metamorphic | granitic | ESM_14 | -1.83776 7 | 41.7968 83 | 1764 | 3 | -1.83776 7 | 41.7968 83 | branch | |
| **Spain** | **Zaragoza, Moncayo** | **23.09. 2020** | **NN W** | **metamorphic** | **granitic** | **ESM_15** | **-1.83027 8** | **41.7966 67** | **1758** | **3** | **-1.83027 8** | **41.7966 67** | **branch** | |
| Spain | Zaragoza, Moncayo | 23.09. 2020 | NN W | metamorphic | granitic | ESM_16 | -1.82333 3 | 41.795 | 1634 | 3 | -1.82333 3 | 41.795 | branch | seeds with white dots |
| Spain | Zaragoza, Moncayo | 23.09. 2020 | EN E | metamorphic | granitic | ESM_E NE | na | na | na | 1,2 | -1.81083 3752 | 41.7903 0042 | ground | 1:300 ; 2:270 |
| Spain | Zaragoza, Moncayo | 23.09. 2020 | NN W | metamorphic | granitic | | na | na | na | 3 | na | na | na | no supp seeds |
| Spain | Aragon, San Juan de la Pena | 24.09. 2020 | WN W | sedimentary | conglomerate, limestones ? | ESP_01 | -0.67597 2056 | 42.5075 783 | 1163.20 0842 | 1 | -0.67597 2056 | 42.5075 783 | branch | |
| Spain | Aragon, San Juan de la Pena | 24.09. 2020 | WN W | sedimentary | conglomerate, limestones ? | ESP_02 | -0.67632 5877 | 42.5099 1354 | 1141.55 5816 | 1 | -0.67632 5877 | 42.5099 1354 | na | |
| Spain | Aragon, San Juan de la Pena | 24.09. 2020 | WN W | sedimentary | conglomerate, limestones ? | ESP_03 | -0.67722 2095 | 42.5103 4443 | 1142.47 2687 | 1 | -0.67722 2095 | 42.5103 4443 | na | |
| **Spain** | **Aragon, San Juan de la Pena** | **24.09. 2020** | **WN W** | **sedimentary** | **conglomerate, limestones ?** | **ESP_04** | **-0.67070 4151** | **42.5096 5094** | **1216.62 39** | **1** | **-0.67070 4151** | **42.5096 5094** | **branch** | |
| **Spain** | **Aragon, San Juan de la Pena** | **24.09. 2020** | **WN W** | **sedimentary** | **conglomerate, limestones ?** | **ESP_05** | **-0.67128 6765** | **42.5098 0474** | **1203.42 68** | **1** | **-0.67128 6765** | **42.5098 0474** | **branch** | |
| Spain | Aragon, San Juan de la Pena | 24.09. 2020 | WN W | sedimentary | conglomerate, limestones ? | ESP_06 | -0.67188 5207 | 42.5096 2977 | 1200.95 8768 | 1 | -0.67188 5207 | 42.5096 2977 | na | |
| Spain | Aragon, San Juan de la Pena | 24.09. 2020 | WN W | sedimentary | conglomerate, limestones ? | ESP_07 | -0.67150 1791 | 42.5096 3039 | 1204.58 954 | 1 | -0.67150 1791 | 42.5096 3039 | branch | |
| **Spain** | **Aragon, San Juan de la Pena** | **24.09. 2020** | **WN W** | **sedimentary** | **conglomerate, limestones ?** | **ESP_08** | **-0.67277 8** | **42.5091 67** | **1129** | **1** | **-0.67277 8** | **42.5091 67** | **branch** | |
| **Spain** | **Aragon, San Juan de la Pena** | **24.09. 2020** | **WN W** | **sedimentary** | **conglomerate, limestones ?** | **ESP_W NW** | **na** | **na** | **na** | **1** | **-0.67597 2056** | **42.5075 783** | **ground** | 1:175 ; 2:225 ;3:54 |

| | | | | | | | | | | | | | | |
|---|---|---|---|---|---|---|---|---|---|---|---|---|---|---|
| Switzerland | Lägern | 07.08.2020 | S | sedimentary | limestone | CHL_01 | na | na | na | 1 | na | na | branch | |
| Switzerland | Lägern | 07.08.2020 | S | sedimentary | limestone | CHL_02 | na | na | na | 1 | na | na | branch | |
| Switzerland | Lägern | 07.08.2020 | S | sedimentary | limestone | CHL_03 | na | na | na | 1 | na | na | branch | |
| Switzerland | Lägern | 07.08.2020 | S | sedimentary | limestone | CHL_04 | na | na | na | 1 | na | na | branch | |
| Switzerland | Lägern | 07.08.2020 | S | sedimentary | limestone | CHL_05 | na | na | na | 1 | na | na | na | |
| Switzerland | Lägern | 07.08.2020 | S | sedimentary | limestone | CHL_06 | na | na | na | 2 | na | na | branch | |
| Switzerland | Lägern | 07.08.2020 | S | sedimentary | limestone | CHL_07 | na | na | na | 2 | na | na | branch | |
| Switzerland | Lägern | 07.08.2020 | S | sedimentary | limestone | CHL_08 | na | na | na | 2 | na | na | branch | |
| Switzerland | Lägern | 07.08.2020 | S | sedimentary | limestone | CHL_09 | na | na | na | 2 | na | na | na | |
| Switzerland | Lägern | 07.08.2020 | S | sedimentary | limestone | CHL_10 | na | na | na | 2 | na | na | branch | |
| **Switzerland** | **Lägern** | **09.10.2020** | **S** | **sedimentary** | **limestone** | **CHL_S** | **na** | **na** | **na** | **1,2** | **na** | **na** | **ground** | 319 + 267 |
| Bosnia | Lopare | 01.09.2020 | S | na | na | HBL_01_S | 18.80783333 | 44.6931111 | 353 | na | 18.80783333 | 44.6931111 | na | |
| Bosnia | Lopare | 01.09.2020 | S | na | na | HBL_02_S | 18.80783333 | 44.6931111 | 353 | na | 18.80783333 | 44.6931111 | na | |
| Bosnia | Lopare | 01.09.2020 | S | na | na | HBL_03_S | 18.80783333 | 44.6931111 | 353 | na | 18.80783333 | 44.6931111 | na | |
| Bosnia | Lopare | 01.09.2020 | S | na | na | HBL_04_S | 18.80783333 | 44.6931111 | 353 | na | 18.80783333 | 44.6931111 | na | |
| Bosnia | Lopare | 01.09.2020 | N | na | na | HBL_01_N | 18.80783333 | 44.6931111 | 353 | na | 18.80783333 | 44.6931111 | na | |
| Bosnia | Lopare | 01.09.2020 | N | na | na | HBL_02_N | 18.80783333 | 44.6931111 | 353 | na | 18.80783333 | 44.6931111 | na | |
| Bosnia | Lopare | 01.09.2020 | N | na | na | HBL_03_N | 18.80783333 | 44.6931111 | 353 | na | 18.80783333 | 44.6931111 | na | |
| Bosnia | Lopare | 01.09.2020 | N | na | na | HBL_04_N | 18.80783333 | 44.6931111 | 353 | na | 18.80783333 | 44.6931111 | na | |
| Bosnia | Lopare | 01.09.2020 | N | na | na | HBL_05_N | 18.80783333 | 44.6931111 | 353 | na | 18.80783333 | 44.6931111 | na | |

| | | | | | | | | | | | | | |
|---|---|---|---|---|---|---|---|---|---|---|---|---|---|
| Bosnia | Lopare | 01.09.2020 | N | na | na | HBL_06_N | 18.80783333 | 44.6931111 | 353 | na | 18.80783333 | 44.6931111 | na |
| Bosnia | Lopare | 01.09.2020 | N | na | na | HBL_N | 18.80783333 | 44.6931111 | 353 | na | 18.80783333 | 44.6931111 | ground |
| Bosnia | Lopare | 01.09.2020 | S | na | na | HBL_S | 18.80783333 | 44.6931111 | 353 | na | 18.80783333 | 44.6931111 | ground |
| Bosnia | Vlasenica | 01.09.2020 | S | na | na | HBV_01_S | 18.91944444 | 44.1683333 | 1050 | na | 18.91944444 | 44.1683333 | na |
| Bosnia | Vlasenica | 01.09.2020 | S | na | na | HBV_02_S | 18.91944444 | 44.1683333 | 1050 | na | 18.91944444 | 44.1683333 | na |
| Bosnia | Vlasenica | 01.09.2020 | S | na | na | HBV_03_S | 18.91944444 | 44.1683333 | 1050 | na | 18.91944444 | 44.1683333 | na |
| Bosnia | Vlasenica | 01.09.2020 | S | na | na | HBV_04_S | 18.91944444 | 44.1683333 | 1050 | na | 18.91944444 | 44.1683333 | na |
| Bosnia | Vlasenica | 01.09.2020 | S | na | na | HBV_05_S | 18.91944444 | 44.1683333 | 1050 | na | 18.91944444 | 44.1683333 | na |
| Bosnia | Vlasenica | 01.09.2020 | S | na | na | HBV_06_S | 18.91944444 | 44.1683333 | 1050 | na | 18.91944444 | 44.1683333 | na |
| Bosnia | Vlasenica | 01.09.2020 | N | na | na | HBV_01_N | 18.91944444 | 44.1683333 | 1050 | na | 18.91944444 | 44.1683333 | na |
| Bosnia | Vlasenica | 01.09.2020 | N | na | na | HBV_02_N | 18.91944444 | 44.1683333 | 1050 | na | 18.91944444 | 44.1683333 | na |
| Bosnia | Vlasenica | 01.09.2020 | N | na | na | HBV_03_N | 18.91944444 | 44.1683333 | 1050 | na | 18.91944444 | 44.1683333 | na |
| Bosnia | Vlasenica | 01.09.2020 | N | na | na | HBV_04_N | 18.91944444 | 44.1683333 | 1050 | na | 18.91944444 | 44.1683333 | na |
| Bosnia | Vlasenica | 01.09.2020 | N | na | na | HBV_05_N | 18.91944444 | 44.1683333 | 1050 | na | 18.91944444 | 44.1683333 | na |
| Bosnia | Vlasenica | 01.09.2020 | N | na | na | HBV_06_N | 18.91944444 | 44.1683333 | 1050 | na | 18.91944444 | 44.1683333 | na |
| **Bosnia** | **Vlasenica** | **01.09.2020** | **S** | **na** | **na** | **HBV_S** | **18.91944444** | **44.1683333** | **1050** | **na** | **18.91944444** | **44.1683333** | **ground** |
| **Bosnia** | **Vlasenica** | **01.09.2020** | **N** | **na** | **na** | **HBV_N** | **18.91944444** | **44.1683333** | **1050** | **na** | **18.91944444** | **44.1683333** | **ground** |
| **Romania** | **Codrul Secular Sinca** | **01.10.2020** | | **na** | **na** | **ROS_01** | **na** | **na** | **na** | **na** | **na** | **na** | **ground** |
| Romania | Codrul Secular Sinca | 01.10.2020 | | na | na | ROS_02 | na | na | na | na | na | na | ground |
| Romania | Codrul Secular Sinca | 01.10.2020 | | na | na | ROS_03 | na | na | na | na | na | na | ground |

| | | | | | | | | | | | | | |
|---|---|---|---|---|---|---|---|---|---|---|---|---|---|
| Romania | Codrul Secular Sinca | 01.10.2020 | na | na | ROS_04 | na | na | na | na | na | na | ground | |
| Romania | Codrul Secular Sinca | 01.10.2020 | na | na | ROS_05 | na | na | na | na | na | na | ground | |
| **Romania** | **Codrul Secular Sinca** | **01.10.2020** | **na** | **na** | **ROS_06** | **na** | **na** | **na** | **na** | **na** | **na** | **ground** | |
| Romania | Codrul Secular Sinca | 01.10.2020 | na | na | ROS_07 | na | na | na | na | na | na | ground | |
| Romania | Codrul Secular Sinca | 01.10.2020 | na | na | ROS_08 | na | na | na | na | na | na | ground | |
| Romania | Codrul Secular Sinca | 01.10.2020 | na | na | ROS_09 | na | na | na | na | na | na | ground | |
| Romania | Codrul Secular Sinca | 01.10.2020 | na | na | ROS_10 | na | na | na | na | na | na | ground | |
| Romania | Codrul Secular Sinca | 01.10.2020 | na | na | ROS_11 | na | na | na | na | na | na | ground | total seeds romania: 346 |
